# Comprehensive annotation of olfactory and gustatory receptor genes and transposable elements revealed their evolutionary dynamics in aphids

**DOI:** 10.1101/2025.04.14.648604

**Authors:** Sergio Gabriel Olvera-Vazquez, Xilong Chen, Aurélie Mesnil, Camille Meslin, Fabricio Almeida-Silva, Johann Confais, Yann Bourgeois, Célia Lougmani, Karine Alix, Nicolas Francillonne, Nathalie Choisne, Stephane Cauet, Jean-Christophe Simon, Christelle Buchard, Nathalie Rodde, David Ogereau, Claire Mottet, Alexandre Degrave, Elorri Segura, Benoît Barrès, Emmanuelle Jacquin-Joly, William Marande, Dominique Lavenier, Fabrice Legeai, Amandine Cornille

## Abstract

Understanding the molecular evolution of genes involved in parasite adaptation and the role of transposable elements (TEs) in driving their diversification is key to unraveling how populations adapt to their environments. In phytophagous insects like aphids, olfactory (OR) and gustatory receptor (GR) genes are crucial for host recognition, yet their post-duplication evolution remains insufficiently explored. Here, we analyzed 521 OR and 399 GR genes, alongside TEs, across 12 aphid genomes with varying host ranges. Aphid lineages with broader host ranges exhibited higher evolutionary rates, driven by gene family expansions linked to host interaction, including lipid metabolism, immune function, and transposase activity. The evolution of OR and GR genes post-duplication was shaped by diversifying selection, with bursts of positive selection followed by long periods of purifying selection, consistent with adaptation to new hosts. OR and GR genes originated from proximal and tandem duplications, with younger TE activity enriched near these genes compared to other genomic regions, suggesting a role for TEs in catalyzing tandem duplications and fueling diversification. The star-like topology of the OR phylogenetic tree, low synteny, and recent TE activity around OR genes support a faster evolutionary rate for ORs than GRs - a trend observed in other insect taxa. This study provides insights into molecular mechanisms underlying host adaptation in aphids and presents the first high-quality genome assembly of *Dysaphis plantaginea*, a major apple pest, with a comprehensive annotation of chemosensory genes and TEs. These resources offer a foundation for research on aphid genome evolution, insect–plant interactions.

## Introduction

Studying the genomic bases of parasite adaptation to their host can contribute to understanding the evolutionary processes underlying population adaptation (Gladieux et al. 2014; Simon et al. 2015; Moller and Stukenbrock 2017) and help tackle pressing issues such as parasite outbreaks (Kirk et al. 2013; Cacciò et al. 2018; Desvignes et al. 2022; Gowler et al. 2023). The questions related to parasite adaptation to their host entail identifying the number and nature of genes involved and their mode of evolution by drift (Barton and Mallet 1996; White et al. 2021; Lynch 2023) or natural selection (Fay and Wittkopp 2008; Karlsson et al. 2014; Arnab et al. 2023). Recently, the role of repetitive regions of the genomes - called transposable elements (TEs hereafter) - in driving genomic variation at loci, fueling the arms race between parasites and their hosts, has also gained attention (Duplessis et al. 2011; Hartmann et al. 2017; Plissonneau et al. 2018; Rocafort et al. 2022; O’Donnell et al. 2024).

Gene duplication is a source of genomic variation that can fuel parasite adaptation to its host (Wagner 1998; Magadum et al. 2013). When a newly duplicated gene is divergent, it can be immediately favored by selection if the host cannot recognize it, improving the parasite’s fitness. The freshly duplicated gene is then fixed by positive selection (Kondrashov et al. 2002; Kondrashov 2012). Periods during which genes evolve under positive selection that may occur recurrently or episodically (also called diversifying selection) and be followed by prolonged episodes of purifying selection (leading to an average rate of sequence evolution, *d*_N_/*d_S_*<1 (Kimura 1977; Kimura 1980)) on a long-term evolutionary time scale. Alternatively, a newly duplicated gene can have redundant functions immediately, and one of the copies could accumulate deleterious mutations and become a pseudogene (Assis and Bachtrog 2013). The role of positive selection and purifying selection in driving the molecular radiation of genes involved in the parasite adaptation to their host remains to be explored beyond model systems such as viruses, bacteria, fungal pathogens, or insect models such as drosophila (McBride 2007; Assis and Bachtrog 2013; McKenzie and Kronauer 2018; Remick et al. 2023; Lehman et al. 2024).

Studying the TE content and localization in parasite genomes is a way to understand the evolutionary dynamics of genes involved in the parasite adaptation to their hosts. The selfish proliferation of TEs that drives their insertion dynamics can also drive the evolutionary dynamics of duplicated genes involved in the parasite adaptation to their host (Gilbert et al. 2021; Oggenfuss et al. 2023). TEs are supposed to be catalyzers of proximal duplicated gene copies, i.e., genes near each other but separated by several genes (10 or fewer genes) (Liu and Wessler 2017; Dupeyron et al. 2019). In the TE-rich genomes of some plant pathogens, genome structure alternating gene-rich and TE-rich regions are observed and suggested to facilitate the generation of variation that fuels the arms race between parasites and their hosts (Seidl and Thomma 2017; Frantzeskakis et al. 2019; Frantzeskakis et al. 2019; Schrader and Schmitz 2019; Torres et al. 2020). While the role of TEs is well documented in plant pathogens, their role in shaping the diversity of genes involved in insect-plant interactions is much less characterized (Stergiopoulos and De Wit 2009; Simon et al. 2015; Torres et al. 2020; Seong and Krasileva 2023). Yet, insects have a massive variation in TE and chemosensory gene content (Elliott and Gregory 2015; Gilbert et al. 2021).

Several chemosensory gene families play a key role in host recognition in herbivorous insects. Olfactory receptors (ORs) and gustatory receptors (GRs) are crucial signal-transducing receptors in insect olfaction and gustation. Other chemosensory genes, such as insect-binding proteins (IBPs) and odorant-binding proteins (OBPs), assist in transporting odorants (Pelosi et al. 2006). Since ORs and GRs trigger neural responses, they are essential targets for evolutionary studies. Additionally, functional data on ORs and GRs have shown their connection to speciation, making them central to understanding insect adaptation (Jacquin-Joly and Merlin 2004; Almeida et al. 2014; Engsontia et al. 2014). OR and GR duplicated genes form highly diversified families whose rapid evolution is driven by gene duplication (Niimura and Nei 2007; Kulmuni et al. 2013). Phylogenetic analyses reveal the origin of OR and GR. GRs emerged early in metazoan evolution and are the oldest chemosensory gene family (Robertson 2019). In arthropods, OR receptor genes likely originated from the diversification of the ancestral gustatory receptor (GR-like) family, as revealed by phylogenetic analyses (Robertson 2015; Eyun et al. 2017; Robertson 2019). However, the fate of OR and GR genes post-duplication is less documented and shows mixed patterns across insect groups (Hallem et al. 2006; McBride 2007; Joseph and Carlson 2015). In lepidopterans, gene gains and occasional losses appear linked to host adaptation, with polyphagous species such as *Helicoverpa armigera* and *Spodoptera frugiperda* showing GR expansions associated with broader host ranges (Xu et al. 2016; Gouin et al. 2017; Pearce et al. 2017). Evidence of positive selection in these genes suggests host-driven adaptation, as seen in the *Spodoptera* species (Meslin et al. 2022). Hemipterans, such as the pea aphid, show rapid OR and GR gene expansions alongside positive selection, supporting a model of adaptation by host specialization (Smadja et al. 2009). Conversely, in dipterans like *Drosophila sp.*, OR/GR diversity seems to be shaped more by genetic drift than selection, influenced by species endemism and smaller effective populations (Gardiner et al. 2008; Hansson and Stensmyr 2011). Therefore, research on the evolution of OR and GR genes still lags behind a few insect model species and is limited primarily due to technical challenges. High sequence divergence across and within species complicates the automated annotation of OR and GR genes, necessitating manual annotation and extensive curation to ensure accuracy.

The relationship between TEs and chemosensory gene evolution also remains an open question. TEs are prevalent across insect genomes and vary significantly in content and type among species (McCullers and Steiniger 2017; Gilbert et al. 2021). In some insects, high densities of TEs are linked to duplicated genes tied to adaptive traits, suggesting that TEs may promote gene duplications that support evolutionary responses to environmental pressures (Gilbert et al. 2021). For example, in the ant *Cardiocondyla obscurior*, OR genes are located in TE-dense regions, or “TE islands”, implying that TEs might facilitate OR gene duplication (McKenzie and Kronauer 2018). Similarly, TEs are found near insecticide-resistance genes in the pest *Helicoverpa armigera*, suggesting that TEs may be instrumental in driving adaptive gene duplications (Klai et al. 2020). However, a detailed analysis of TE density and diversity surrounding GR and OR genes beyond a few model insects is still needed to fully understand the relationship between TEs and OR/GR evolution. TE-mediated tandem duplications may significantly shape the evolution of OR and GR gene families, contributing to their adaptation to diverse host plants.

Due to their diverse host range, aphids are ideal for studying the genomic basis of host adaptation (Peccoud et al. 2010; Simon et al. 2015; Shih et al. 2023). OR and GR genes are key receptors for aphids, which use chemicals to screen their environmental landscape and detect their host (Robertson 2019; Robertson et al. 2019). The sequencing of several genomes of aphid species and the annotations of OR and GR genes in two aphid models, *Acyrthosiphon pisum* Harris and *Myzus persicae* Sulzer provided the first hints on the diversity and evolution of chemosensory genes and TE content in aphids (Smadja et al. 2009; The International Aphid Genomics Consortium 2010; Thorpe et al. 2018; Julca et al. 2020; Mathers et al. 2020; Biello et al. 2021; Byrne et al. 2022; Zhang, S., et al. 2022; He et al. 2023). The role of TEs in the evolution of aphids has been investigated in only a few studies (Bouallègue et al. 2017; Ahmad et al. 2021). Comparative studies on the evolutionary dynamics of olfactory receptor (OR) and gustatory receptor (GR) genes across diverse aphid species with different host plants remain scarce. Additionally, the evolutionary dynamics of TEs in aphid genomes, particularly near OR and GR genes, are poorly understood.

This study investigates the evolutionary forces shaping gene repertoire evolution in 12 aphid species with varying host plants. We focus on the olfactory receptor (OR) and gustatory receptor (GR) multigene families, particularly in the Aphidinae subfamily. Notably, the Aphidinae subfamily, which includes aphids with complex life cycles that colonize diverse host plants, provides a valuable comparative framework for understanding the evolutionary trajectories of OR and GR gene families during divergence. Additionally, we analyze TE diversity and dynamics to understand their role in OR and GR evolution. Using a newly assembled genome of the rosy apple aphid (*Dysaphis plantaginea* Passerini) and 11 publicly available aphid genomes from both the Aphidinae and Eriosomatinae subfamilies—along with one aphid-like outgroup species—we systematically annotated OR and GR genes and classified TEs using standardized pipelines. We address four key questions: (1) Do aphid species in the Aphidinae subfamily show expansion of the OR and GR gene family, or any other gene family that could be related to host adaptation? (2) What duplication mechanisms drive the evolution of OR and GR genes? (3) What types of selective pressures act on OR and GR genes post-duplication? (4) Do TE content and dynamics vary across aphid species, and more specifically, is there TE enrichment near OR and GR genes? These questions aim to elucidate the genomic bases driving aphid speciation and adaptation to their hosts.

## Results

### Annotation of GR and OR genes in the aphid and aphid-like genomes

We retrieved 11 high-quality genome assemblies from the Aphidinae and Eriosomatinae subfamilies, along with one aphid-like species, *Daktulosphaira vitifoliae* Fitch (Phyloxerinae subfamily) (BUSCO completed gene values > 95.7%, Tables S1 and S2). Additionally, we included a newly assembled genome of *D. plantaginea* (version 3; see details in Text S1, Tables S2 and S3). We then identified the OR and GR gene repertoire in the 13 selected genome assemblies.

Through a systematic homology search and manual curation across the 13 genomes (Figure S1), we identified 668 OR and 580 GR genes. Among these, 521 OR and 399 GR genes were newly annotated in 10 aphid species (*A. gossypii*, *E. lanigerum*, *D. noxia*, *D. plantaginea*, *M. cerasi*, *M. persicae*, *P. nigronervosa*, *R. maidis*, *R. padi*, and *S. miscanthi*; Table S3). In the remaining three species (*A. glycines, A. pisum,* and *D. vitifoliae*), we confirmed 147 OR and 181 GR genes previously annotated without identifying any additional OR or GR genes. On average, we detected 44 ± 16.85 GR and 51 ± 12.78 OR genes per species (Table S1, Figure S2), and the number of OR and GR genes was significantly lower only in *D. noxia* (*P*<0.001 Model 1, Table S4, Figure S3). Although not statistically significant (*P* > 0.05, Model 1, Table S5, Figure S3), the number of OR genes was higher than the GR genes. Additionally, there was no significant correlation between genome size and OR or GR gene repertoires (Spearman’s correlation: *ρ_GR_* = 0.23, non-significant; *ρ_OR_* = −0.02, non-significant).

### Gene gains and losses, including OR and GR genes, in the Aphidinae subfamily

As a preliminary step before analyzing the evolutionary history of gene repertoires in aphids, we confirmed the genetic relationships among the studied species based on single-copy genes detected with OrthoFinder (Emms and Kelly 2019). From the 13 assemblies, OrthoFinder identified 5,868 orthogroups shared among all species, including 891 single-copy orthogroups (Figure S4 and Table S5). The phylogenetic tree constructed using these 891 single-copy orthogroups, representing 240,958 conserved amino acid residues and scaled with TimeTree, was highly statistically supported (Figure 1). The ultrametric tree provided a more comprehensive and well-supported phylogenetic representation, consistent with the known taxonomy of the species (Choi et al. 2018; Mathers et al. 2020; Byrne et al. 2022; Huang et al. 2023).

**Figure 1.**
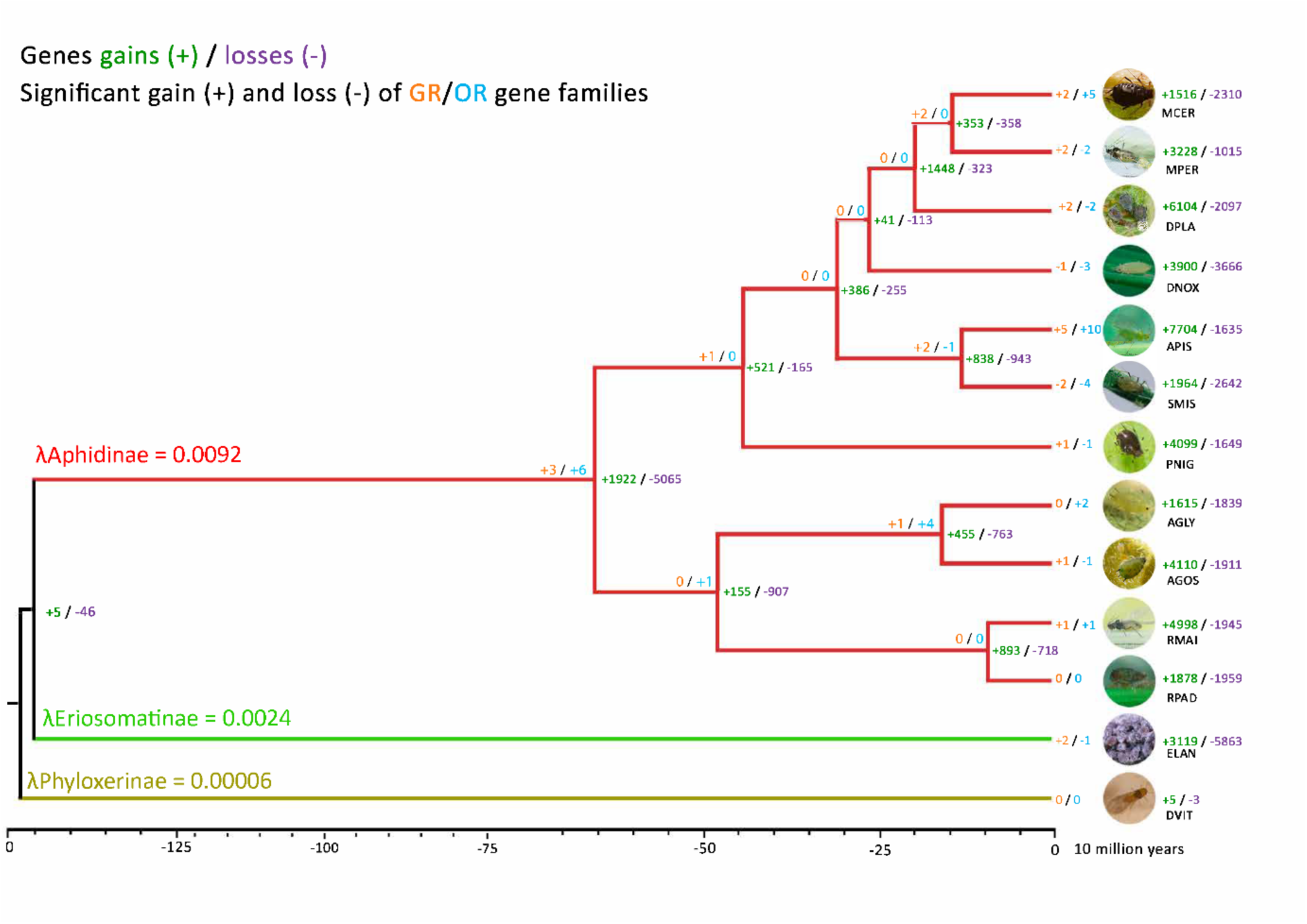
Gene family expansion and contraction events in 12 aphid species and the outgroup *Daktulosphaira vitifoliae* detected using CAFE. The internal branch with the most significant number of gene gains (expansions) and losses (contractions) corresponds to the most recent common ancestor of aphid species from the *Aphidinae* subfamily. Species abbreviations are as follows: AGLY = *Aphis glycines* clone BT1; AGOS = *Aphis gossypii*; APIS = *Acyrthosiphon pisum* clone LSR1; DNOX = *Diuraphis noxia*; DPLA = *Dysaphis plantaginea*; DVIT = *Daktulosphaira vitifoliae* (outgroup); ELAN = *Eriosoma lanigerum*; MCER = *Myzus cerasi*; MPER = *Myzus persicae* clone O; PNIG = *Pentalonia nigronervosa*; RMAI = *Rhopalosiphum maidis*; RPAD = *Rhopalosiphum padi*; SMIS = *Sitobion miscanthi*. Gene gains (+) are shown in green, and gene losses (-) in purple. Significant gains and losses of olfactory receptor (OR) and gustatory receptor (GR) genes are highlighted in orange and blue, respectively. All branches are supported by 100% bootstrap values. **●** indicates the estimated evolutionary rate (birth-and-death rate) for each branch.

Subsequently, we employed CAFE (Mendes et al. 2021) to analyze the evolutionary dynamics of gene repertoires in the 12 aphid species (excluding ***Daktulosphaira vitifoliae*** because it is a aphid-like species), including the previously detected OR and GR genes. For CAFE, we used the ultrametric tree inferred earlier and the multi-copy orthogroups identified by OrthoFinder as inputs. OrthoFinder identified 28,672 multi-copy orthogroups (Figure S2 and Table S5). Following the CAFE pipeline’s recommendations, we removed gene copies identified in more than one species or those with fewer than 100 gene copies to avoid bias in our inferences (using a customized Python script from the CAFE pipeline). As a result, we retained 22,948 orthogroups, which included 42 GR orthogroups (with 578 GR genes out of the 580 previously annotated) and 47 OR orthogroups (comprising 617 OR genes out of the 668 previously annotated) for further analyses.

We statistically analyzed the evolutionary rates (gene gain and loss) among gene families (orthogroups) across the phylogeny. We compared models assuming a single evolutionary rate along the phylogeny with models allowing evolutionary rates to vary based on taxonomic rank (genus, subfamily, family, species) or host preferences (details in Table S6). Among these, CAFE provided the most substantial support for the model with subfamily-specific evolutionary rates (log-likelihood -lnL = 243,475, ε = 0.02878, *P-value* < 0.05; Table S7). In this model, the Aphidinae subfamily exhibited the highest estimated gene gain and loss rate (birth-and-death rate, *λ* = 0.0092, P-value < 0.05, Figure 1, Table S7), which was 3.8 times greater than that of the Eriosomatinae subfamily (λ = 0.0024, P-value < 0.05, Figure 1). This result indicates that gene family turnover was higher in Aphidinae than in Eriosomatinae and Phylloxerinae (Figure 1).

We focused on 147 significantly expanded and 51 significantly contracted gene families (*P-value* < 0.01) in the Aphidinae subfamily, containing at least 50 genes, for gene ontology (GO) analysis. GO analysis of expanded gene families revealed pathways related to metabolism, transcription, lipid molecules and interactions, immune responses, and transposase activity (Figure S5, Table S8). The GO analysis of contracted gene families included general functions (Figure S6, Table S9). Notably, a large gene gain was observed in the clade of *M. persicae, M. cerasi, and D. plantaginea* clade (+1,148 genes). This clade includes species that infest fruit trees but also display a wide range of other life-history traits, including variation in the life cycle, reproductive strategies, and host specialization (see Table S6 for details).

Of the 42 GR orthogroups, 11 exhibited significant expansions, and five showed significant contractions across the 12 aphid species. Six of the 47 OR orthogroups had significant expansions and nine significant contractions. We identified four expanded chemosensory receptors (two GRs and two ORs) in the Aphidinae (Figure 1). The highest gain of GR and OR genes was observed in *A. pisum* and *M. cerasii* Fabricius, which are known for their broad host-plant range and ecological diversity (Blackman and Eastop 2008).

These findings highlight significant changes in OR and GR genes within a species subfamily known for its broad host range. However, their expansion and contraction occurred less than in other gene families associated with metabolism, transcription, lipid signaling, molecular interactions, immune responses, and transposase activity. To further understand the evolutionary dynamics of OR and GR genes, we investigated their origins and the selective pressures acting upon them to assess whether they evolved faster than the rest of the genome.

### Weak micro-synteny and evolution by tandem and proximal duplication of GR and OR genes

First, we examined the preserved order of genes on the same chromosome (Duran et al., 2009) and the arrangement between related species (also called micro-synteny). Second, we explored the mechanisms driving the evolution of OR and GR genes, focusing on gene duplication events, including proximal and tandem duplications, that contribute to the evolution of these families.

We investigated the microsynteny of OR and GR protein sequences in the 12 aphid genomes with the syntenet R package (Almeida-Silva et al. 2023). A total of 216 out of 668 ORs (33.8%) and 228 out of 580 GRs (39.5%) belonged to syntenic clusters (37 and 60 clusters, respectively) across the 12 aphid species. Syntenic clusters were, therefore, not deeply conserved across taxa (i.e., not specific species; Table S10). The GR genes were more syntenic than the OR genes (Table S10). Syntenet identified 70 and 31 genes with putative chemosensory functions as they were clustered with manually annotated GR and OR genes, some of which were clustered into orthogroups by OrthoFinder (Table S10). However, these genes were excluded from further analyses as they were not detected during the systematic manual annotation of OR and GR genes. Consequently, only those genes confirmed through manual annotation as OR/GR genes were considered for detailed analysis.

We investigated the mode of evolution of OR and GR genes with the R package doubletrouble (Almeida-Silva and Van de Peer 2025). We identified and classified them as dispersed duplicates (gene copies neither neighboring nor colinear), proximal duplicates (gene copies separated by 10 or fewer genes on the same chromosome), tandem duplicates (gene copies closely adjacent to each other in the same chromosome), and segmental duplicates (gene copy block with high sequence identity; (Qiao et al. 2019; Abdullaev et al. 2021), depending on their copy number and genomic distribution. Most OR and GR genes originated from tandem and proximal duplications (Figure 2). For some species, most genes originated from “dispersed duplication,” meaning they could not be classified into segmental, tandem, or proximal duplications. This is often because dispersed duplicates were duplicated too long ago, and subsequent genomic rearrangements have obscured their original duplication mechanisms, such as tandem duplications that were later broken apart.

**Figure 2.**
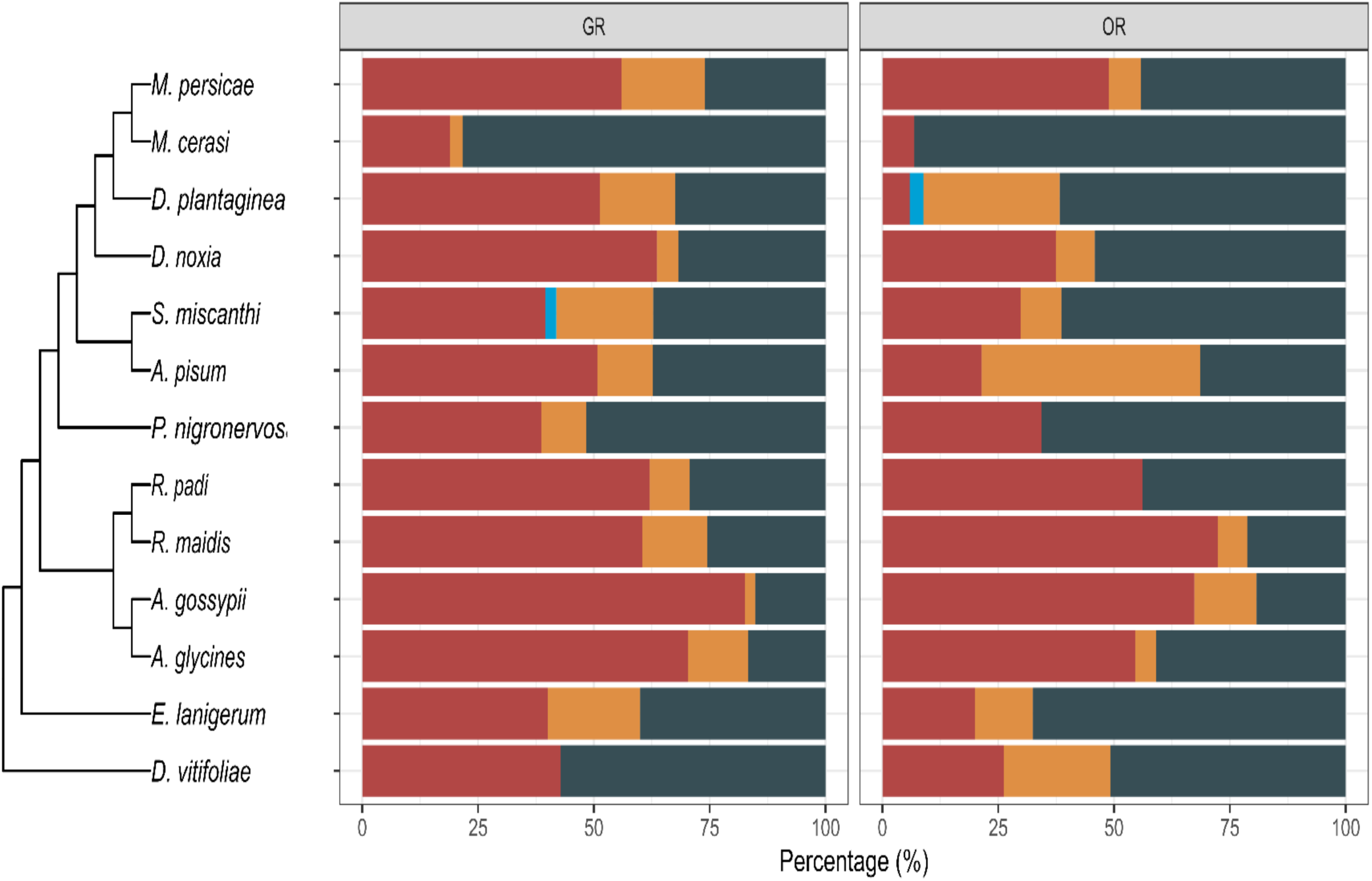
Mode of evolution of olfactory and gustatory receptor genes (OR and GR, respectively) distribution in the 12 aphid species and the grape phylloxera *D. vitifoliae*. a. Percentage of duplication modes of each aphid species inferred with R package doubletrouble. Dispersed duplication: genes cannot be classified as segmental, tandem, or proximal

Therefore, the OR and GR gene families evolved mostly by tandem duplications and are not deeply conserved across taxa, but the GR genes are more conserved than the OR. Our results suggested different evolutionary histories between OR and GR genes. We therefore further investigated selective pressures acting on OR and GR genes following tandem duplication.

### A faster evolution of OR and GR genes compared to genome-wide levels

We calculated the substitution rate of protein sequences (ω) and the ratio of nonsynonymous (dN) to synonymous (dS) substitutions for OR and GR genes, comparing them to the rest of the genome (hereafter referred to as OG for single-copy orthologous genes). This analysis aimed to determine whether OR and GR genes evolved under neutral evolution, positive selection, or purifying selection.

First, we defined the subfamilies of OR and GR to investigate the selective pressure. We classified the subfamilies of OR and GR genes, building a Maximum Likelihood tree on 365 OR amino-acid sequences and 381 GR amino-acid sequences from our set of 12 aphid and one aphid-like genomes (Figures S7 and S8, Table S11). The GR genes clustered into 12 subfamilies (monophyletic groups with high bootstrap values > 80%, Figure S8), and the OR genes clustered into 24 subfamilies (high bootstrap values > 80%, Figure S9). We detected two known GR subfamilies, the sugar and fructose receptors (GR01 and GR02, respectively, Figure S8), based on the similarities with previously reported ones in *A. pisum* and *A. glycines.* We also detected the conserved odorant co-receptor (Orco or OR01, Figure S9) subfamily based on the clustering with the already reported Orco genes of *A. pisum*, *A. glycines*, and *D. vitifoliae*. The high number of OR subfamilies relative to GR subfamilies and the star-like topology of the phylogenetic tree (Figure S9) suggested that OR genes are evolving faster than GR genes or may have diverged more recently, as previously suggested (Robertson 2015; Eyun et al. 2017; Robertson 2019). We explore these hypotheses in further detail below.

We used the codeml program in the PAML package v4 (Yang 2007) to estimate the substitution rate of the protein sequences (*ω*) of the 12 and 24 GR and OR gene subfamilies, respectively. Codeml computes the ratio (*ω*) for various models of sequence evolution (tree branch or/and codon-site). The most likely model explaining the data is chosen based on a likelihood ratio test computed among models. Here, we used the branch and site models. We did not use the branch-site model as we had no previous information about the phylogeny of these genes or an apparent chemosensory gene family clustering to test for selection. For most GR and OR subfamilies, the free ratio (different *ω* values among branches, Yang, 2007) was the most statistically supported model (Table S12), indicating that within each OR and GR subfamily, sequences evolved with their *ω* (Table S12). One of the exceptions was the fructose receptor genes (GR02), which displayed a single evolutionary rate among the sequences (Table S12). Of the 381 GR sequences analyzed, 57.7% evolved under purifying selection, 21.62% under positive selection, and 20.61% neutrally. Of the 365 OR sequences, 46.11% evolved under purifying selection, 25.47% under positive selection, and 27.34% under neutral evolution. The signatures of selection were not OR/GR subfamily-specific: each subfamily presented each of the three signatures of selection (Tables S13). However, lineage-specific models (branch models) have a limited ability to detect short episodes of positive selection that affect only a few amino acids or amino acids under recurrent diversifying selection. We, therefore, used site-specific models implemented in codeml on sequences of each GR and OR subfamily. The site-specific models can account for site rate variation and detect recent episodes of positive selection (diversifying selection). We used three pairs of site-specific models to test for recurrent, diversifying selection: M0 (one ratio) and M3 (Discrete), M1 (Neutral) and M2 (Selection), and M7 (Beta) and M8 (Beta & ω). The likelihood ratio test showed higher support for models assuming neutral or evolution by purifying selection (M0, M1a, and M7) models than the alternative ones supporting diversifying selection (M3, M2a, and M8) (Tables S14 and S15), except the gene groups GR02 (fructose receptor) and OR03 (Table S15). The posterior probability (PPs > 95%) of the Bayes empirical Bases (BEB) distribution of the inferred positively selected sites under the M8 model further indicated 19 sites in the GR02 (seven overlapping sites between M2a and M8) and one site in the OR03 (this site overlapped in M2a and M8; Table S16) under positive selection. These inferred sites should be taken cautiously because the likelihood inference method is known to falsely detect positively selected sites (Suzuki and Nei 2002; Zhang 2004). Altogether, branch and site models indicate that OR and GR genes mainly evolved under purifying selection.

We extracted the nonsynonymous-to-synonymous substitution ratio (*ω* or *dN/dS*) from the best-fit models. We compared these values between GR and OR genes and orthologous genes across the entire genome. We used 50 randomly selected single-copy gene families identified by OrthoFinder to represent the rest of the genome, as single-copy genes provide a more accurate representation of background evolutionary rates, avoiding the potential confounding effects of gene duplication and copy number variation seen in multi-copy gene families. We ran the same pipeline explained above for GR and OR genes to estimate *ω* for each OG family. Fourteen percent of the single-copy orthologous gene groups evolved under the same *ω*, while 86 % fitted the branch models with a specific *ω* (Table S17). For the site models, all OG families (except one out of the 49) evolved under the same *ω* (M0), including 72% evolving under purifying selection (*ω*<1; Tables S18 to S21). We then statistically compared OG’s *dN*/*dS* ratio (ω) to the OR and GR gene subfamilies. The OR and GR genes showed significantly weaker purifying selection than single-copy genes (Model 1, Figure S10, Table S22). Our results indicate that GR and OR genes mainly evolve under purifying selection (Figure S10), with nonsignificant differences in selection pressure on OR genes than on GRs. We also observed a negative correlation between the *dN*/*dS* ratio and divergence among species, indicating that genes with the lower divergence were under the highest evolutionary rates (*p-value*<0.0004, *r^2^*= −0.09, Figure S11).

Purifying selection was the dominant signature observed in OR and GR genes, affecting over 50% of these genes. However, these signatures are notably weaker than those in other genomic regions (OG), indicating that OR and GR genes in aphids evolve at higher rates than the rest of the genomes. Additionally, approximately 20-30% of OR and GR genes show evidence of recent positive selection, together with the observed negative correlation between the evolutionary rates of GR and OR genes and species divergence, suggesting that these genes undergo rapid episodes of positive selection, followed by sustained periods of purifying selection, which aligns with a pattern of diversifying selection.

### Variable TE contents and evolutionary dynamics among aphid species

We predicted and annotated TEs as a preliminary step to examine their enrichment near OR and GR genes (see next result part). The analysis revealed that TE prediction and annotation were not significantly correlated with genome assembly quality (see Tables S23 and S24 and Materials and Methods for details). We observed significant variation in TE content and evolutionary dynamics across aphid species. While no correlation between TE content and genome size was found (*r²*=0.02, NS, Table S3), differences in the proportion of TE types were noted, particularly Terminal Inverted Repeat (TIR) class II elements. Kimura distance-based divergence revealed varying evolutionary histories among species, with some species, such as *D. plantaginea* and *M. persicae*, showing recent TE activity, particularly in TIR elements. In contrast, others exhibited older and more stable TE dynamics. These differences suggest distinct evolutionary pressures on TE content across aphid species (Table S3, Figures 3 and S12).

**Figure 3.**
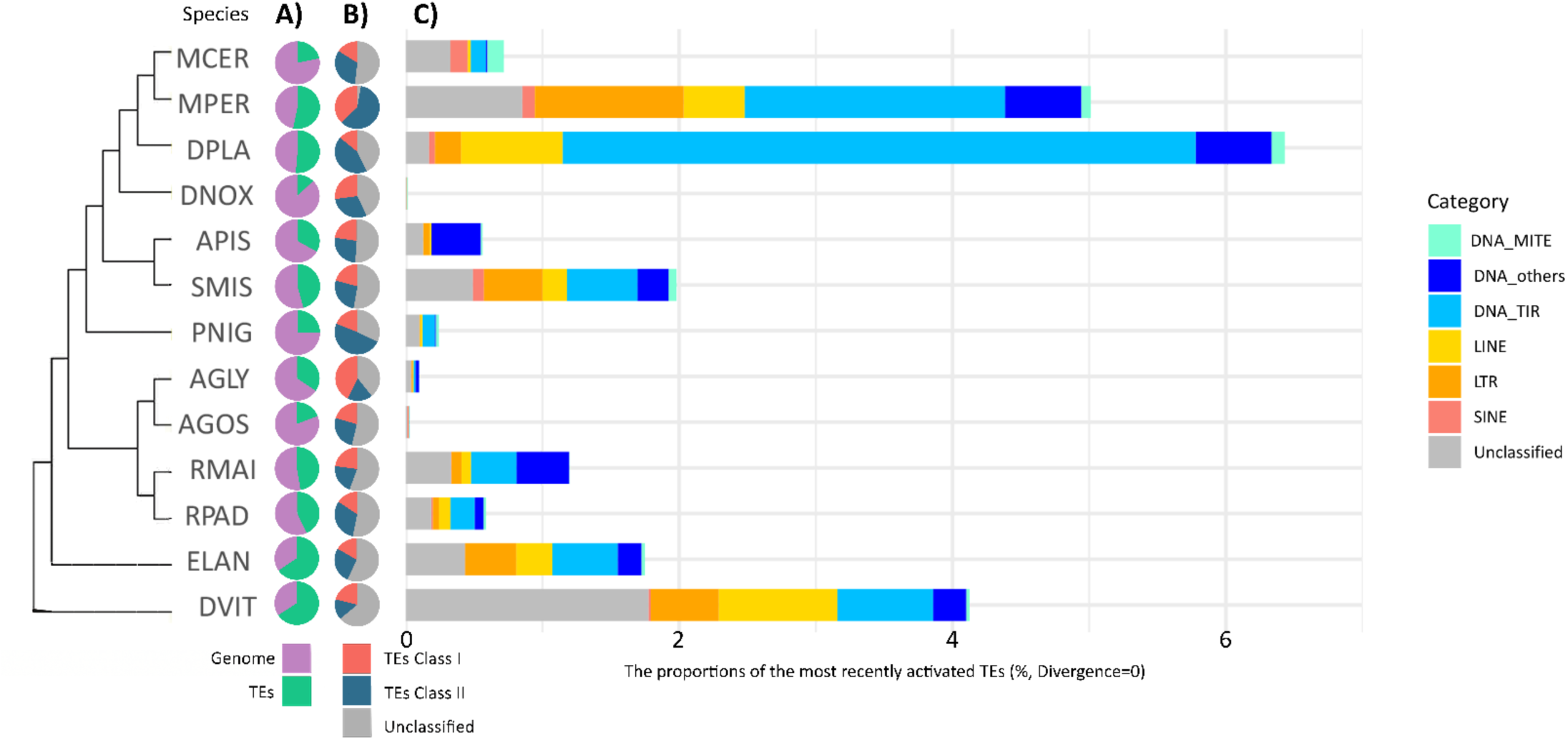
Transposable element (TE) content in the twelve aphid species and the grape phylloxera in a phylogenetic context. (IQtree, with all bootstraps > 0.99). A) Proportion of TEs in the genome for each analyzed species. The purple portion represents the fraction of the genome occupied by TEs. This proportion is based on the respective TE library for each species. B) Proportion of Class I (yellow to red) and II (green to blue) consensus TEs in each analyzed species. C) Kimura distance distribution for the most recently active TE across the analyzed species (0% divergence). The x-axis represents the Kimura distance for each TE category. “Order” is used instead of “category” for precision. The TE categories (e.g., DNA_MITE, LINE, SINE, etc.) are indicated on the x-axis. Species abbreviations: AGLY = *Aphis glycines* clone BT1; AGOS = *Aphis gossypii*; APIS = *Acyrthosiphon pisum* clone LSR1; DNOX = *Diuraphis noxia*; DPLA = *Dysaphis plantaginea*; DVIT = *Daktulosphaira vitifoliae*; ELAN = *Eriosoma lanigerum*; MCER = *Myzus cerasi*; MPER = *Myzus persicae* clone O; PNIG = *Pentalonia nigronervosa*; RMAI = *Rhopalosiphum maidis*; RPAD = *Rhopalosiphum padi*; SMIS = *Sitobion miscanthi*.

### Limited TE enrichment, but younger TEs are more frequently found near OR genes than GR genes and other regions of the genome

To investigate the role of TEs in the diversity of GR and OR genes in aphids, we assessed TE enrichment in the vicinity of these genes. TEs were detected using the LOLA tool (Sheffield and Bock 2016), and relationships between genes and TE enrichment were determined using TEgrip (Meguerditchian et al. 2021).

We observed a more significant enrichment of TEs near OR genes than GR genes (using 10 Kb or 2 Kb regions). However, not all aphid species showed TE enrichment (Fisher’s test, Bonferroni correction: q-values < 0.05, Figure 4, Tables S25). TEs significantly enriched in *D. plantaginea* and *D. vitifoliae* included DNA transposons and retrotransposons with variable relationships to genes (Table S25, Figure 3). In *D. plantaginea*, 62.5% of GR genes and 88.6% of OR genes were associated with enriched TEs, with TEs encompassing GR genes or located proximally (< 2 kb), while seldom overlapping OR genes. In *D. vitifoliae*, six distinct TEs were found near OR genes, including two sequences associated with 31.7% and 55.5% of OR genes. *R. padi* and *M. cerasi* also showed TE enrichment around OR genes (Table S26). We identified evidence of purifying selection across all OR and GR genes associated with TEs (Table S25).

**Figure 4.**
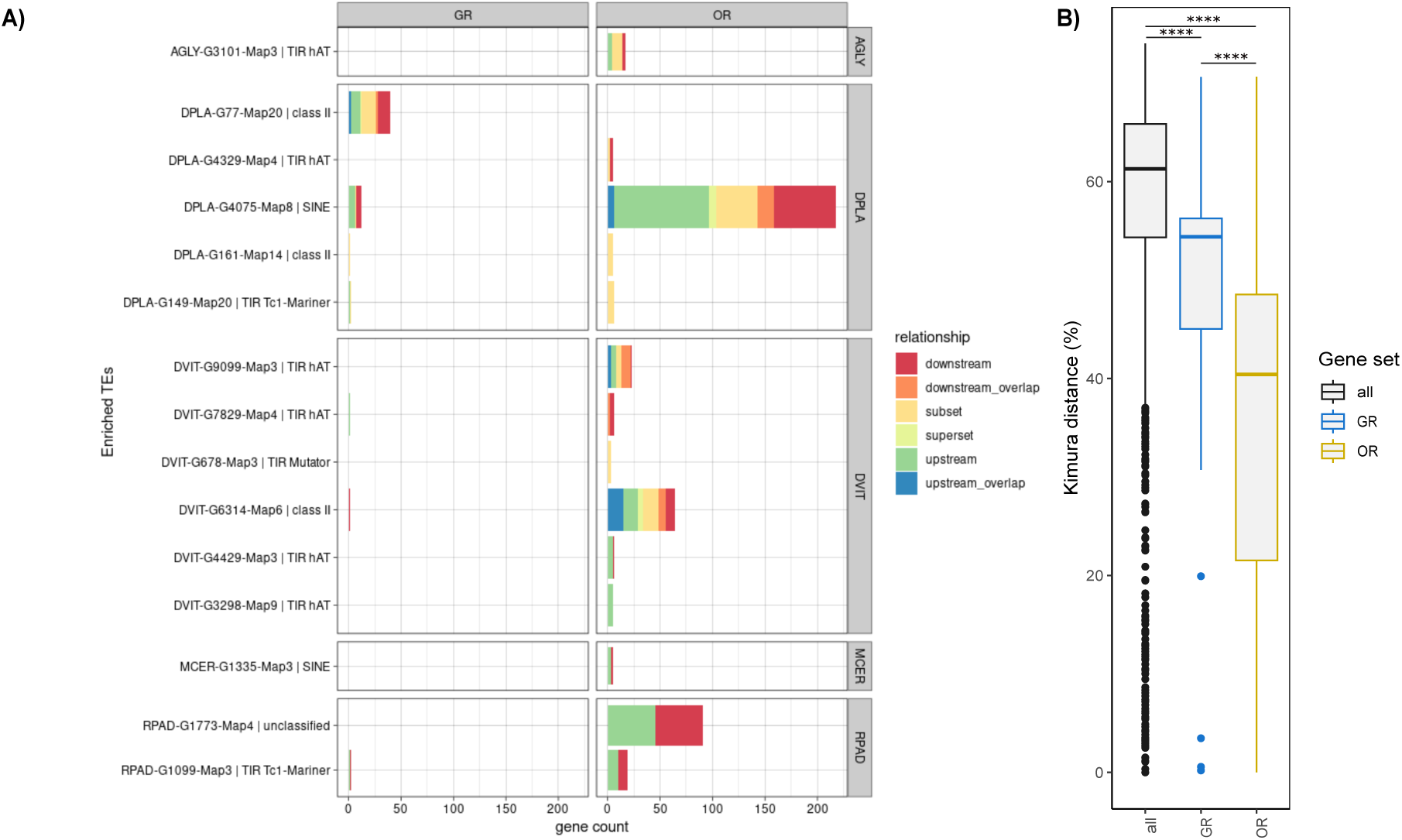
Distribution and age of transposable element (TE) enrichment near olfactory and gustatory receptor genes in 12 aphid species. (A) TE Enrichment Analysis This panel illustrates the enrichment of individual TE copies near olfactory receptor (OR) and gustatory receptor (GR) genes based on 15 annotated consensus TE sequences. The analysis identifies significant enrichment within 10 kb of OR and GR genes, determined by mapping TE copies to their genomic locations. Since consensus TE sequences represent generalized forms, they do not directly indicate enrichment but serve as a reference for identifying individual TE insertions. TE-gene relationships are categorized as follows: Subset (light orange): TE located within the gene. Superset (light yellow): Gene located within the TE. Downstream (red): TE located downstream of the gene. Upstream (light green): TE located upstream of the gene. Overlap (orange): Gene overlaps with TE upstream/downstream regions by at least one base. Among the 12 aphid species analyzed, only five species—*Aphis glycines* (AGLY), *Dysaphis plantaginea* (DPLA), *Daktulosphaira vitifoliae* (DVIT), *Myzus cerasi* (MCER), and *Rhopalosiphum padi* (RPAD)—exhibited significant TE enrichment near OR and GR genes. (B) TE Age Distribution (Kimura Distance Analysis) The distribution of Kimura distances is shown for TE copies associated with OR and GR genes compared to all TE copies in the genome. The Kimura distance measures sequence divergence, estimating TE insertion age. TE copies are categorized as follows: All TE copies (black), TE copies associated with GR genes (blue), TE copies associated with OR genes (yellow). Statistical differences between these distributions were assessed using Kruskal-Wallis tests, with significance levels indicated as ns: Not significant (p > 0.05), *p < 0.05, p < 0.01, p < 0.005, p < 0.001*. This analysis provides insights into whether TE insertions near OR and GR genes are evolutionarily recent or ancient across aphid species.

We compared the ages of TEs enriched in OR and GR genes to those in other genomic regions, using OG single-copy groups as a reference. Age estimation was based on the Kimura distance, a proxy for sequence divergence. Our analysis revealed that TEs associated with OR genes were significantly younger than those linked with GR genes or other genome regions (Kruskal-Wallis test, *P*<0.01, Figure 4). This finding suggests more recent TE activity near OR genes, potentially contributing to the recent evolution in OR genes.

OR and GR originated from proximal and tandem duplication and showed TE enrichment nearby. TEs near OR genes were younger than those near GR genes. The star-like shape of the OR phylogenetic tree, low synteny, and more recent TE dynamics near OR genes indicate a more recent evolution of OR than GR genes, as suggested in other systems (Robertson 2019).

## Discussion

Our goal was to identify the repertoire and mode of evolution of chemosensory receptor genes that enable phytophagous insects to adapt to their host plants, as well as to explore the potential role of transposable elements in driving the evolutionary dynamics of these genes. We focused on aphids, a key family of phytophagous insects, for which a comprehensive analysis of gene family evolution has not yet been conducted beyond studies focused on crop pest species. We investigated gene family evolution, focusing specifically on OR and GR genes, which are essential for host recognition in phytophagous insects, as well as the diversity and evolutionary dynamics of TEs in aphids. This comprehensive study required extensive manual annotation, covering hundreds of OR and GR genes and TEs across the genomes of 12 aphid species, including several significant crop pests. Among these, we analyzed a newly assembled, high-quality genome of *D. plantaginea*, a key pest affecting fruit trees. Our findings indicate that gene families related to aphid-plant interactions, particularly OR and GR gene families, have undergone expansion through proximal duplications in an aphid subfamily characterized by its wide host range. Additionally, the evolution of these OR and GR genes is faster than the rest of the genome and influenced by diversifying selection, characterized by rapid episodes of positive selection followed by longer periods of purifying selection, reflecting the adaptive dynamics involved in host recognition and specialization (Smadja et al. 2009; Engsontia et al. 2014; Gouin et al. 2017). The presence of younger TEs associated with OR genes, compared to GR genes and other genomic regions, along with the star-like structure of the OR phylogenetic tree and low synteny, suggests that OR genes have experienced more recent evolutionary changes. This aligns with observations in other model systems (Robertson 2019). The higher TE enrichment near OR and GR genes indicates their potential role in catalyzing tandem duplications, contributing to the diversification of these gene families. Additionally, we observed species-specific TE content and evolutionary dynamics, particularly in aphids infesting fruit trees, such as *D. plantaginea* and *M. persicae*. This study enhances our understanding of the evolutionary mechanisms that drive the adaptation of phytophagous to hosts, as well as the role of TEs in shaping genomic architecture (Smadja et al., 2009; Engsontia et al., 2014; Gouin et al., 2017; Klai et al., 2020).

### The impact of host range on chemosensory gene evolution in aphids

We found that the Aphidinae subfamily exhibited 3.8 times higher evolutionary rates compared to Eriosomatinae, driven by changes in gene families (including OR and GR gene families) associated with host interactions, including those related to lipid metabolism, immune responses, and transposase activity. The Aphidinae is the most species-rich subfamily of the Aphidinae and includes over 200 genera and various life cycles, including shifts between woody perennial primary hosts and multiple herbaceous secondary hosts (Blackman and Eastop 2000; Blackman and Eastop 2008). The higher evolutionary rates of duplicated gene families associated with plant interactions in this aphid subfamily, which exhibits a high host plant diversity, may suggest that host diversity can drive speciation in this subfamily (Peccoud et al. 2010). Previous studies indicate a positive correlation between host range and speciation rates (Weingartner et al. 2006; Hardy and Otto 2014; Nicholson et al. 2015; Robertson 2019). While OR and GR genes and other gene families exhibited duplications in some aphid species that could be linked to host diversity, further investigation is needed to determine how these expansions correlate with ecological adaptation. Complex factors, including seasonal and geographic variability, influence host breadth in aphids (Blackman and Eastop 2008). Some species are specialists, depending on a single host throughout their life cycle, whereas others are generalists, feeding on multiple hosts across different seasons (Dixon 1977). Therefore, while gene family expansions and contractions provide insights into aphid genomic evolution, additional ecological and functional analyses are required to clarify their adaptive role.

Purifying selection is the dominant signature observed in OR and GR genes, affecting over 50% of these genes. However, these signatures are notably weaker than those in other genomic regions (OG), indicating that OR and GR genes in aphids evolve at higher rates. Additionally, approximately 20-30% of OR and GR genes show evidence of recent positive selection. These results, with the observed negative correlation between the evolutionary rates of GR and OR genes and species divergence, suggest that these genes undergo rapid episodes of positive selection, followed by sustained periods of purifying selection, consistent with a pattern of diversifying selection. The interaction between host plants and herbivorous insects can create selective pressures that drive the diversification of chemosensory genes. In host-parasite systems, such as those involving aphids and their host plants, variations in these genes are essential for adapting to defensive volatile or tastant compounds produced by plants (Simon et al. 2015; Züst and Agrawal 2016; Beran and Petschenka 2022). For example, aphids with enhanced olfactory capabilities are better equipped to detect and navigate toward suitable hosts from a distance, increasing their fitness. Once on the plant, improved gustatory perception helps aphids identify appropriate feeding sites (Joseph and Carlson 2015). The faster evolutionary rate observed in OR genes compared to GR genes may reflect their primary role in long-distance host recognition, while GRs are involved at a later stage, facilitating feeding once the aphid has settled on the plant. However, a population genomic study of the selection pattern in candidate genes or through genome scans is needed to characterize the footprint of selection in aphid genomes on a short time scale (Simon et al. 2015). Such studies exist for *A. pisum* (Smadja et al. 2009; Eyres et al. 2016; Nouhaud et al. 2018), the aphid model species, but they still need to be extended to more species and populations.

Although this was not the primary objective of our study, our comprehensive annotation of OR and GR genes across several aphid species provided an opportunity to compare their repertoires among species. We did not find any variation in the size of the OR/GR gene repertoires among aphid species. As we used a standardized pipeline on high-quality reference genomes followed by manual curation, this lack of differences cannot be due to the missing of OR and GR genes. Some studies concluded that the numbers of OR and GR genes are associated with the host breadth of each aphid species (Quan et al. 2019; Robertson 2019). In other insect species, various conclusions are drawn from different host specializations, such as the specialist fig wasp *Ceratosolen some* Mayr (Xiao et al. 2013) or the specialized human body louse *Pediculus humanus* Haeckel, which has 10 ORs and 6 GRs (Kirkness et al. 2010).

Despite the variable host preferences of the studied aphid species, a higher sampling of more aphid species across various subfamilies would help to answer better whether the increase of the gene repertoires of OR and GR genes occurred associated with host breadth. Even other chemosensory genes (e.g., CPs and OBPs (Vogt 2003; Pelosi et al. 2006)) could be investigated, but additional massive manual annotation would be required. Finally, specific genes may undergo pseudogeneization following specialization in the host plants in phytophagous conditions. This adaptation may result from a reduced need for specific sensory capabilities when they specialize in particular plant types. Here, we decided to focus on non-degraded sequences; these genomic fossils or evolutionary relics may help better understand each genome’s evolution rate, as they generally evolve without selective constraints (Podlaha and Zhang 2010).

### Transposable elements and the evolution of GR and OR gene duplication in aphids

TE content and organization in aphid genomes are crucial data for understanding the evolutionary dynamics of duplicated genes that contribute to insect adaptation to their hosts. Our research revealed that OR and GR genes in aphids likely originated from tandem duplications. Similar tandem duplication has been observed in *Drosophila* species related to genes involved in defense against pathogens, insecticide resistance, chorion development, cuticular peptides, and lipases (Rogers et al. 2015).

High densities of transposons are frequently associated with duplicated genes in various insect species, indicating that TEs may facilitate the formation of proximal duplicated gene copies (Liu and Wessler 2017; Dupeyron et al. 2019; Gilbert et al. 2021). We found only a few enrichments of TEs near OR and GR genes. However, we found that TEs near OR and GR genes are younger than elsewhere in the genome, indicating heightened TE activity in these regions, particularly in species like *D. plantaginea*. The rapid turnover of TEs, characterized by high birth and death rates and the strong purifying selection, may account for the limited detection of TE enrichment (Nei et al. 2000). TEs only persist as long as active elements can move to new positions within a genome. Over time, individual transposon lineages are often lost within specific clades. Transposons may have played a significant role in catalyzing the tandem duplication of OR and GR genes, thereby contributing to the diversity observed within these gene families. For example, in the ant *Cardiocondyla obscurior*, OR genes are located in TE-rich regions, suggesting a significant role for TEs in their duplication (McKenzie and Kronauer 2018). In the moths Spodoptera spp., some TEs are significantly enriched near GR gene clusters (Meslin et al. 2022). TEs are also close to insecticide resistance genes in *Helicoverpa armigera* Hübner. However, evidence of this effect appears to be limited to genes that have recently diversified.

We detected variations in the evolutionary dynamics of TEs among different aphid species. However, it is premature to establish a direct connection between these TE dynamics and the ecological or evolutionary histories of the species involved. Further investigation of the transposon insertion landscape along the genome of aphid populations should provide a deeper understanding of the selection and the demographic forces shaping the adaptation of aphids to their host plant. Such a population genomics lens would provide insights into the role of TEs in aphid evolution and, more generally, into the balance between demography and selection in the evolution of insect (Lockton et al. 2008; Oggenfuss et al. 2021).

## Conclusion

This study provides new insights into the molecular evolution of genes involved in adaptation and diversification in phytophagous insects. Our findings indicate that OR and GR genes evolve faster than the rest of the genome following gene duplication, shaped by diversifying selection with episodic bursts of positive selection interspersed with periods of purifying selection. This pattern supports the hypothesis that shifts to novel host plants in aphids drive adaptation to new environments, eventually leading to species diversification. Additionally, our results suggest that TEs may play a role in shaping key genes associated with aphid adaptation. However, establishing a direct causal link between TE activity and the ecological or evolutionary trajectories remains premature. Future research incorporating a broader range of aphid genomes with varying host breadths will help clarify the contributions of gene duplication and TE dynamics to host plant adaptation. Comparative analyses between highly polyphagous species and host-specialized aphids could provide further insights into the relationship between host range and OR and GR gene evolution. Additionally, investigating other gene families involved in aphid-host interactions, such as digestion genes (Baril et al. 2023), other chemosensory genes (e.g., CSPs and OBPs (Vogt 2003; Pelosi et al. 2006)), and chemosensory ionotropic receptors (Rytz et al. 2013), alongside population genomic analyses and TE dynamics, will deepen our understanding of insect adaptation to their host. This study also delivers valuable genomic resources, including a high-quality genome assembly for *D. plantaginea* (the rosy apple aphid) and comprehensive annotations of ORs, GRs, and TEs. OR genes under recent positive selection present promising targets for environmentally sustainable pest control strategies. We anticipate that these genomic datasets will be essential for future investigations into the molecular mechanisms of host adaptation in aphids and other phytophagous insects.

## Material and methods

### Aphid genomic resources

We assembled a high-quality reference genome of *D. plantaginea,* the rosy apple aphid (see details in Text S1, Tables S2 and S3). We obtained the genomes of twelve aphid species and one aphid-like species from previous publications (Table S1): *Aphis glycines* (bt1), *Acyrthosiphon pisum* LSR1 (v2), *Aphis gossypii*, *Diuraphis noxia*, *Eriosoma lanigerum*, *M. persicae* (clone O), *Myzus cerasi*, *Pentalonia nigronervosa*, *Rhopalosiphum maidis*, *Rhopalosiphum padi*, and *Sitobion miscanthi*, and one aphid-like species of the Phylloxeridae family, *Daktulosphaira vitifoliae* Fitch. We selected aphid species identified as polyphagous or generalists with a broad host range to study gene evolution in a subfamily exhibiting the greatest diversity of life history traits, mainly to host breadth, when definable. The quality of the genomes was assessed using N50 (the shortest contig length needed to cover 50% of the genome) and BUSCO (Benchmarking Universal Single-Copy Orthologs) scores, as detailed in the publications (Table S1). We downloaded the genome assemblies and annotations of the aphid species from the AphidBase (Bioinformatics platform for Agroecosystem Arthropods called BIPAA (Legeai et al. 2010): https://bipaa.genouest.org/is/aphidbase/).

### Annotation of chemosensory genes

We utilized previously reported annotated OR and GR genes for *A. glycines*, *A. pisum*, and *D. vitifoliae* ((Robertson et al. 2019; Rispe et al. 2020); Table S1) as input for Exonerate (Slater and Birney 2005), the InsectOR pipeline (Karpe et al. 2021), and Scipio (Keller et al. 2008) to predict genes in both chemosensory gene families across the 13 genomes mentioned above (Figure S1). We manually curated and filtered the predicted gene models on the Apollo server (Lee et al. 2013) hosted on the BIPAA platform using BLAST on NCBI to identify similarity percentages with previously reported OR/GR gene sequences (Robertson et al., 2019) and confirmed the presence of conserved protein domains (Sayers et al. 2019). After annotating the candidate OR/GR genes for each aphid species, we initiated a second round of gene detection using the InsectOR pipeline (Karpe et al., 2021) to refine and improve the final set of OR and GR gene annotations. As a final validation, we aligned the protein sequences for each gene family using MAFFT v7.4 (Kuraku et al. 2013; Katoh et al. 2019) with the standard parameters and a gap-opening penalty of two. We removed or split sequences with extremely short lengths or highly divergent sequences compared to known protein sequences using available OR and GR sequences from *A. pisum* and *A. glycines* reported by Robertson et al. as references (2019).

We performed Spearman rank correlation tests between OR or GR repertoires and genome size estimated from the assemblies. We statistically tested for differences in numbers of OR and GR genes among aphid species using a generalized linear model (glm.nb function) as follows:

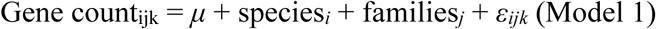

Where *Gene count_ijk_* represented the number of genes of species *i* from the OR or GR family *j*, while *μ* denoted the overall mean. Species and families were treated as fixed effects, and *ε_ijk_* was the residual term. The residual was fitted to a Negative Binomial distribution using the R package MASS, version 7.3-58.3 (Venables and Ripley 2013).

We incorporated the annotated GR and OR genes into the previously retrieved genome annotation files (GFF) for the 12 aphid species and one aphid-like species. To eliminate overlapping regions, we utilized BCFTools version 1.5 (Danecek et al. 2021). These updated files served as input for detecting orthogroups and conducting the analyses described below.

### Ortholog identification and species tree inferences

We used the combined set of predicted proteins from the 13 species to infer orthogroups using OrthoFinder v2.4. This analysis allowed us to identify conserved gene families and assess the evolutionary relationships among the species, facilitating a comprehensive understanding of gene repertoire evolution in the aphid lineages studied (Emms and Kelly 2019). We then constructed a phylogenetic tree of the 13 species using single-copy orthologous groups identified by OrthoFinder v2.4 (Emms and Kelly 2019). This approach enabled us to accurately depict the evolutionary relationships among the aphid species accurately and provided a foundation for further analyses of gene evolution. We aligned and concatenated the single-copy orthologs using MAFFT (Kuraku et al. 2013; Katoh et al. 2019), ensuring a comprehensive alignment for subsequent phylogenetic analyses. We used TrimAl (Capella-Gutiérrez et al. 2009) to remove the poorly aligned regions with a conservation threshold of 60% and a gap threshold of 90%. We constructed a maximum likelihood phylogenetic tree using IQ-TREE v2.0.6 (Minh et al. 2020) based on the aligned single-copy orthologous sequences and conducted 10,000 bootstrap replicates. The phylogenetic tree was edited with FigTree v1.4.4 (http://tree.bio.ed.ac.uk/software/figtree/), with *D. vitifoliae* serving as the outgroup.

### Expansion and contraction of gene families

We utilized the Computational Analysis of Gene Family Evolution (CAFE v5; (Mendes et al. 2021)) to identify rapidly evolving gene families, which indicate a high rate of genomic turnover (gains and losses per gene per million years). AFE estimates the birth-death parameter λ (probability that any gene will be gained or lost at each node and terminal branch, accounting for gene family expansions and contractions) along a provided phylogenetic tree scaled to time with the same length of root tips and gene family count estimated with OrthoFinder (Emms and Kelly 2019).

First, we constructed an ultrametric species tree from the IQ-TREE analysis using the r8s software (http://sourceforge.net/projects/r8s/). We used divergence times and a reference species tree created with the TimeTree website (www.timetree.org) as inputs, along with five available time references (Kim et al. 2011; Ren et al. 2013; Hardy et al. 2015).

We ran the CAFE software using the ultrametric tree and gene family counts inferred from OrthoFinder. To ensure accuracy, we employed a Python script to eliminate gene families with gene copies in only one species or that contained more than 100 gene copies in one or more species. This correction in gene counting helps to avoid potential errors in the estimation of evolutionary rates ((Mendes et al. 2021); https://github.com/hahnlab/cafe_tutorial/). We conducted the CAFE analysis using several modes to estimate the net gain and loss rates. These included 1) a single parameter (λ) across the entire phylogeny and 2) multiple parameters for each gene family. We established a single evolutionary rate (gene birth-death rate) as our null hypothesis and proposed different evolutionary rates along each branch of the phylogeny as our alternative hypothesis. The evolutionary hypotheses tested included factors such as host preference (tree versus herbaceous host), specialization (generalist vs. specialist), phylogeny, and taxonomic rank (family, subfamily, tribe, and genus). Finally, we evaluated the fit of the models by comparing the log-likelihoods of the alternative and null models, utilizing a likelihood-ratio test to identify significant differences (LRT; (Magis et al. 2010; Thissen et al. 2013)) script on RStudio v1.3.1.

We performed a Gene Ontology (GO) enrichment analysis on the 147 expanded and 51 significantly contracted gene groups identified by CAFE. We employed the TopGO R package (Alexa and Rahnenfuhrer 2010) with the “elim” method to identify genes associated with biological processes or functions based on Gene Ontology (GO) terms.

### Micro-synteny analyses and mode of evolution of GR and OR genes

Expanded gene families often occur as tandem arrays, a genomic architecture that can contribute to increased gene birth and death rates, increasing copy number variation among species (Ohno 2013). Therefore, we examined how genomic organization varies between GR and OR subfamilies in aphid species to generate insights into the molecular evolution shaping OR and GR subfamily diversity. We built a synteny network using the R/Bioconductor package Syntenet (Almeida-Silva et al. 2023). We used chemosensory protein sequences and the genomic coordinates for each species (gff files). We explored the synteny of OR and GR genes, focusing only on gene clusters containing the OR and GR genes previously annotated (Table S1). We visualized the phylogenomic profiles of the syntenic genes as a heat map. We performed the protein alignment using DIAMOND (Buchfink et al. 2015) in sensitive mode (*e-value* = 1e-10, top hits = 5). To identify and classify duplicated gene pairs, we used the doubletrouble R package (Almeida-Silva and Van de Peer 2025), applying its default parameters for detecting whole genome duplication (WGD), small-scale duplications (SSD), and paralogous genes based on the syntenic blocks.

### Selective pressures acting on OR and GR genes

We inferred selection pressures within OR and GR gene subfamilies defined from the phylogenetic trees we built. OR and GR subfamilies were monophyletic groups supported by high bootstrap values (>80%). We also visually inspected the tree topology. We then defined the foreground and background branches for the selection analysis. We detailed the methods below.

First, we built two phylogenetic trees for GR and OR genes. Fructose and Orco genes were used as outgroups (Robertson et al. 2019) to root the trees. From the protein files of each GR and OR genes, we extracted their coding sequences (CDS) from the original FASTA files of the respective genome assemblies. We aligned the sequences using MUSCLE (Edgar 2004) with default parameters and a maximum of 16 iterations. We utilized Gblocks (Castresana 2000) to curate the sequences, restricting contiguous non-conserved positions and removing any lines that introduced significant gaps in the alignment. We also removed sequences <150 pb or presenting stop codons (Figure S7 and Table S11). The sequences were then converted to codon alignments using PAL2NAL (Suyama et al. 2006). We constructed separate phylogenetic trees for the OR and GR genes using IQ-TREE v2.0 (Minh et al. 2020), employing the model finder option, ultrafast bootstrap with 1,000 replicates, and a branch support test (SH-aLRT) with 1,000 replicates. The OR and GR subfamilies were visually identified on the phylogenetic trees as monophyletic groups with bootstrap support exceeding 80%.

We used the maximum likelihood approach implemented in codon-based analysis (*codeml*) with PAML version 4.2 (Yang 2007) to assess positive and purifying selection for each GR and OR subfamily. Codeml computes the likelihood of a large set of molecular evolution models that estimate *ω,* the ratio of the non-synonymous mutation rate (*dN*) over the synonymous mutation rate (*dS*) (Kimura 1980). The most likely model of molecular evolution explaining the data is chosen based on a likelihood ratio test (LTR) statistic 2∂L (with a *χ^2^* distribution with degrees of freedom equal to the differences in the number of parameters) computed among models. Once the most likely model was selected, evidence of positive selection was inferred from the estimates of *ω*: values of *ω* > 1 indicated positive selection, while values of *ω* < 1 indicated purifying selection.

We utilized the branch (free-ratio) and the site models. We did not use the branch-site model as we neither had any phylogenetic background nor an apparent OR or GR gene family clustering to test for selection. The branch model assumes different *ω* ratios for different branches on the phylogeny (Nielsen and Yang 1998; Yang 1998) and was used to detect positive selection acting on particular sequences of the subfamilies without averaging *ω* throughout the phylogenetic tree. Using a Likelihood Ratio Test, the branch model was compared to a null hypothesis (*ω*=1, one-ratio M0; (Nielsen and Yang 1998)).

We also tested the site model, which can identify positively selected sites in a multiple-sequence alignment (Yang and Nielsen 2002). The models can be site-class-specific, all of which assume that the *ω* ratio is the same across branches of the phylogeny but different among sites in the alignment. These codon substitution models are M0 (one-ratio), M1a (nearly neutral), M2a (positive selection), M3 (discrete), M7 (beta), M8 (beta and *ω* > 1), and M8a (beta and *ω* = 1). We applied the LTR statistics to evaluate the fit of the nested models to the data for positive selection (Nielsen and Yang 1998) confronting M0 vs. M3, M1a vs. M2a, and M7 vs. M8 models. Support for positive selection can be identified if M2a provides a better fit than M1a or M8 provides a better fit than M7 or M8a (Yang et al. 2000; Anisimova et al. 2001). We confirmed signals of positive selection when a better fit of the selected model was supporting M2a and M8.

The distribution of *dN/dS* ratio estimated from branch models for each gene among OR, GR, and Orthologous Groups (OG) families and species was analyzed with the following the general linear model (function *glm.nb* in R package lme4) as follows :

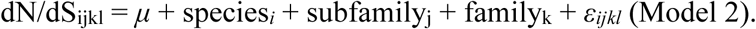

Where dN/dS*_ijkl_* is the *dN/dS* ratio of the sequence from subfamily *j* in the OR/GR/OG family *k* of species *i*, *μ* represents the overall mean, species and families are treated as fixed effects, and *εᵢⱼₖ_l_* is the residual. The residual was fitted to a Negative Binomial using the R package Mass, version 7.3-58.3 (Venables and Ripley 2013). We compared the fit of three models to the data using the *anova* function (aov R package, (Chambers et al. 2017)). The statistical significance of an effect was assessed with an ANOVA-type test and Tukey’s Honest Significance Test (R function Tukey-HSD) to compare *dN/dS* among gene families (OG, GR, OR) pairwise. Predicted *dN/dS* values from the best were then plotted with the plot_model function (sjPlot R package, (Lüdecke 2013)).

### Repetitive DNA element annotation

Examining transposable element (TE) content and evolutionary patterns around OR and GR genes in various aphid species can provide insights into the role of TEs driving OR and GR gene diversification. However, variability in TE annotation methods across aphid genomes poses a challenge for direct comparisons. Standardizing annotation tools is crucial for understanding TE-mediated evolution in these genes and clarifying the genetic mechanisms underlying host adaptation in aphids.

We used the REPET package version 3.0 (https://urgi.versailles.inra.fr/Tools/REPET) (Quesneville et al. 2005; Flutre et al. 2011) for the de novo identification and annotation of TEs in the 13 genomes under survey. REPET includes two main pipelines: TEdenovo (Flutre et al. 2011), which detects repeated elements in a genome to build a TE reference library, and TEannot (Quesneville et al. 2005), which annotates TEs based on a TE consensus library. Consensus sequences were built from at least three repeated sequences identified in the genome using the TEdenovo pipeline (Flutre et al. 2011). The GROUPER program (Quesneville et al. 2005) was used for the TEdenovo High Scoring Pairs (HSP) clustering step. These consensus sequences were then classified using the Wicker code (Wicker et al. 2007) and PASTEC (Hoede et al. 2014). We used the TEannot pipeline (Quesneville et al. 2005) to annotate all repetitive elements in the genome, utilizing the TE consensus library from TEdenovo. TE annotation was done using similarity searches with BLASTER (Quesneville et al. 2003), RepeatMasker (Smit et al. 2015), and CENSOR (Jurka et al. 1996). Short simple repeats (SSRs) were annotated using TRF (Benson 1999) and RepeatMasker (Smit et al. 2015) and removed from the TE annotations. Fragments of the same copy were joined using MATCHER (Quesneville et al., 2003, 2005) and the “long join procedure” (Ahmed et al. 2011). To improve the quality of the TE reference library from TEdenovo, we applied a “second TEannot process” (Jamilloux et al. 2017). The first round of TEannot identifies consensuses that annotate at least one full-length copy (FLC), with both fragmented and unfragmented annotations aligned over more than 95% of the consensus TE sequence. The second round of TEannot is then performed using only those consensuses that annotated at least one FLC.

We test whether the TE content order differed among aphid species using a general linear model (*glm*) with a quasipoisson distribution and ANOVA test with R/Rstudio (http://cran.r-project.org/mirrors.html) as follows:

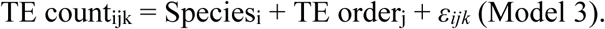

The TE count represented the number of TE copies, *species* indicated the species analyzed, and *TEorder referred to* the TE orders corresponding to each TE copy. We also retrieved the sequences of the TE copies identified by REPET to prepare the consensus. We compared the identity of the TE copies among species using an upset plot (UpSetR R package).

### TE family divergence

We compared TE content and age along the 12 aphid and the one aphid-like genomes. We calculated the Kimura distances (sequence divergence) using scripts of the RepeatMasker package (Flynn et al. 2020). We used the RepeatMasker annotation as input to generate the landscape of aphid species TEs. Kimura’s two-parameter (*K2P*) model is a nucleotide substitution model for estimating genetic distances and phylogenetic relationships, considering transitional and transversional substitution rates (Kimura 1980). We plot the proportion of base pair repeats (Y-axis) at different levels of divergence (X-axis) calculated as a *2-p* Kimura distance using the ggplot2 R package (Wickham 2016). We also utilized the ggplot2 R package to create a plot describing the aphid TE content distribution using the Kimura substitution level as a function of the percentage of the genome. We defined the TE content according to the Wicker TE classification (Wicker et al. 2007) utilized in the PASTEC classifier pipeline from the REPET package (Quesneville et al. 2005).

### TE enrichment near OR and GR genes

We used the locus overlap analysis within the LOLA R package Bioconductor (Sheffield and Bock 2016) to test for TE enrichment near OR/GR genes in the 12 aphid and the one aphid-like genomes. We built three different datasets to launch the analysis. The query set contained genomic regions of 10 Kb around each OR/GR gene. The region universe (or the control genomic region) comprised gene coordinates of all genic areas of the genome and expanded 10 Kb around each gene. We tested for TE enrichment within OR/GR regions compared to the control regions using the Fisher’s Exact Test with false discovery rate correction (*q-value*<0.1). We used the TEGRIP algorithm (Meguerditchian et al. 2021) to characterize relationships between enriched TEs and the chemosensory genes to which they are associated: “subset” means that the TE is included inside the gene, “superset” that the gene is included inside the TE, “upstream” indicate a TE located upstream of the gene, otherwise “downstream”. Finally, “overlap” denotes at least one base of the gene overlapping the downstream or upstream part of the TE. As TEgrip only returns the closest TE to the gene, we used a modified script to conserve all TEs (https://forgemia.inra.fr/sergio.olvera-vazquez/aphid_chemosensory_genes/-/blob/main/Genomic_architecture_TE/relationships_analysis.R). We also calculated Kimura distances between each copy of enriched TE and the consensus sequences using ape R package (Paradis and Schliep 2019) (https://forgemia.inra.fr/sergio.olvera-vazquez/aphid_chemosensory_genes/-/blob/main/Kimura_distance/Kimura_distances.R). For TE detected enriched and without classification following the REPET pipeline, we use DeepTE to improve its classification (Yan et al. 2020). Statistical differences between the distribution of Kimura distances of TE copies associated with GR genes and those associated with OR genes were computed using Wilcoxon signed-rank test (Wilcoxon 1945).

## Data availability

The newly assembled genome of *D. plantaginea* is publicly available on the BIPAA platform (https://bipaa.genouest.org/sp/dysaphis_plantaginea/). The Transposable element consensus sequence and annotations are available on RepetDB (https://urgi.versailles.inrae.fr/repetdb); see detailed URLs in Table S27. The OR/GR CDS sequence and gene structure annotations files are available on the ZENODO repository: DOI 10.5281/zenodo.12750441. All analysis files and codes are available at Forgemia (https://forgemia.inra.fr/sergio.olvera-vazquez/aphid_chemosensory_genes).

## Acknowledgments

We appreciate the support of Alexandre Degrave (Université Angers, Institut Agro, INRAE, IRHS, SFR QUASAV), Claire Mottet, Elorri Segura (Unité Résistance aux Produits Phytosanitaires, Laboratoire de Lyon, ANSES), and Christelle Buchard (Institut de Génétique, Environnement et Protection des Plantes, INRAE, Institut Agro, Université Rennes) to get the *D. plantaginea* individuals for the *de novo* assembly. We thank the Bioinformatics Platform for Agroecosystem Arthropods (BIPAA) platform, particularly Fabrice Legeai (Institut de Génétique, Environnement et Protection des Plantes, INRAE, Institut Agro, Université Rennes) and the Centre National de Ressources Génomiques Végétales-CNRGV, especially William Marande, Nathalie Rodde, and Stephane Cauet (INRAE) in the *de novo* assembly of *D. plantaginea*. We thank the Plant Bioinformatics Facility, BioinfOmics, INRAE, Université Paris-Saclay. We thank Hugh David Loxdale for the discussion on specialization in aphids.

## Author contributions

AC obtained funding, and AC, EJJ, and CM designed the experiment. CB, DO, AD, ES, BB, FL, and AC prepared the material for genome assemblies; SGOV, AC, XC, AM, FAS, JC, CL, IE, CM, EJJ, and YB analyzed the data; All co-authors discussed the results. SGOV, AM, and AC wrote the manuscript with critical input from other co-authors.

## Funding

This research was funded by the ATIP-CNRS Inserm, IDEEV, LabEx BASC, and Tamkeen, under the grant AD454 from the New York University Abu Dhabi Research Institute.

## Conflict of interest

The authors of this preprint declare that they have no financial conflict of interest with the content of this article.

## Notes

### Competing Interest Statement

The authors have declared no competing interest.

