## Supplementary Material for "Comprehensive annotation of olfactory and gustatory receptor genes and transposable elements revealed their evolutionary dynamics in aphids"

**Supplementary materials Parts 1,2,3**

**Table S1.** Details of the 13 genomes (12 aphid and one aphid-like species) used in this study, including their genome sizes, assembly quality, and chemosensory gene and transposable elements contents (REPET pipeline (Quesneville et al. 2005; Flutre et al. 2011)).

**Text S1.** Protocols for DNA extraction and genome sequencing and assembly for *Dysaphis plantaginea*.

**Table S2.** Main statistics of the *Dysaphis plantaginea* assembly versions 1, 2, and 3.

**Table S3.** Statistics for the *Dysaphis plantaginea* genome assembly version 3 and transposable elements content.

**Figure S1.** Workflow for manual annotation of olfactory and gustatory genes across the 12 aphid genomes, and one aphid-like genome.

**Figure S2.** Number of olfactory and gustatory genes detected in 12 aphid species and one aphid-like species.

**Figure S3.** Distribution of the number of olfactory and gustatory genes detected for the 12 aphid species and one aphid-like species predicted with Model 1 (Table S4). Gene count was fitted to a Negative Binomial generalized linear model (glm.nb function).

**Table S4.** Statistical test of the differences in gene numbers among species and gene families (OR and GR) annotated in the 12 aphid species and one aphid-like species. Gene count was fitted to a Negative Binomial generalized linear model (glm.nb function).

**Table S5.** Summary of the orthogroups detected by OrthoFinder V (Emms and Kelly, 2019) for the aphid species.

**Figure S4.** Gene orthology among aphid and one aphid-like species. Bars represent the gene number of different types of orthologues: multi-copy orthologous genes in all species, species-specific (unique genes to specific species), and unassigned (genes cannot be assigned to any orthogroups).

**Table S6.** Information about the aphid host preferences and life cycle used in this study.

**Table S7.** Likelihood-ratio test of models with different evolutionary rates of the 12 aphids and one aphid-like species depending on the taxonomic ranks (genus, subfamilies, families, species) or host preferences (see details in Table S6) or with a single evolutionary rate along the phylogeny inferred with CAFE.

**Figure S5.** Gene ontology analysis of the 147 significantly expanded gene families in the aphid and aphid-like species.

**Figure S6.** Gene ontology analysis of the 50 significantly contracted gene families in the aphid and aphid-like species.

**Table S8.** Enrichment analysis of expanded gene families in the 12 aphid species genomes inferred with CAFE.

**Table S9.** Enrichment analysis of contracted gene families in the 12 aphid species genomes inferred with CAFE**.**

**Table S10.** Gene clusters detected by Syntenet (Almeida-Silva et al. 2023) which overlap with orthogroups detected with OrthoFinder (Emms and Kelly 2019), including the chemosensory genes, in the 12 aphid species and one aphid-like species.

**Table S1.** Details of the 13 genomes (12 aphid and one aphid-like species) used in this study, including their genome sizes, assembly quality, and chemosensory gene and transposable contents (REPET pipeline, Quesneville et al., 2005; Flutre et al., 2011).

| **Aphid species information** | | | | | |
|---|---|---|---|---|---|
| **Species** | **Common name** | **Acronyms** | **Publication** | **Raw data coverage for assembly** | **Genome Size (Mb)** |
| *Aphis glycines (clone BT1)* | Soybean aphid | AGLY | Mathers, 2020 | 5X | 308 |
| *Aphis gossypii* | Cotton aphid | AGOS | Quan et al., 2019 | - | 294 |
| *Acyrthosiphon pisum (clone LSR1)* | Pea aphid | APIS | The International Aphid Genomics Consortium, 2010 | 39X | 541 |
| *Diuraphis noxia* | Russian wheat aphid | DNOX | Nicholson et al., 2015 | 53X | 393 |
| *Dysaphis plantaginea* | Rosy apple aphid | DPLA | ECLECTIC-GQE-INRAE-CNRS | 19X | 485 |
| *Daktulosphaira vitifoliae* | Grape phylloxera | DVIT | Rispe et al., 2020 | 26X | 283 |
| *Eriosoma lanigerum* | Wolly apple aphid | ELAN | Biello et al., 2021 | 309X | 335 |
| *Myzus cerasi* | Black cherry aphid | MCER | Thorpe et al., 2018 | 82X | 406 |
| *Myzus persicae (clone O)* | Green peach aphid | MPER | Mathers et al., 2021 | 131X | 391 |
| *Pentalonia nigronervosa* | Bannana aphid | PNIG | Mathers et al., 2020 | 100X | 375 |
| *Rhopalosiphum maidis* | Maize aphid | RMAI | Chen et al., 2019 | 21X | 326 |
| *Rhopalosiphum padi* | Bird-cherry oat aphid | RPAD | Thorpe et al., 2018 | 761X | 322 |
| *Sitobion miscanthi* | Indian grain aphid | SMIS | Jiang et al., 2019 | 172X | 398 |

**Table S1 continuation.** Details of the 13 genomes (12 aphid and one aphid-like species) used in this study, including their genome sizes, assembly quality, and chemosensory gene and transposable contents (REPET pipeline, Quesneville et al., 2005; Flutre et al., 2011).

| **Scaffold assembly information** | | | | | | | | |
|---|---|---|---|---|---|---|---|---|
| **Acronyms** | **Total assembly size (bp)** | **Num scaffolds** | **Longest scaffold** | **Shorter scaffold** | **Mean scaffold size** | **N50 scaffold (bp)** | **L50 scaffold (bp)** | **scaffold GC (%)** |
| AGLY | 308,065,118 | 3,224 | 23,246,776 | 60 | 95,554 | 5,864,887 | 15 | 26.41 |
| AGOS | 294,278,726 | 4,718 | 5,574,629 | 889 | 2,91 | 437,96 | 195 | 25.78 |
| APIS | 541,137,574 | 21,92 | 170,740,645 | 200 | 24,687 | 132,544,852 | 2 | 29.63 |
| DNOX | 395,073,589 | 5,637 | 2,142,037 | 928 | 70,086 | 397,774 | 281 | 21.78 |
| DPLA | 485,979,193 | 2,366 | 88,606,889 | 13 | 205,401 | 17,731,133 | 7 | 27.37 |
| DVIT | 282,671,353 | 10,492 | 2,080,308 | 141 | 26,942 | 341,59 | 236 | 26.63 |
| ELAN | 334,868,377 | 7,146 | 71,231,741 | 1 | 46,861 | 62,861,676 | 3 | 25.35 |
| MCER | 405,711,039 | 49,286 | 265,361 | 1,001 | 8,232 | 23,273 | 4472 | 29.86 |
| MPER | 390,790,610 | 360 | 105,178,091 | 5 | 1,097,619 | 69,480,500 | 3 | 30.21 |
| PNIG | 375,348,459 | 18,348 | 631,822 | 1 | 20,457 | 103,994 | 1038 | 28.64 |
| RMAI | 326,023,155 | 220 | 94,224,415 | 1,096 | 1,481,923 | 93,298,903 | 2 | 27.69 |
| RPAD | 321,589,008 | 2,172 | 4,088,110 | 1,131 | 148,061 | 652,723 | 140 | 27.81 |
| SMIS | 397,852,643 | 653 | 101,470,385 | 2,682 | 609,269 | 36,263,045 | 4 | 30.06 |

**Table S1 continuation.** Details of the 13 genomes (12 aphid and one aphid-like species) used in this study, including their genome sizes, assembly quality, and chemosensory gene and transposable contents (REPET pipeline, Quesneville et al., 2005; Flutre et al., 2011).

| **Contigs assembly information** | | | | | | | | |
|---|---|---|---|---|---|---|---|---|
| **Acronyms** | **Number of contigs** | **Total size of contigs** | **Longest contigs** | **Shortest contigs** | **Mean contig size** | **N50 contig (bp)** | **L50 contig (bp)** | **contig GC (%)** |
| AGLY | 3,587 | 302,879,913 | 7,698,601 | 60 | 84,438 | 1,958,040 | 47 | 27.27 |
| AGOS | 12,144 | 278,291,622 | 709,413 | 415 | 22,916 | 77,914 | 960 | 27.26 |
| APIS | 68,185 | 499,739,085 | 424,12 | 1 | 73,29 | 25,858 | 4,765 | 29.65 |
| DNOX | 50,723 | 296,543,584 | 167,15 | 60 | 5,846 | 13,309 | 5,772 | 29.01 |
| DPLA | 3,398 | 441,227,050 | 5,044,119 | 12 | 12,9849 | 539,163 | 186 | 30.14 |
| DVIT | 17,162 | 276,429,103 | 718,286 | 83 | 16,107 | 74,75 | 1,012 | 27.23 |
| ELAN | 9,919 | 327,134,977 | 2,875,571 | 3 | 32,981 | 60,282 | 163 | 25.95 |
| MCER | 51,353 | 405,518,793 | 209,856 | 1,001 | 7,897 | 19,719 | 5,425 | 29.88 |
| MPER | 914 | 394,755,899 | 18,090,249 | 2,301 | 43,1899 | 4,169,528 | 23 | 30.23 |
| PNIG | 20,87 | 375,096,534 | 400,055 | 1 | 3,822 | 64,061 | 1,681 | 28.66 |
| RMAI | 689 | 325,976,255 | 42,509,236 | 1,096 | 45,424 | 9,046,396 | 10 | 27.69 |
| RPAD | 2,172 | 321,589,008 | 4,088,110 | 1,131 | 14,8061 | 652,723 | 140 | 27.81 |
| SMIS | 1,145 | 397,803,443 | 10,203,765 | 2,682 | 34,7427 | 1,613,763 | 58 | 30.06 |

**Table S1 continuation.** Details of the 13 genomes (12 aphid and one aphid-like species) used in this study, including their genome sizes, assembly quality, and chemosensory gene and transposable contents (REPET pipeline, Quesneville et al., 2005; Flutre et al., 2011).

| **BUSCO statistics** | | | | | | | | | | | | |
|---|---|---|---|---|---|---|---|---|---|---|---|---|
| **Acronyms** | **Number of BUSCOs** | **Completed BUSCOs (C)** | **%C** | **Completed and single-copy BUSCOs (S)** | **%S** | **Completed and Duplicated BUSCOs (D)** | **%D** | **Fragmented BUSCOs (F)** | **%F** | **Missing BUSCOs (M)** | **%M** | **Total BUSCOs groups searched** |
| AGLY | 2510 | 2427 | 96.7 | 2404 | 95.8 | 23 | 0.9 | 0 | 0.0 | 83 | 3.3 | 2510 |
| AGOS | 2510 | 2442 | 97.3 | 2371 | 94.5 | 71 | 2.8 | 14 | 0.6 | 54 | 2.1 | 2510 |
| APIS | 2510 | 2486 | 99.0 | 2443 | 97.3 | 43 | 1.7 | 4 | 0.2 | 20 | 0.8 | 2510 |
| DNOX | 2510 | 2432 | 96.9 | 2413 | 96.1 | 19 | 0.8 | 38 | 1.5 | 40 | 1.6 | 2510 |
| DPLA | 2510 | 2495 | 99.4 | 2424 | 96.6 | 71 | 2.8 | 4 | 0.2 | 11 | 0.4 | 2510 |
| DVIT | 2510 | 2480 | 98.8 | 2428 | 96.7 | 52 | 2.1 | 7 | 0.3 | 23 | 0.9 | 2510 |
| ELAN | 2510 | 2484 | 99.0 | 2484 | 97.7 | 32 | 1.3 | 9 | 0.4 | 17 | 0.6 | 2510 |
| MCER | 2510 | 2472 | 96.8 | 2430 | 96.8 | 42 | 1.7 | 21 | 0.8 | 17 | 0.7 | 2510 |
| MPER | 2510 | 2484 | 99.0 | 2474 | 98.6 | 10 | 0.4 | 2 | 0.1 | 24 | 0.9 | 2510 |
| PNIG | 2510 | 2491 | 99.3 | 2477 | 98.7 | 14 | 0.6 | 7 | 0.3 | 12 | 0.4 | 2510 |
| RMAI | 2510 | 2489 | 99.1 | 2463 | 98.1 | 26 | 1.0 | 5 | 0.2 | 16 | 0.7 | 2510 |
| RPAD | 2510 | 2487 | 99.1 | 2462 | 98.1 | 25 | 1.0 | 7 | 0.3 | 16 | 0.6 | 2510 |
| SMIS | 2510 | 2402 | 95.7 | 2313 | 92.2 | 89 | 3.5 | 7 | 0.3 | 101 | 4.0 | 2510 |

**Table S1 continuation.** Details of the 13 genomes (12 aphid and one aphid-like species) used in this study, including their genome sizes, assembly quality, and chemosensory gene and transposable contents (REPET pipeline, Quesneville et al., 2005; Flutre et al., 2011).

| | **Chemosensory Genes** | | **Transposable elements (TE)** | | | |
|---|---|---|---|---|---|---|
| **Acronyms** | **GR** | **OR** | **TEs in genome (Mb)** | **TE (%)** | **TE Class I (%)** | **TE Class II(%)** |
| AGLY | 58 | 47 | 170 | 55.98 | 39.31 | 16.67 |
| AGOS | 49 | 55 | 57 | 43.07 | 18.98 | 24.09 |
| APIS | 71 | 70 | 150 | 46.10 | 21.40 | 24.70 |
| DNOX | 26 | 30 | 50 | 52.36 | 26.46 | 28.90 |
| DPLA | 40 | 35 | 255 | 53.42 | 12.94 | 40.48 |
| DVIT | 18 | 64 | 200 | 34.82 | 20.31 | 14.51 |
| ELAN | 23 | 43 | 225 | 41.35 | 15.79 | 25.56 |
| MCER | 40 | 65 | 89 | 45.25 | 14.83 | 30.42 |
| MPER | 56 | 48 | 192 | 53.79 | 20.78 | 33.01 |
| PNIG | 37 | 37 | 94 | 64.27 | 17.95 | 46.32 |
| RMAI | 47 | 52 | 160 | 43.24 | 22.23 | 21.01 |
| RPAD | 64 | 61 | 148 | 44.66 | 14.98 | 29.68 |
| SMIS | 51 | 61 | 184 | 43.46 | 19.38 | 24.58 |

**Table S2.** Main statistics of the *Dysaphis plantaginea* assembly versions 1, 2, and 3.

| ***Dysaphis plantaginea*** **version** | **Assembler** | **Number scaffolds** | **Scaffolds (Mb)** | **Longest scaffold (Mb)** | **N50 (Mb)** | **L50** | **BUSCO genes** | **Complete and single-copy BUSCOs (S)** | **Complete and duplicated BUSCOs (D)** | **Fragments BUSCOs (F)** | **Missing BUSCOs (M)** |
|---|---|---|---|---|---|---|---|---|---|---|---|
| 1 | Supernova 100X | 24,694 | 446.4 | 10.7 | 2.5 | 48 | 94.1% | 22,655 | NA | 337 | 1,083 |
| 1 | Supernova 56X | 24,075 | 389.2 | 15.2 | 2.1 | 41 | 92.9% | 22,366 | NA | 554 | 1,156 |
| 2 | Wtdbg | 3,458 | 411.7 | 13.8 | 2.5 | 40 | ~75% | 1,188 | 48 | 14 | 117 |
| 3 | FLYE | 2,366 | 486 | 88.6 | 17.7 | 7 | ~96% | 1,251 | 63 | 10 | 43 |

**N50: the sequence length of the shortest contig at 50% of the total assembly length; L50: count of the smallest number of contigs whose length sum makes up half of genome size; BUSCO: Benchmarking Universal Single-Copy Orthologs.**

**Table S3.** Statistics for the *Dysaphis plantaginea* genome assembly version 3 and transposable elements content predicted with REPET (Quesneville et al. 2005; Flutre et al. 2011).

| | | | | | | | |
|---|---|---|---|---|---|---|---|
| **Genome size** | **485 Mb** | | | | | | |
| **Assembly information** | | | **Contig information** | | | **BUSCO analysis** | |
| **Total assembly size** | **485,979,193** | | **Num contigs** | **3,398** | | **Completed BUSCOs (C)** | **2,495 (99.4%)** |
| **Num scaffolds** | **2,366** | | **Total contig size** | **441,227,050** | | **Completed and single-copy BUSCOS (S)** | **2,424 (96.6%)** |
| **Longest scaffold** | **88,606,889** | | **Longest contig** | **5,044,119** | | **Completed and duplicated BUSCOs (D)** | **71 (2.8%)** |
| **Shorter scaffold** | **13** | | **Shorter contigs** | **12** | | **Fragmented BUSCOs (F)** | **4 (0.2%)** |
| **Mean scaffold size** | **205,401** | | **Mean contig size** | **12,9849** | | **Missing BUSCOs (M)** | **11 (0.4%)** |
| **N50 scaffold (bp)** | **17,731,133** | | **N50 contig (bp)** | **539,163** | | | |
| **L50 scaffold (bp)** | **7** | | **L50 contig (bp)** | **186** | | **Transposable elements content** | |
| **GC (%)** | **27.37** | | **GC (%)** | **30.14** | | **TEs in genome (Mb)** | **255** |
| | | | | | | **TEs in genome (%)** | **63.78** |

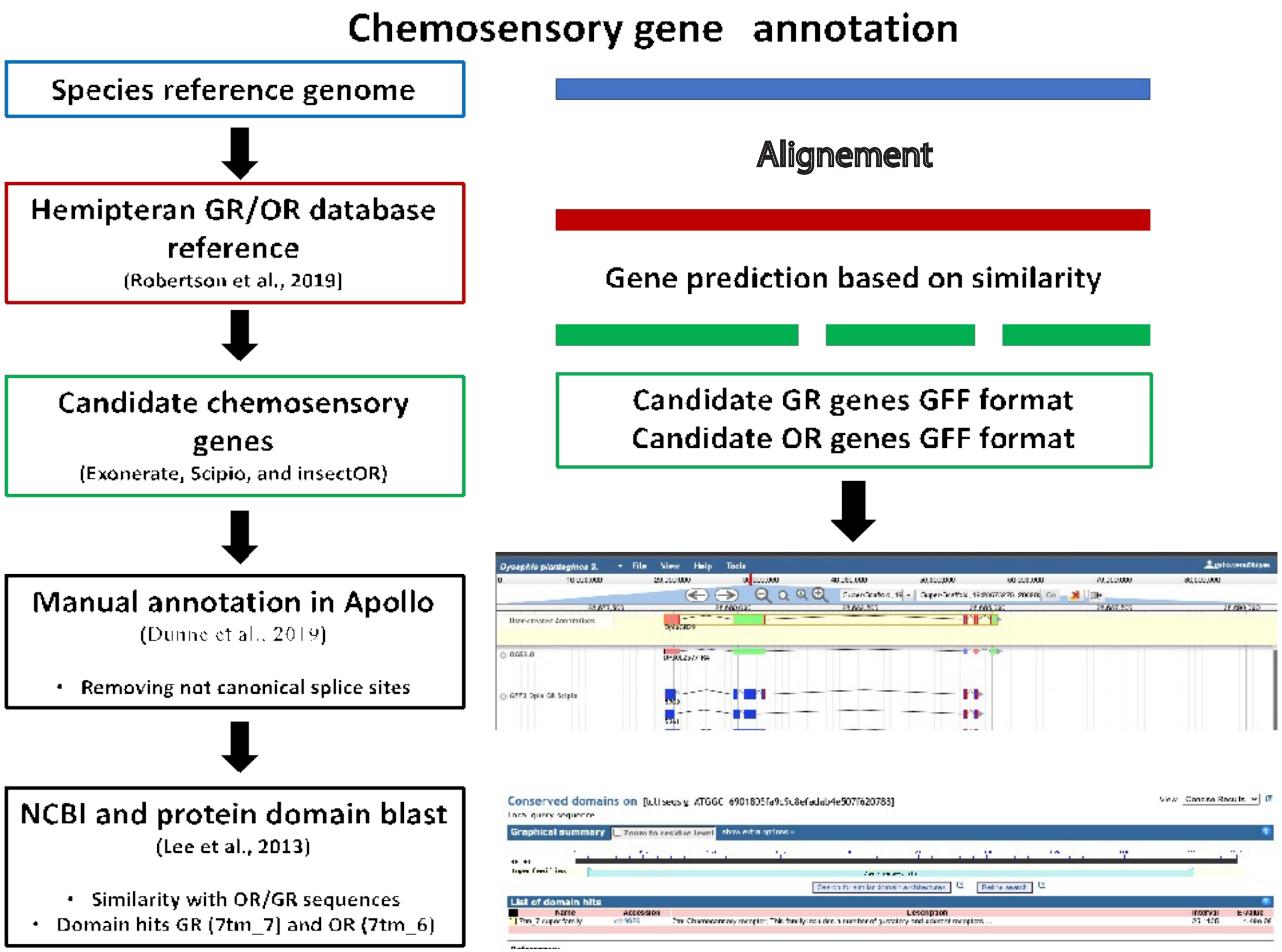

**Figure S1.** Workflow for manual annotation of olfactory and gustatory genes across the 12 aphid genomes, and one aphid-like genome.

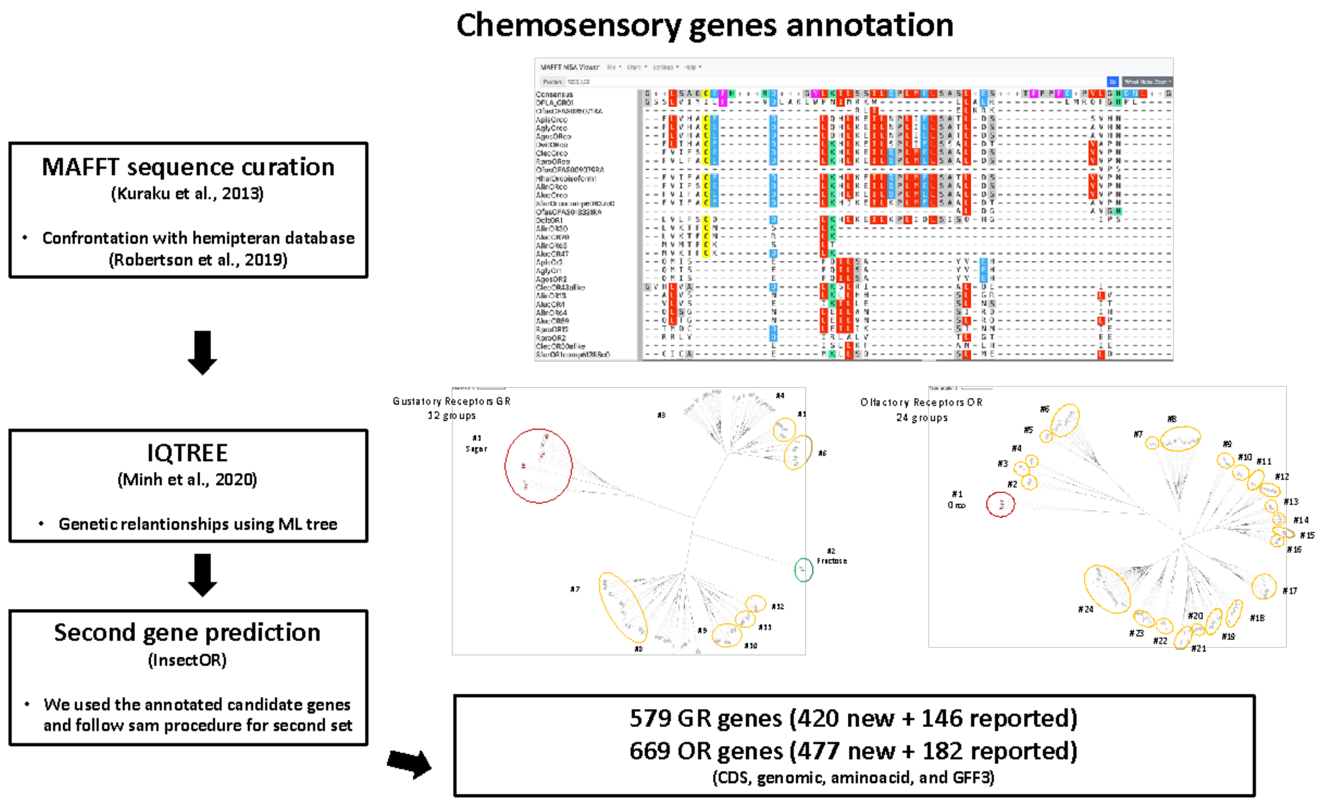

**Figure S1 continuation.** Workflow for manual annotation of olfactory and gustatory genes across the 12 aphid genomes, and one aphid-like genome.

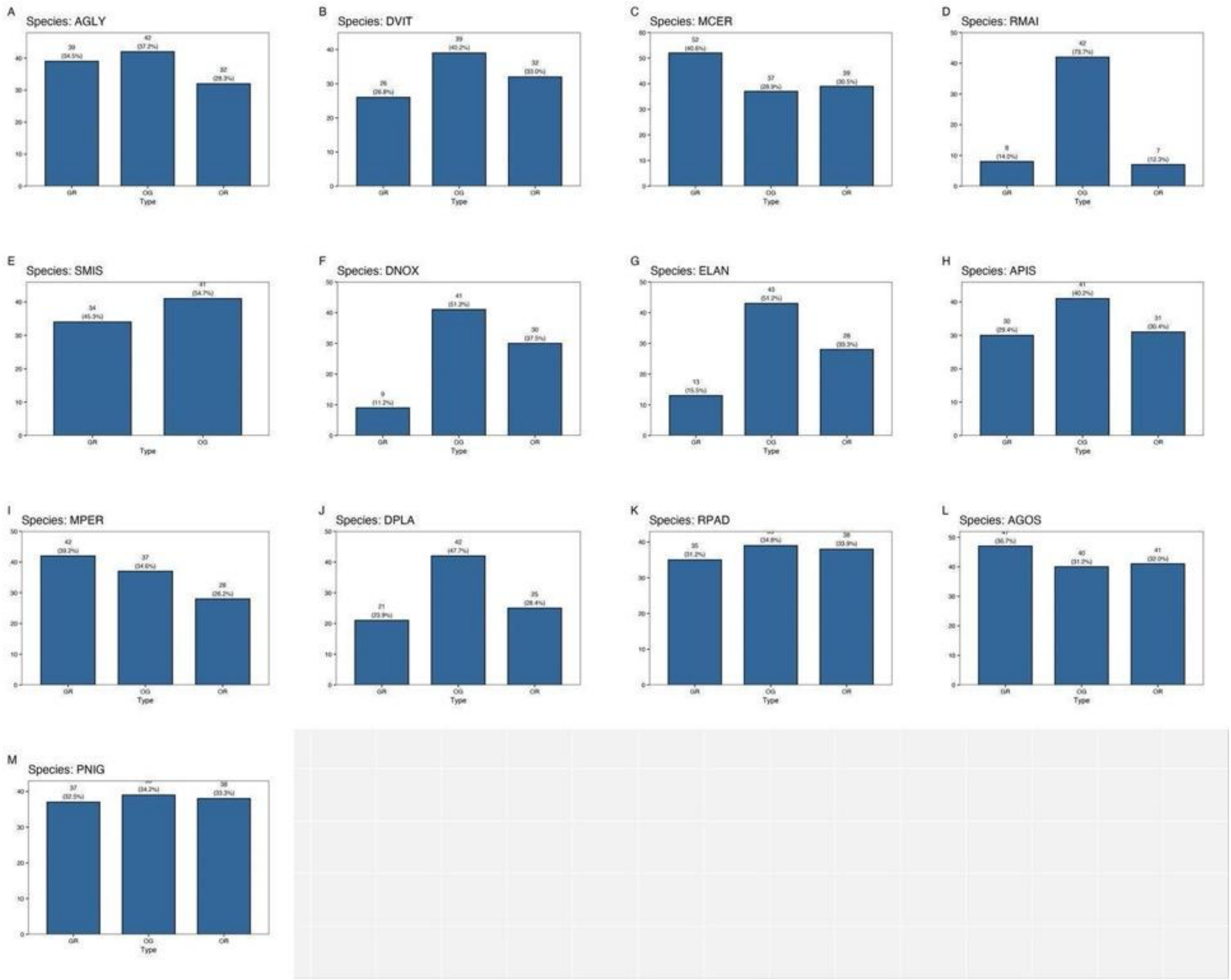

**Figure S2. Number of olfactory and gustatory genes detected in 12 aphid species and one aphid-like species.** This figure displays the counts of olfactory receptor (OR) and gustatory receptor (GR) genes identified across 12 aphid species and one aphid-like species. It also includes the number of single-copy orthologous groups used as references for the genomic background. MCER: *Myzus cerasi;* MPER: *Myzus persicae*; DPLA: *Dysaphis plantaginea;* DNOX: *Diuraphis noxia*; APIS: *Acyrthosiphon pisum*; SMIS: *Sitobion miscanthi;* PNIG: *Pentalonia*

*nigronervosa*; AGLY: *Aphis glycines*; AGOS: *Aphis gossypii*; RMAI: *Rhopalosiphum maidis*; RPAD: *Rhopalosiphum padi* ; ELAN: *Eriosoma lanigerum* ; DVIT: *Daktulosphaira vitifoliae*.

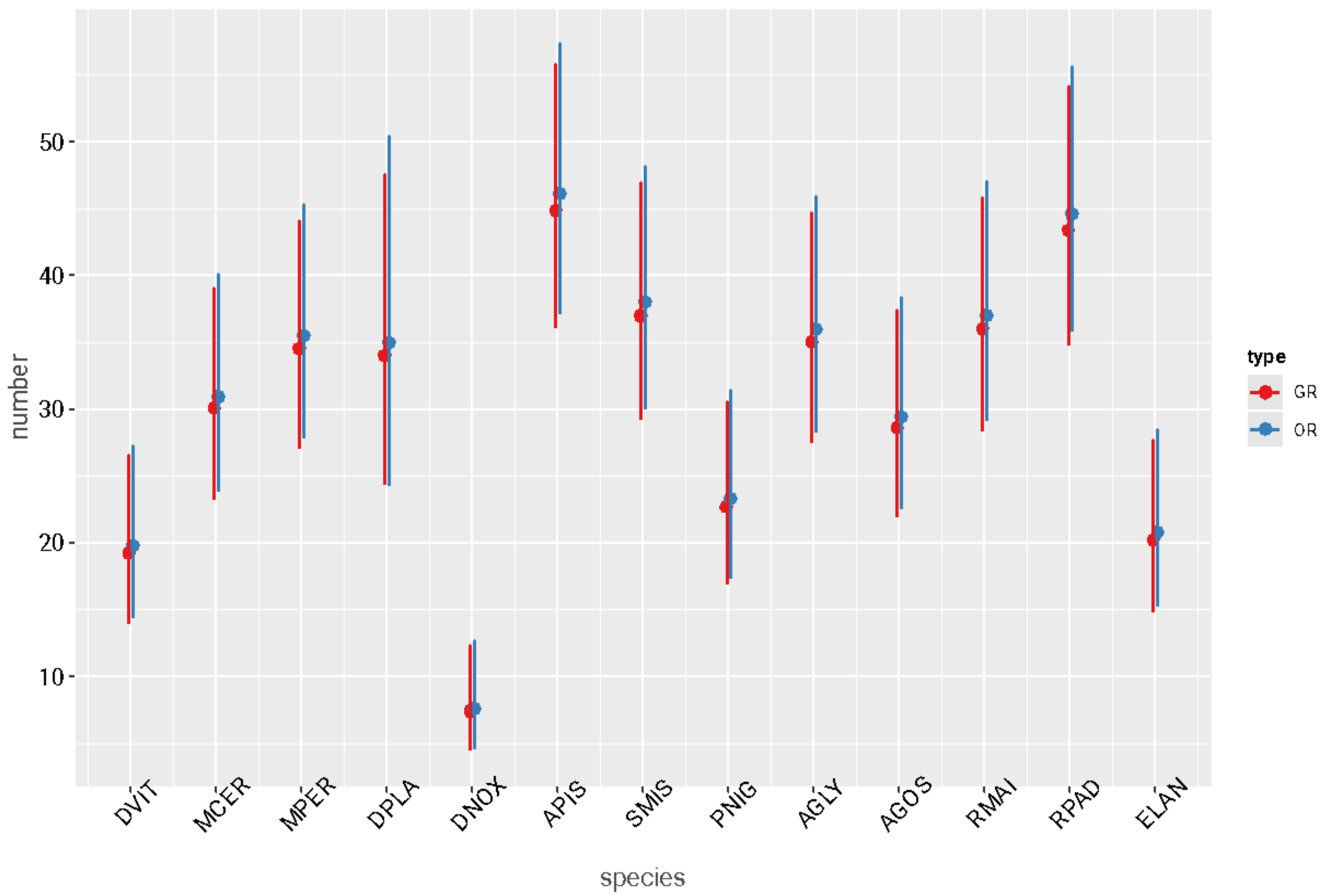

**Figure S3.** Number of olfactory and gustatory genes detected for the 12 aphid species and one aphid-like species predicted with Model 1 (Table S4). MCER: *Myzus cerasi;* MPER: *Myzus persicae*; DPLA: *Dysaphis plantaginea;* DNOX: *Diuraphis noxia*; APIS: *Acyrthosiphon pisum*; SMIS: *Sitobion miscanthi;* PNIG: *Pentalonia nigronervosa*; AGLY: *Aphis glycines*; AGOS: *Aphis gossypii*; RMAI: *Rhopalosiphum maidis*; RPAD: *Rhopalosiphum padi* ; ELAN: *Eriosoma lanigerum* ; DVIT: *Daktulosphaira vitifoliae*.

**Table S4.** Statistical test of the differences in gene numbers among species and gene families (OR and GR) annotated in the 12 aphid species and one aphid-like species. Gene count was fitted to a Negative Binomial generalized linear model (glm.nb function) (Model 1).

| | Df | Deviance Residuals | Pr(>Chi) |
|---|---|---|---|
| **species** | 12 | 102.341 | **<2e-16** |
| **family (GR/OR)** | 1 | 0.137 | 0.7109 |

Df: degree of freedom; Pr(>Chi): P-values.

**Table S5.** Summary of the orthogroups detected by OrthoFinder V (Emms and Kelly, 2019) for the 13 studied species.

| Summary of the orthologous groups detected | |
|---|---|
| **Number of species** | 13 |
| **Number of genes** | 380,653 |
| **Number of genes in orthogroups** | 334,936 |
| **Number of unassigned genes** | 45,717 |
| **Percentage of genes in orthogroups** | 88 |
| **Percentage of unassigned genes** | 12 |
| **Number of orthogroups** | 28,672 |
| **Number of species-specific orthogroups** | 5,713 |
| **Number of genes in species-specific orthogroups** | 28,227 |
| **Percentage of genes in species-specific orthogroups** | 7.4 |
| **Mean orthogroup size** | 11.7 |
| **Median orthogroup size** | 6 |
| **G50 (assigned genes)** | 18 |
| **G50 (all genes)** | 16 |
| **O50 (assigned genes)** | 44,05 |
| **O50 (all genes)** | 5,741 |
| **Number of orthogroups with all species present** | 5,868 |
| **Number of single-copy orthogroups** | 891 |

**Table S5.** Summary of the orthogroups detected by OrthoFinder V (Emms and Kelly, 2019) for the 13 aphid species (continuation)

| **Orthologous groups statistics per species** | | | | | | | | | | | | | |
|---|---|---|---|---|---|---|---|---|---|---|---|---|---|
| | **AGLY** | **AGOS** | **APIS** | **DNOX** | **DPLA** | **DVIT** | **ELAN** | **MCER** | **MPER** | **PNIG** | **RMAI** | **RPAD** | **SMIS** |
| Number of genes | 42244 | 18504 | 28016 | 17484 | 53923 | 25794 | 28310 | 28793 | 31593 | 29706 | 19498 | 26551 | 30237 |
| Number of genes in orthogroups | 28489 | 18236 | 27416 | 17169 | 46719 | 23400 | 23108 | 24686 | 29943 | 27604 | 19376 | 22573 | 26217 |
| Number of unassigned genes | 13755 | 268 | 600 | 315 | 7204 | 2394 | 5202 | 4107 | 1650 | 2102 | 122 | 3978 | 4020 |
| Percentage of genes in orthogroups | 67,4 | 98,6 | 97,9 | 98,2 | 86,6 | 90,7 | 81,6 | 85,7 | 94,8 | 92,9 | 99,4 | 85 | 86,7 |
| Percentage of unassigned genes | 32,6 | 1,4 | 2,1 | 1,8 | 13,4 | 9,3 | 18,4 | 14,3 | 5,2 | 7,1 | 0,6 | 15 | 13,3 |
| Number of orthogroups containing species | 15567 | 10414 | 11495 | 10133 | 18085 | 10884 | 12308 | 14784 | 13863 | 13778 | 10036 | 13753 | 15142 |
| Percentage of orthogroups containing species | 54,3 | 36,3 | 40,1 | 35,3 | 63,1 | 38 | 42,9 | 51,6 | 48,4 | 48,1 | 35 | 48 | 52,8 |
| Number of species-specific orthogroups | 664 | 43 | 270 | 32 | 1550 | 719 | 660 | 186 | 288 | 540 | 35 | 280 | 446 |
| Number of genes in species-specific orthogroups | 4975 | 106 | 1153 | 71 | 5719 | 6067 | 3626 | 506 | 1272 | 1855 | 166 | 1357 | 1354 |
| Percentage of genes in species-specific orthogroups | 11,8 | 0,6 | 4,1 | 0,4 | 10,6 | 23,5 | 12,8 | 1,8 | 4 | 6,2 | 0,9 | 5,1 | 4,5 |

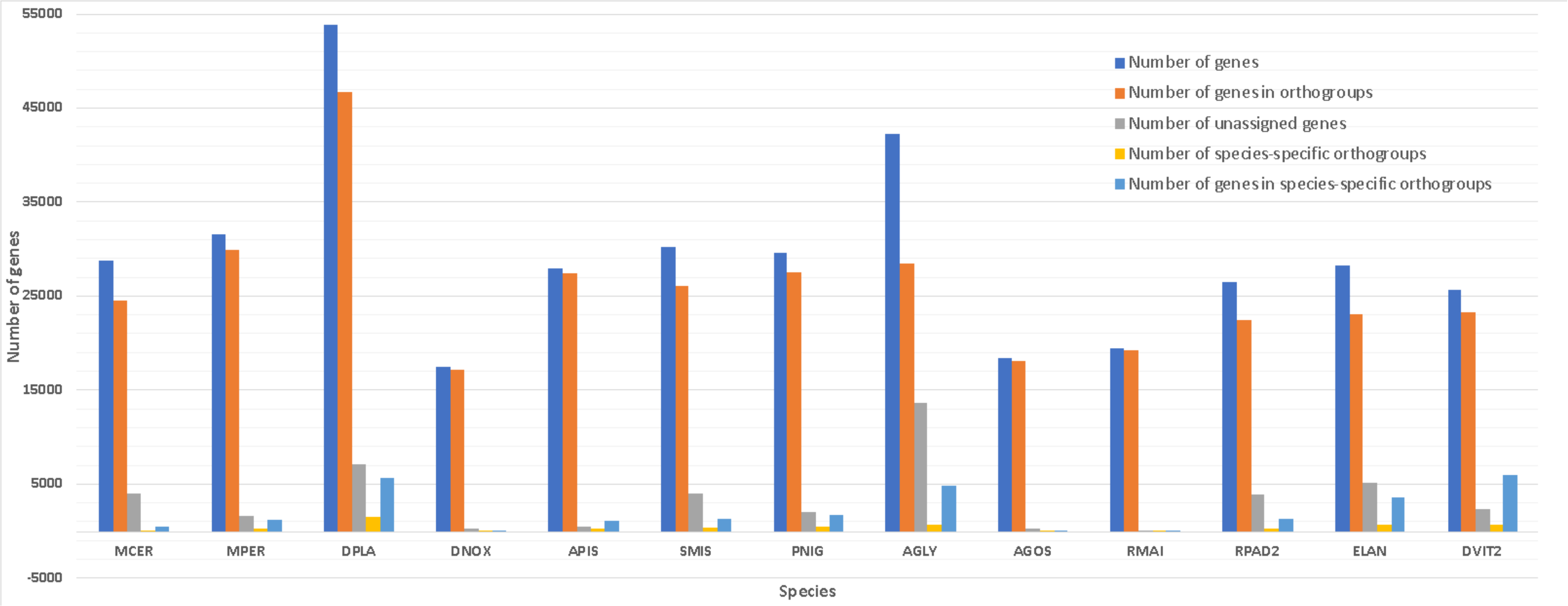

**Figure S4.** Gene orthology among aphid and one aphid-like species. Bars represent the gene number of different types of orthologues: multi-copy orthologous genes in all species, species-specific (unique genes to specific species), and unassigned (genes cannot be assigned to any orthogroups). MCER: *Myzus cerasi;* MPER: *Myzus persicae*; DPLA: *Dysaphis plantaginea;* DNOX: *Diuraphis noxia*; APIS: *Acyrthosiphon pisum*; SMIS: *Sitobion miscanthi;* PNIG: *Pentalonia nigronervosa*; AGLY: *Aphis glycines*; AGOS: *Aphis gossypii*; RMAI: *Rhopalosiphum maidis*; RPAD: *Rhopalosiphum padi* ; ELAN: *Eriosoma lanigerum* ; DVIT: *Daktulosphaira vitifoliae*.

**Table S6.** Information about the aphid host preferences and life cycle used in this study.

| **Aphid host preference information and life cycle** | | | | |
|---|---|---|---|---|
| **Family** | **Subfamily** | **Tribe** | **Genera** | **Species** |
| Aphididae | Aphidinae | Aphidini | *Aphis* | *Aphis glycines* (clone BT1) |
| Aphididae | Aphidinae | Aphidini | *Aphis* | *Aphis gossypii* |
| Aphididae | Aphidinae | Macrosiphini | *Acyrthosiphon* | *Acyrthosiphon pisum* (clone LSR1) |
| Aphididae | Aphidinae | Macrosiphini | *Diuraphis* | *Diuraphis noxia* |
| Aphididae | Aphidinae | Macrosiphini | *Dysaphis* | *Dysaphis plantaginea* |
| Phylloxeridae | Phyloxerininae | Phylloxerini | *Daktulosphaira* | *Daktulosphaira vitifoliae* |
| Aphididae | Eriosomatinae | Eriosomatini | *Eriosoma* | *Eriosoma lanigerum* |
| Aphididae | Aphidinae | Macrosiphini | *Myzus* | *Myzus cerasi* |
| Aphididae | Aphidinae | Macrosiphini | *Myzus* | *Myzus persicae* (clone O) |
| Aphididae | Aphidinae | Macrosiphini | *Pentalonia* | *Pentalonia nigronervosa* |
| Aphididae | Aphidinae | Aphidini | *Rhopalosiphum* | *Rhopalosiphum maidis* |
| Aphididae | Aphidinae | Aphidini | *Rhopalosiphum* | *Rhopalosiphum padi* |
| Aphididae | Aphidinae | Macrosiphini | *Sitobion* | *Sitobion miscanthi* |

**Table S6 continuation.** Information about the aphid host preferences and life cycle used in this study.

| Aphid host preference information and life cycle | | | | | |
|---|---|---|---|---|---|
| **Species** | **Common name** | **Acronyms** | **Genome_publication** | **Genome Size (Mb)** | **Genome_assembly_origin** |
| *Aphis glycines* (clone BT1) | Soybean aphid | AGLY | Mathers, 2020 | 308 | USA, Illinois Urbana |
| *Aphis gossypii* | Cotton aphid | AGOS | Quan et al., 2019 | 294 | Shanghai, China |
| *Acyrthosiphon pisum* (clone LSR1) | Pea aphid | APIS | The International Aphid Genomics Consortium, 2010 | 541 | Itaca, New York |
| *Diuraphis noxia* | Russian wheat aphid | DNOX | Nicholson et al., 2015 | 393 | Biotype2_Oklahoma_USA |
| *Dysaphis plantaginea* | Rosy apple aphid | DPLA | ECLECTIC-GQE-INRAE-CNRS | 485 | France |
| *Daktulosphaira vitifoliae* | Grape phylloxera | DVIT | Rispe et al., 2020 | 283 | Bordeaux, France |
| *Eriosoma lanigerum* | Wolly apple aphid | ELAN | Biello et al., 2021 | 335 | United Kingdonm |
| *Myzus cerasi* | Black cherry aphid | MCER | Thorpe et al., 2018 | 406 | UK |
| *Myzus persicae* (clone O) | Green peach aphid | MPER | Mathers et al., 2021 | 391 | Norwich UK |
| *Pentalonia nigronervosa* | Bannana aphid | PNIG | Mathers et al., 2020 | 375 | Nairobi, Kenya |
| *Rhopalosiphum maidis* | Maize aphid | RMAI | Chen et al., 2019 | 326 | New York, USA |
| *Rhopalosiphum padi* | Bird-cherry oat aphid | RPAD | Thorpe et al., 2018 | 322 | Henan, China |
| *Sitobion miscanthi* | Indian grain aphid | SMIS | Jiang et al., 2019 | 398 | Hebei, China |

Notes. *A. gossypii* feeds on about 900 plants (Blackman and Eastop 2000; Ma et al. 2019), *M. persicae* on around 400 plant species, and *D. noxia* on > 140 plant species (Blackman and Eastop, 2000). We also included polyphagous species with more restricted host ranges, such as the model species *A. pisum*, which feeds on plants from the Fabaceae family, and *A. glycines*, which feeds on *Glycine willd.* spp. and *Rhamnus* L. spp. (Ragsdale et al. 2004; Wang et al. 2019). We included aphid species that primarily infest the Poaceae family, encompassing essential cereal crops like maize, barley, oats, and wheat, such as *R. maidis* (Chen et al. 2019), *R. padi* (Thorpe et al. 2018), and *S. miscanthi* (this species also infests Cyperaceae members; (Jiang et al. 2019)). Additionally, we included more specialized aphids, *D. plantaginea* and *E. lanigerum*, which primarily host on apple trees (*M. domestica* Borkh), but on elm, in North America for *E. lanigerum* (Biello et al. 2021). *Myzus cerasi* (Dondini, L. et al. 2018) infests a limited host range, including cherry trees (*Prunus avium* L. and *Prunus cerasus* L.). The aphid *P. nigronervosa* infests *Musa* L. spp (Mathers et al. 2020), and we also included the aphid-like species *D. vitifoliae,* which feeds on different species of the *Vitis* L. genus (Rispe et al. 2020)

**Table S6 continuation.** Information about the aphid host preferences and life cycle used in this study.

| Acronyms | *Age | *Developmental stage | *Sex | *Host | *Tissue | *Origin | Distribution |
|---|---|---|---|---|---|---|---|
| AGLY | Adult | parthenogenic adult | Female | *Glycine max* | Whole aphid | Asia | Asia, North America, Australia, and Russia |
| AGOS | Adult | Adult | Female | *NA* | Whole aphid | Europe (tentative) | Tropical and temperate region in throughout the world |
| APIS | Adult | Adult | Female | *Medicago sativa* | NA | Europe or Asia | Temperate region in throughout the world |
| DNOX | Adult | Adult | Female | *Triticum aestivum* | Whole aphid | Central Asia | Widespread in southern Europe, Middle East, Central Asia, Africa, South and North America and Australia, but apparently not in western or northern Europe. |
| DPLA | Adult | Adult | Female | *Plantago lanceolata* | Whole aphid | Middle-east (under research) | Europe, Africa, much of Asia and North and South America |
| DVIT | Adult | Mixed leaf gall | Female | *Vitis vinifera* | Whole aphid | North America | North, Central and South America, Europe, the Mediterranean, the Middle East, Africa, China and Australia |
| ELAN | Adult | Adult | Female | *Apple plant* | Whole aphid | North America | Temperate region in throughout the world |
| MCER | Adult | Adult | NA | *Barbarea verna* | Whole aphid | Mediterranean | Temperate region in throughout the world |
| MPER | Adult | Single apterous female | Female | *Brassica rapa* | Whole aphid | Asia | Temperate region in throughout the world |
| PNIG | Adult | Adult | Female | *Musa sp.* | Whole aphid | Africa (tentative) | Tropical and subtropical region in throughout the world |
| RMAI | Adult | Adult | Female | *Zea Mays* | Whole aphid | Asia | Tropical and subtropical region in throughout the world |
| RPAD | Adult | Adult | Female | *Triticum aestivum* | Whole aphid | North America | Temperate region in throughout the world |
| SMIS | Adult | Newborn nymph | Female | *Triticum aestivum* | Whole aphid | Asian | Indian subcontinent, east and south-east Asia and Australasia |

*Information extracted from the genome assembly reference manuscript, genome publication column.

**Table S6 continuation.** Information about the aphid host preferences and life cycle used in this study.

| Acronyms | Life cycle | Reproduction | Host specialization | Host_primary | Host_secondary |
|---|---|---|---|---|---|
| AGLY | Heteroecious | Holocyclic | biotypes reported | *Rhamnus* sp. | Mainly *Glycine max* (soybean) but also *Glycine sp*. and other Fabaceae members |
| AGOS | Monoecious | Holocyclic/Anholocyclic in tropics regions | biotypes reported | Catalpa bignonioides (Indian bean tree), Hibiscus syriacus (Korean rose), Celastrus orbiculatus (oriental bittersweet), Rhamnus species (buckthorns) and Punica granatum (pomegranate) | Cucurbitaceae, Malvaceae, and Rutaceae species |
| APIS | Monoecious | Holocyclic/Anholocyclic in mild climates | biotypes reported | Family Fabaceae, but especially on *Medicago*, *Melilotus*, *Trifolium*, *Dorycnium* and *Lotus* | |
| DNOX | Monoecious | Holocyclic/mainly Anholocyclic | biotypes reported | Grasses and cereals (such as *Agropyron, Bromus, Elymus, Hordeum, Triticum*) | |
| DPLA | Dioecious | Holocyclic | not reported | *Malus domestica* (cultivated apple) | *Plantago lanceolata* (ribwort plantain) |
| DVIT | Monoecious | Anholocyclic | not reported | *Vitis sp.* (grape) | |
| ELAN | Dioecious in America/Monoecious most part of the world | Anholocyclic/Holocyclic (in America) | not reported | *Ulmus americana* (Elm) in America / *Malus domestica* (other parts of the world) | *Malus domestica* (cultivated apple) |
| MCER | Dioecious | Holocyclic | not reported | *Prunus cerasus/Prunus avium* (cherry trees) | bedstraws (*Galium*), eyebrights (*Euphrasia*) and speedwell (*Veronica* sp.) |
| MPER | Heteroecious | Holocyclic/Anholocyclic in *M. persicae nicotianae* | biotypes reported | *Prunus persica* (peach) | Brassicaceae, Solanaceae, Poaceae, Leguminosae, Cypraceae, Convolvulaceae, Chenopodiaceae, Compositae, Cucurbitaceae, and Umbelliferae |
| PNIG | Monoecious | Anholocyclic | not reported | Musaceae | |
| RMAI | Monoecious/Dioecious in Asia | Anholocyclic/Holocyclic (in Asia) | biotypes reported | Poaceae/ *Prunus* sp. (Asia) | Poaceae |
| RPAD | Dioecious | Holocyclic/Anholocyclic in mild climates | not reported | *Prunus padus* (bird cherry) | Poaceae |
| SMIS | Monoecious | Anholocyclic | biotypes reported | | |

Holocyclic: reproduce both asexually and sexually; Anholocyclic: asexual reproduction only; Monoecious: same host throughout their life stages; Dioecious/Heteroecious: host alternates between sexual and asexual stages (Hardy et al., 2015; Emden and Harrington, 2017).

**Table S7.** Likelihood-ratio test of models with different evolutionary rates of the 12 aphids and one aphid-like species depending on the taxonomic ranks (genus, subfamilies, families, species) or host preferences (see details in Table S6) or with a single evolutionary rate along the phylogeny inferred with CAFE.

| **Model** | **P vs. S** | **F vs.S** | **Sf vs. S** | **T vs. S** | **G vs. S** | **F vs. P** | **Sf vs.s P** | **T vs. P** | **P vs. G** | **F vs. Sf** |
|---|---|---|---|---|---|---|---|---|---|---|
| ***Ho*** **(InL)** | 254806 | 247150 | 243475 | 244584 | 266625 | 247150 | **243475** | 244584 | 254806 | 247150 |
| ***Ha*** **(InL)** | 255636 | 255636 | 255636 | 255636 | 255636 | 254806 | **254806** | 254806 | 266625 | 243475 |
| **2ΔL** | 1660 | 16972 | 24322 | 22104 | -21978 | -15312 | **-22662** | -20444 | 23638 | 7350 |
| ***K*** | 5 | 1 | 2 | 3 | 9 | 7 | **6** | 5 | 1 | 1 |
| ***alpha*** | 0.05 | 0.05 | 0.05 | 0.05 | 0.05 | 0.05 | **0.05** | 0.05 | 0.05 | 0.05 |
| ***p-value*** | <2.2e-16 | <2.2e-16 | <2.2e-16 | <2.2e-16 | 1 | 1 | **1** | 1 | <2.2e-16 | <2.2e-16 |
| $\chi^2$ | 11.07 | 3.8415 | 5.9915 | 7.8147 | 16.919 | 14.067 | **12.592** | 11.07 | 3.8415 | 3.8415 |
| **Best hypothesis** | P | F | Sf | T | S | F | **Sf** | T | G | Sf |

| **Model** | **F vs. T** | **F vs. G** | **Sf vs. T** | **Sf vs. G** | **T vs. G** | **P vs. S** | **S vs. Gr** | **S vs. H** | **P vs. Sp** | **P vs. H** | **H vs. Gr** |
|---|---|---|---|---|---|---|---|---|---|---|---|
| ***Ho*** **(InL)** | 247150 | 247150 | 243475 | 243475 | 244584 | 267684 | 255490 | 267678 | 255490 | 267678 | 255490 |
| ***Ha*** **(InL)** | 244584 | 266625 | 244584 | 266625 | 266625 | 284810 | 255490 | 267684 | 284810 | 284810 | 267678 |
| **2ΔL** | 5132 | -38950 | -2218 | -46300 | -44082 | 34252 | 24388 | 12 | 58640 | 34264 | 24376 |
| ***K*** | 2 | 8 | 1 | 7 | 6 | 1 | 1 | 1 | 1 | 1 | 1 |
| ***alpha*** | 0.05 | 0.05 | 0.05 | 0.05 | 0.05 | 0.0500 | 0.0500 | 0.0500 | 0.0500 | 0.0500 | 0.0500 |
| ***p-value*** | <2.2e-16 | 1 | 1 | 1 | 1 | 2.2e-16 | 2.2e-16 | 0.000532 | 2.2e-16 | 2.2e-16 | 2.2e-16 |
| $\chi^2$ | 5.9915 | 15.507 | 3.8415 | 14.067 | 12.592 | 3.8415 | 3.8415 | 3.8415 | 3.8415 | 3.8415 | 3.8415 |
| **Best hypothesis** | T | F | Sf | Sf | T | P | Gr | H | P | P | H |

***Ho*** **(InL) : Nule hypothesis ;** ***Ha*** **(InL) : Alternative hypothesis;** ***K*** **:Degrees of freedom*; alpha : 0.05 ; p-value : <0.05 ;*** $\chi$ **: chi-square distribution; Hypotheses acronyms P=Phylogeny, S=single, F=Family, Sf=Subfamily, T=Tribe, G=Genus, H=Host, Gr=Generalist, Sp : Specialis**

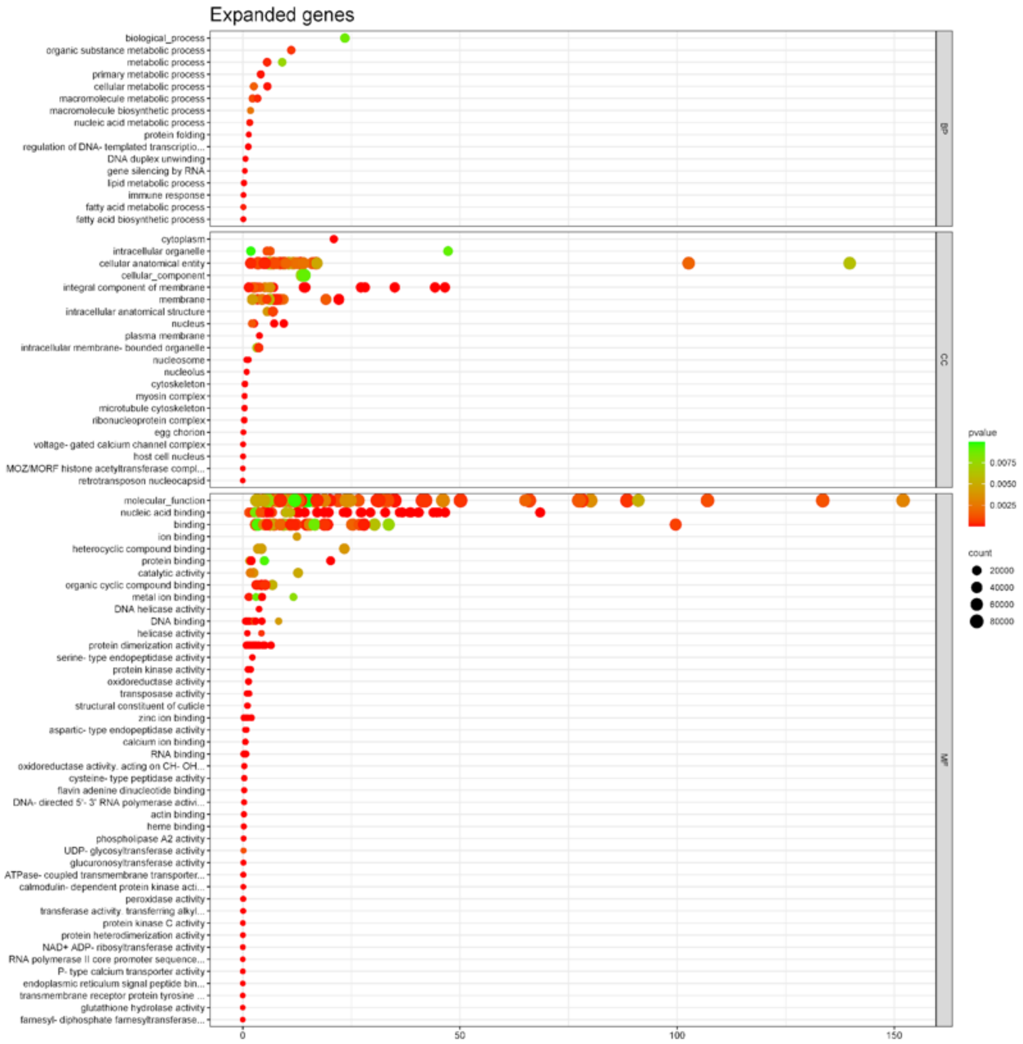

**Figure S5.** Gene ontology analysis of the 147 significantly expanded gene families in the aphid and aphid-like species.

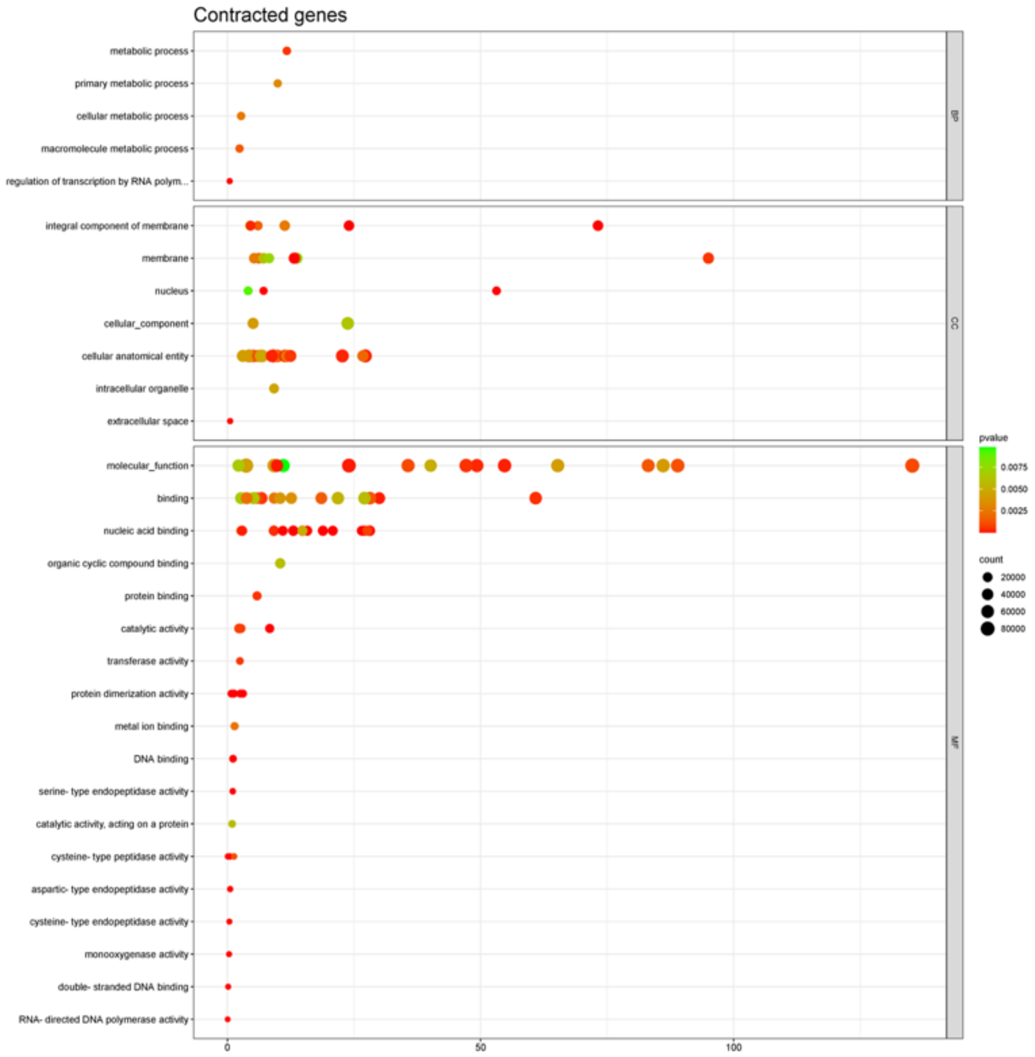

**Figure S6.** Gene ontology analysis of the 50 significantly contracted gene families in the aphid and aphid-like species.

**Table S8.** Enrichment analysis of expanded gene families in the 12 aphid species genomes inferred with CAFE.
See Excel File Table S8.
**Table S9.** Enrichment analysis of contracted gene families in the 12 aphid species genomes inferred with CAFE**.**
See Excel File Table S9.
**Table S10.** Gene clusters detected by Syntenet (Almeida et al., 2023) overlap with orthogroups (OrthoFinder, Emms and Kelly, 2019, including the chemosensory genes)**.** ND = Not defined.
See Excel file Table S10.
