## Supplementary Material for "Comprehensive annotation of olfactory and gustatory receptor genes and transposable elements revealed their evolutionary dynamics in aphids"

**Figure S7.** Length distribution of annotated chemosensory genes for each aphid species studied. Sequences <150pb and with a stop codon were removed and kept for subsequent clustering analysis and subfamily detection.

**Table S11.** Number of annotated GR and OR genes before and after filtering for their length and stop-codon presence for each aphid species to infer the OR and GR subfamilies.

**Figure S8.** Twelve subfamilies of gustatory receptors (GR) genes were detected using a phylogenetic tree built with IQ-TREE v2.0 using 373 amino-acid sequences**.**

**Figure S9.** Twenty-four groups of the candidate olfactory receptors (OR) genes were detected using a phylogenetic tree built with IQ-TREE v2.0 using 393 amino-acid sequences.

**Table S12**. Chemosensory genes (gustatory receptors - GR and olfactory receptors - OR) branch model summary statistics inferred from PAML v4 (Yang, 2007).

**Table S13**. Number of gustatory receptors (GR) and olfactory receptors (OR) subfamilies under positive ($\omega>0$), purifying ($\omega<0$), and neutral ($\omega=0$) selection based on the branch model analysis PAML v4 (Yang, 2007).

**Table S14**. Summary statistics from site-specific models of evolution for olfactory receptor (OR) and gustatory receptor (GR) subfamilies, inferred using PAML v4 (Yang, 2007).

**Table S15.** Likelihood-ratio test results for site-specific evolutionary models of olfactory receptor (OR) and gustatory receptor (GR) subfamilies, conducted using PAML v4 (Yang, 2007).

**Table S16**. Number of gustatory receptors (GR) and olfactory receptors (OR) subfamilies under positive selection (M8 model) based on the site model analysis PAML v4 (Yang, 2007).

**Table S17**. Summary of selection test results based on the branch model applied to single-copy orthologous groups. Selection types identified include positive selection ($\omega > 0$), purifying selection ($\omega < 0$), and neutral selection ($\omega = 0$), as inferred from PAML v4 analysis (Yang, 2007).

**Table S18**. Single-copy orthologous groups (OGs) classified under positive selection ($\omega > 0$), purifying selection ($\omega < 0$), and neutral selection ($\omega = 0$), based on branch model analysis using PAML v4 (Yang, 2007).

**Table S19**. Summary statistics for branch-specific evolutionary models of single-copy orthologous subfamilies, analyzed using PAML v4 (Yang, 2007).

**Table S20.** Likelihood-ratio test results for site-specific evolutionary models of single-copy orthologous subfamilies were conducted using PAML v4 (Yang, 2007).

**Table S21**. Number of gustatory receptors (GR) and olfactory receptors (OR) subfamilies under positive selection (M8 model) and purifying (M0 and M1a, M7) based on the site model analysis PAML v4 (Yang, 2007).

**Table S22**. General linear model (GLM) summary of different models to test the distribution of the selection pressure among the different chemosensory genes and the single-copy orthologous gene groups (OG).

**Figure S10.** Distribution of dN/dS ratios for each olfactory receptor (OR) and gustatory receptor (GR) gene subfamily, as well as for single-copy orthologous gene families across different aphid species. For further details, refer to Table S22 (Model 1)

**Figure S11.** Correlation between the dN/dS ratio and synonymous divergence (dS) among gustatory receptor (GR) and olfactory receptor (OR) gene subfamilies, as well as single-copy orthologous gene families (OG) across 12 aphid species.

**Table S23**. Key statistics from the genome assemblies utilized in the REPET pipeline for analyzing transposable elements across 12 aphid species and one aphid-like species.

**Table S24.** Statistics of the transposable element consensus library from the REPET pipeline of the 12 aphid species and one aphid-like species.

**Table S25**. Transposable elements (TEs) enrichment analysis in the Gustatory receptor (GR) and Olfactory receptor (OR) genes in a 2 kb and 10Kb window detected using Locus Overlap Analysis (LOLA)

**Table S26**. Relationships between enriched transposable elements (TEs) and associated Gustatory receptor (GR) and Olfactory receptor (OR) genes estimated using TEgriP.

**Figure S12.** Transposable elements (TEs) content among twelve aphid species and one aphid-like species represented with the kimura distribution.

**Figure S13.** TE-copy identity number among the 12 aphid species using an upset plot (UpSetR R package, Conway et al., 2017).

**Figure S14.** Distribution of Kimura distances between TE consensus sequence and each TE copies associated or not to chemosensory genes.

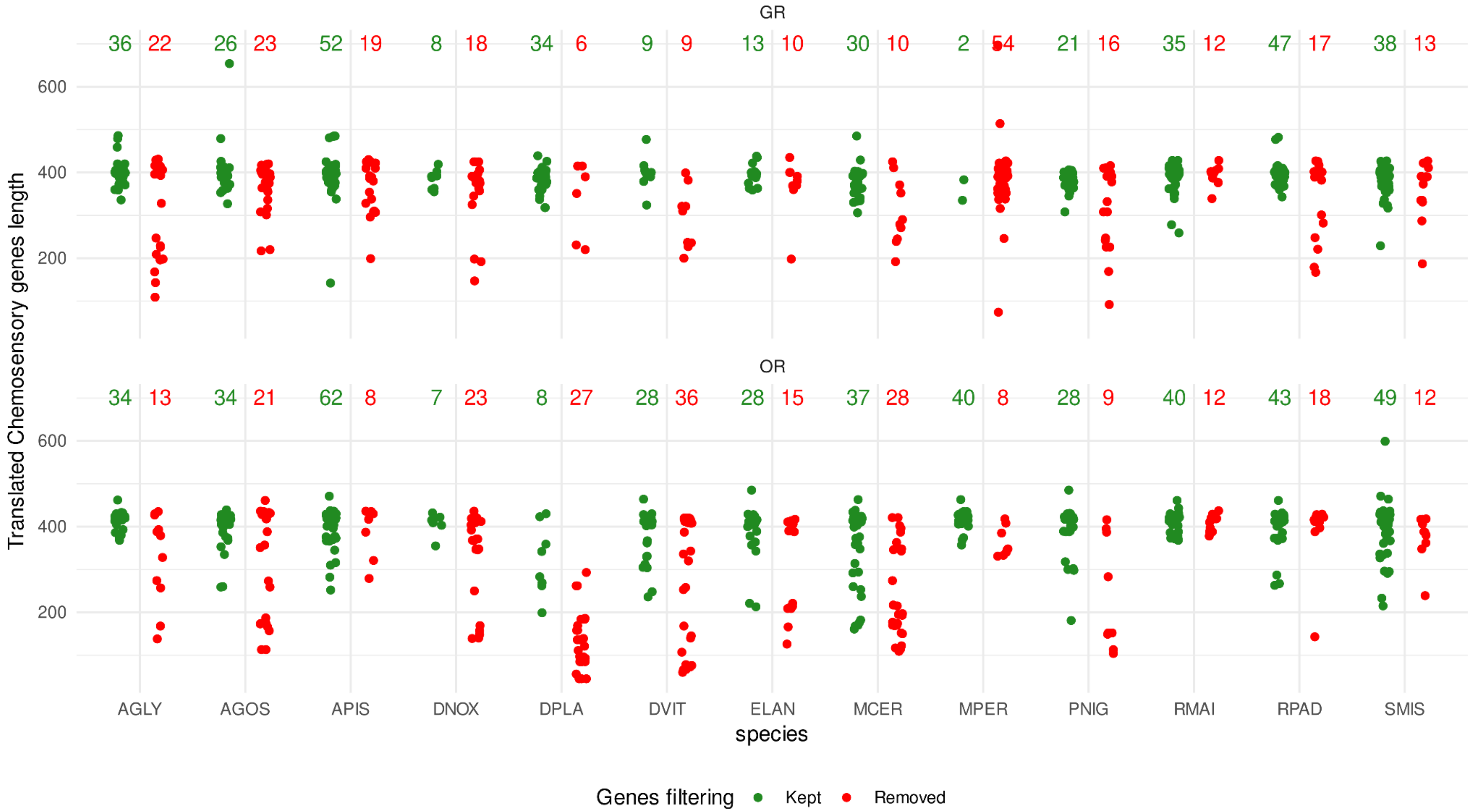

**Figure S7.** Length distribution of annotated GR (upper) and OR (lower) genes for each aphid species studied. Sequences <150pb and with a stop codon, were removed and kept for subsequent clustering analysis and subfamilies detection. Number of genes kept before (green) and after filtering (red) are shown above each distribution.

**Table S11.** Number of annotated GR and OR genes before and after filtering for their length and stop-codon presence for each aphid species to then infer the OR and GR subfamilies.

| Species | OR genes, before filtering | OR genes, after filtering | GR genes, before filtering | GR genes, after filtering |
|---|---|---|---|---|
| AGLY | 47 | 34 | 58 | 36 |
| AGOS | 55 | 34 | 49 | 26 |
| APIS | 70 | 62 | 71 | 52 |
| DNOX | 30 | 7 | 26 | 8 |
| DPLA | 35 | 8 | 40 | 34 |
| DVIT | 64 | 28 | 18 | 9 |
| ELAN | 43 | 28 | 23 | 13 |
| MCER | 65 | 37 | 40 | 30 |
| MPER | 48 | 40 | 56 | 42 |
| PNIG | 37 | 28 | 37 | 21 |
| RMAI | 52 | 40 | 47 | 35 |
| RPAD | 61 | 43 | 64 | 47 |
| SMIS | 61 | 49 | 51 | 38 |
| **TOTAL** | **667** | **365** | **579** | **381** |

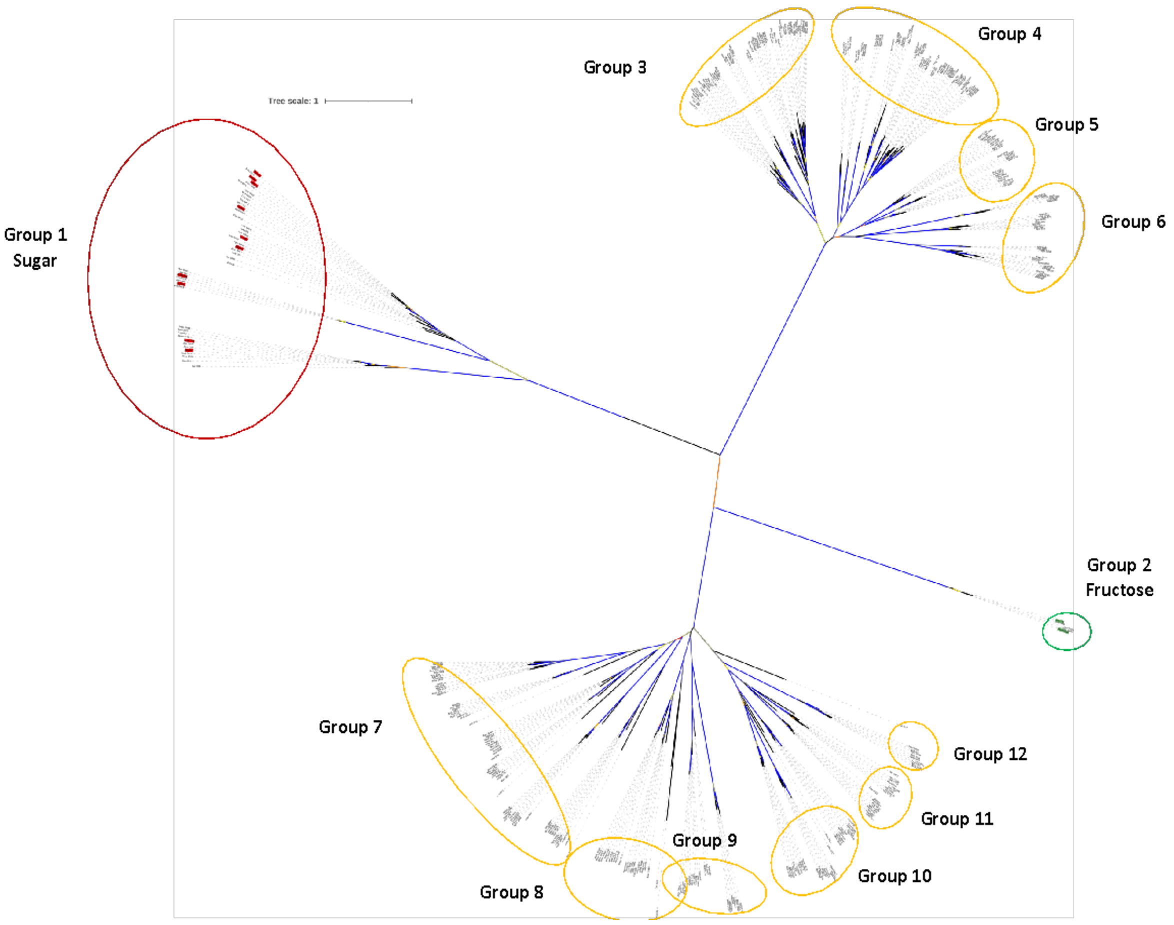

**Figure S8.** Twelve groups of the candidate gustatory receptors (GR) genes were detected using a phylogenetic tree built with IQ-TREE v2.0 using 373 amino-acid sequences. The sugar and fructose gustatory receptors genes were represented in red and green circles, respectively. Bootstrap values were represented in the branches from the lowest 80% in red to the highest 100% in blue.

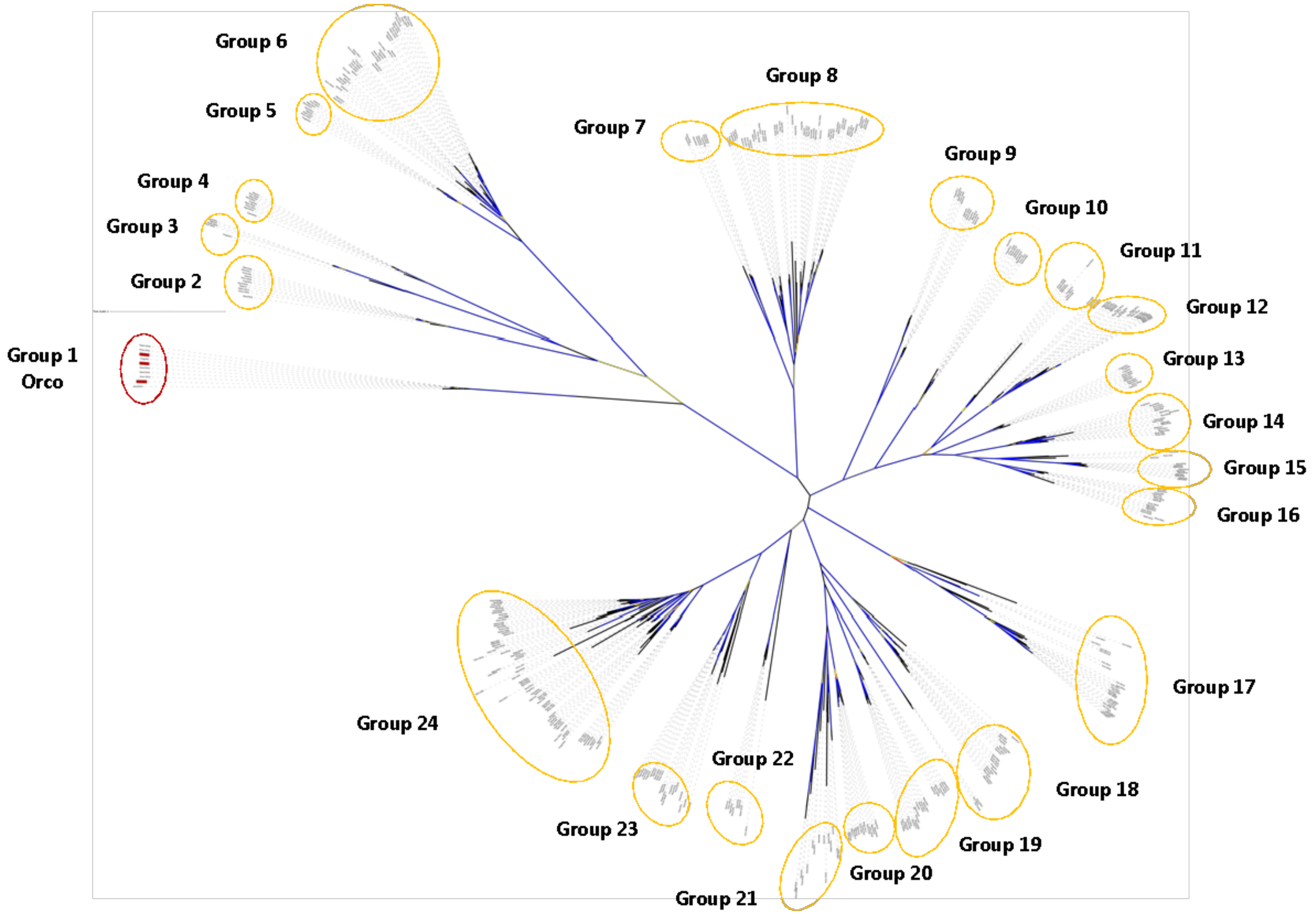

**Figure S9.** Twenty-four groups of the candidate olfactory receptors (OR) genes were detected using a phylogenetic tree built with IQ-TREE v2.0 using 393 amino-acid sequences. The Orco olfactory receptors are represented in red color. Bootstrap values were represented in the branches from the lowest 80% in red to the highest 100% in blue.

**Table S12**. Summary statistics for branch-specific evolutionary models of olfactory receptor (OR) and gustatory receptor (GR) subfamilies, conducted using PAML v4 (Yang, 2007).

| | **GR1 Sugar** | **GR2 Fructose** | **GR3** | **GR4** | **GR5** | **GR6** | **GR7** | **GR8** | **GR9** | **GR10** | **GR11** | **GR12** |
|---|---|---|---|---|---|---|---|---|---|---|---|---|
| ***N*** | 44 | 5 | 56 | 57 | 25 | 32 | 67 | 24 | 22 | 32 | 19 | 10 |
| ***S*** | 354 | 285 | 234 | 408 | 519 | 456 | 441 | 822 | 441 | 639 | 858 | 789 |
| ***ω*** | **0.56463** | **0.88726** | **1.02950** | **0.72983** | **0.95776** | **0.87368** | **0.75979** | **0.79087** | **1.28869** | **0.93441** | **0.48488** | **0.56346** |
| ***$H_o$ln L*** | - 5231.688326 | -1216.789547 | -5789.109381 | - 9980.164042 | -5478.870866 | -6048.019804 | -12088.855011 | -9355.605423 | - 4445.296008 | -9732.432759 | -8102.421100 | -4029.935943 |
| ***$H_a$ln L*** | - 5185.375442 | -1213.859064 | -5738.300335 | - 9919.114544 | -5457.974317 | -6023.154680 | -12017.378990 | -9305.779103 | - 4418.048900 | -9701.739763 | -8074.971065 | -4023.319379 |
| ***2ΔL*** | 92.62577 | 5.860966 | 101.62 | 122.1 | 41.793 | 49.73 | 142.95 | 99.653 | 54.49 | 61.386 | 54.9 | 13.233 |
| ***K*** | 43 | 4 | 55 | 56 | 24 | 31 | 66 | 23 | 21 | 31 | 18 | 9 |
| ***alpha*** | 0.050 | 0.050 | 0.050 | 0.050 | 0.050 | 0.050 | 0.050 | 0.050 | 0.050 | 0.050 | 0.050 | 0.050 |
| ***p*** | 1.702e-05 | **0.2098** | 0.0001335 | 8.08e-07 | 0.01361 | 0.01783 | 1.321e-07 | 1.616e-11 | 8.374e-05 | 0.0009238 | 1.327e-05 | 0.1523 |
| ***$\chi^2$*** | 59.304 | 9.4877 | 73.311 | 74.468 | 36.415 | 44.985 | 85.965 | 35.172 | 32.671 | 44.985 | 28.869 | 16.919 |

***N*= number of genes*; S* = number of sites*; ω= dN/dS* value*; Ho* (InL) : One ratio hypothesis likelihood score; *Ha* (InL) : Free ratio hypothesis likelihood score; *K* :Degrees of freedom*; alpha : 0.05 ; p-value : <0.05 ;* $\chi^2$ : chi-square distribution; GR = Gustatory receptor group**

**Table S12 continuation.** Summary statistics for branch-specific evolutionary models of olfactory receptor (OR) and gustatory receptor (GR) subfamilies, conducted using PAML v4 (Yang, 2007).

| | OR1 Orco | OR2 | OR3 | OR4 | OR5 | OR6 | OR7 | OR8a | OR8b | OR9 | OR10 | OR11 | OR12 |
|---|---|---|---|---|---|---|---|---|---|---|---|---|---|
| ***N*** | 10 | 11 | 5 | 9 | 9 | 37 | 8 | 15 | 31 | 12 | 10 | 8 | 25 |
| ***S*** | 270 | 1128 | 1104 | 795 | 795 | 312 | 1002 | 444 | 60 | 939 | 1011 | 579 | 468 |
| ***ω*** | **0.94407** | **0.67866** | **0.43454** | **1.24689** | **1.50124** | **0.84971** | **0.68829** | **1.47700** | **1.87058** | **1.73311** | **1.03558** | **0.99001** | **1.20879** |
| ***$H_o$lnL*** | -1268.682720 | -4855.073809 | -3317.411588 | -3848.094699 | -3564.354753 | -3407.882779 | -4468.344612 | -3627.976230 | -856.474042 | -5318.031193 | -3850.311741 | -2047.397742 | -3586.931783 |
| ***$H_a$lnL*** | -1259.470606 | -4839.687812 | -3312.508079 | -3842.027866 | -3557.624712 | -3362.358362 | -4459.555754 | -3616.782859 | -834.696518 | -5313.399292 | -3844.135355 | -2043.921807 | -3569.650301 |
| **2ΔL** | 18.424 | 30.772 | 9.807 | 12.134 | 13.46 | 91.049 | 17.578 | 22.387 | 43.555 | 9.2638 | 12.353 | 6.9519 | 34.563 |
| ***K*** | 9 | 10 | 4 | 8 | 8 | 36 | 7 | 14 | 30 | 11 | 9 | 7 | 24 |
| ***alpha*** | 0.050 | 0.050 | 0.050 | 0.050 | 0.050 | 0.050 | 0.050 | 0.050 | 0.050 | 0.050 | 0.050 | 0.050 | 0.050 |
| ***p*** | 0.03056 | 0.0006398 | 0.04381 | 0.1453 | 0.09697 | 1.156e-06 | 0.01403 | 0.07101 | 0.05231 | 0.5976 | 0.1941 | 0.4339 | 0.07515 |
| ***$\chi^2$*** | 16.919 | 18.307 | 9.4877 | 15.507 | 15.507 | 50.998 | 14.067 | 23.685 | 43.773 | 19.675 | 16.919 | 14.067 | 36.415 |

***N***= number of genes; ***S*** = number of sites; ***ω= dN/dS*** value; ***Ho*** (lnL) : One ratio hypothesis likelihood score; ***Ha*** (lnL) : Free ratio hypothesis likelihood score; ***K*** :Degrees of freedom; ***alpha* : 0.05 ; *p-value* : <0.05 ; $\chi^2$** : chi-square distribution; OR = Olfactory receptor group

**Table S12 continuation**. Summary statistics for branch-specific evolutionary models of olfactory receptor (OR) and gustatory receptor (GR) subfamilies, conducted using PAML v4 (Yang, 2007).

| | OR13 | OR14 | OR15 | OR16 | OR17 | OR18 | OR19 | OR20 | OR21 | OR22 | OR23 |
|---|---|---|---|---|---|---|---|---|---|---|---|
| ***N*** | 11 | 17 | 14 | 15 | 25 | 21 | 21 | 14 | 15 | 8 | 22 |
| ***S*** | 885 | 306 | 678 | 378 | 387 | 702 | 1062 | 1086 | 195 | 618 | 300 |
| ***ω*** | **0.44286** | **1.29363** | **0.84264** | **0.77929** | **0.50600** | **0.92750** | **0.45569** | **0.76303** | **0.87819** | **1.11350** | **1.35136** |
| ***$H_o$lnL*** | -3745.413397 | -2194.749421 | -5204.634667 | -2238.207750 | -4932.915224 | -8533.306542 | -8986.854202 | -6162.496007 | -2123.429093 | -3328.270610 | -2677.570631 |
| ***$H_a$lnL*** | -3704.800841 | -2179.939307 | -5187.516447 | -2227.199444 | -4898.485935 | -8510.967430 | -8939.307333 | -6147.827618 | -2109.404275 | -3320.568051 | -2656.982334 |
| ***2ΔL*** | 81.225 | 29.62 | 34.236 | 22.017 | 68.859 | 44.678 | 95.094 | 29.337 | 28.05 | 15.405 | 41.177 |
| ***K*** | 10 | 16 | 13 | 14 | 24 | 20 | 20 | 13 | 14 | 7 | 21 |
| ***alpha*** | 0.050 | 0.050 | 0.050 | 0.050 | 0.050 | 0.050 | 0.050 | 0.050 | 0.050 | 0.050 | 0.050 |
| ***p*** | 2.887e-13 | 0.02007 | 0.001108 | 0.07827 | 3.251e-06 | 0.00122 | 9.411e-12 | 0.005862 | 0.01401 | 0.03114 | 0.005332 |
| ***$\chi^2$*** | 18.307 | 26.296 | 22.362 | 23.685 | 36.415 | 31.41 | 31.41 | 22.362 | 23.685 | 14.067 | 32.671 |

***N*= number of genes*; S* = number of sites*; ω= dN/dS* value*; Ho* (InL) : One ratio hypothesis likelihood score; *Ha* (InL) : Free ratio hypothesis likelihood score; *K* :Degrees of freedom*; alpha : 0.05 ; p-value : <0.05 ;* $\chi^2$ : chi-square distribution; OR = Olfactory receptor group**

**Table S13**. Number of gustatory receptors (GR) and olfactory receptors (OR) subfamilies under positive ($\omega$>0), purifying ($\omega$<0), and neutral ($\omega$=0) selection based on the branch model analysis PAML v4 (Yang, 2007).

| Gene family | Gene subfamily | Number of genes | Sites | Free Omega | Number of sequences with $\omega$>0 | Number of sequences with $\omega$<0 | Number of sequences with $\omega$=0 | Saturated |
|---|---|---|---|---|---|---|---|---|
| GR | GR01 | 44 | 354 | Yes | 1 | 29 | 14 | 0 |
| GR | GR02 | 5 | 285 | No | 0 | 4 | 1 | 0 |
| GR | GR03 | 56 | 234 | Yes | 15 | 22 | 19 | 0 |
| GR | GR04 | 57 | 408 | Yes | 12 | 34 | 11 | 0 |
| GR | GR05 | 25 | 519 | Yes | 5 | 14 | 6 | 0 |
| GR | GR06 | 32 | 456 | Yes | 10 | 16 | 6 | 0 |
| GR | GR07 | 67 | 441 | Yes | 14 | 40 | 13 | 0 |
| GR | GR08 | 24 | 822 | Yes | 5 | 17 | 2 | 0 |
| GR | GR09 | 22 | 441 | Yes | 10 | 6 | 6 | 0 |
| GR | GR10 | 32 | 639 | Yes | 12 | 17 | 3 | 0 |
| GR | GR11 | 19 | 858 | Yes | 0 | 19 | 0 | 0 |
| GR | GR12 | 10 | 789 | No | 1 | 9 | 0 | 0 |
| | Total | 393 | 6246 | | 85 | 227 | 81 | 0 |
| **Gene family** | **Gene subfamily** | **Number of genes** | **Sites** | **Free Omega** | **Number of sequences with $\omega$>0** | **Number of sequences with $\omega$<0** | **Number of sequences with $\omega$=0** | **Saturated** |
| OR | OR01 (Orco) | 10 | 270 | Yes | 1 | 4 | 3 | |
| OR | OR02 | 11 | 1128 | Yes | 2 | 8 | 1 | 0 |
| OR | OR03 | 5 | 1104 | Yes | 1 | 4 | 0 | 0 |

| | | | | | | | | |
|---|---|---|---|---|---|---|---|---|
| OR | OR04 | 9 | 795 | No | 7 | 0 | 2 | 0 |
| OR | OR05 | 9 | 795 | Yes | 2 | 3 | 4 | 0 |
| OR | OR06 | 37 | 312 | Yes | 7 | 19 | 11 | 0 |
| OR | OR07 | 8 | 1002 | Yes | 1 | 4 | 3 | 0 |
| OR | OR08a | 15 | 444 | No | 6 | 3 | 5 | 1 |
| OR | OR08b | 31 | 60 | No | 1 | 3 | 27 | 0 |
| OR | OR09 | 12 | 939 | No | 7 | 1 | 4 | 0 |
| OR | OR10 | 10 | 1011 | No | 2 | 5 | 3 | 0 |
| OR | OR11 | 8 | 579 | No | 2 | 4 | 2 | 0 |
| OR | OR12 | 25 | 468 | No | 8 | 11 | 6 | 0 |
| OR | OR13 | 11 | 885 | Yes | 1 | 9 | 1 | 0 |
| OR | OR14 | 17 | 306 | Yes | 4 | 10 | 3 | 0 |
| OR | OR15 | 14 | 678 | Yes | 4 | 10 | 0 | 0 |
| OR | OR16 | 15 | 378 | No | 6 | 6 | 2 | 1 |
| OR | OR17 | 25 | 387 | Yes | 3 | 15 | 7 | 0 |
| OR | OR18 | 21 | 702 | Yes | 10 | 10 | 1 | 0 |
| OR | OR19 | 21 | 1062 | Yes | 2 | 18 | 1 | 0 |
| OR | OR20 | 14 | 1086 | Yes | 2 | 12 | 0 | 0 |
| OR | OR21 | 15 | 195 | Yes | 4 | 8 | 3 | 0 |
| OR | OR22 | 8 | 618 | Yes | 3 | 3 | 2 | 0 |
| OR | OR23 | 22 | 300 | Yes | 9 | 2 | 11 | 0 |
| | Total | 373 | 15504 | | 95 | 172 | 102 | 2 |

**Table S14**. Summary statistics from site-specific models of evolution for olfactory receptor (OR) and gustatory receptor (GR) subfamilies, inferred using PAML v4 (Yang, 2007).

**Table S14**. **For GR subfamilies.**

| Model | | GR1 Sugar | GR2 Fructose | GR 3 | GR4 | GR5 | GR6 | GR7 | GR8 | GR9 | GR10 | GR11 | GR12 |
|---|---|---|---|---|---|---|---|---|---|---|---|---|---|
| | ***N*** | 44 | 5 | 56 | 57 | 25 | 32 | 67 | 24 | 22 | 32 | 19 | 10 |
| ***M0*** | ***S*** | 354 | 285 | 234 | 408 | 519 | 456 | 441 | 822 | 441 | 639 | 858 | 789 |
| | ***ω*** | 0.56979 | 0.88676 | 1.03516 | 0.76955 | 0.94636 | 0.89746 | 0.77104 | 0.80583 | 1.29361 | 0.93989 | 0.48382 | 0.57127 |
| | ***lnL*** | -5231.825340 | -1216.789767 | -5789.152113 | -9998.205966 | -5479.083036 | -6049.200897 | -12089.821784 | -9356.298793 | -4445.435595 | -9732.621983 | -8102.433935 | -4030.348531 |
| | ***p0*** | 0.83458 | 0.30857 | 0.83073 | 0.86967 | 0.76096 | 0.80609 | 0.86730 | 0.69101 | 0.61788 | 0.75584 | 0.82457 | 0.69644 |
| ***M1a*** | ***p1*** | 0.16542 | 0.69143 | 0.16927 | 0.13033 | 0.23904 | 0.19391 | 0.13270 | 0.30899 | 0.38212 | 0.24416 | 0.17543 | 0.30356 |
| | ***lnL*** | -5077.671870 | -1212.480402 | -5490.213977 | -9402.039709 | -5353.305732 | -5767.175345 | -11420.967944 | -9101.153882 | -4366.772067 | -9296.978281 | -7794.902158 | -3961.899495 |
| **Model** | | **GR1 Sugar** | **GR2 Fructose** | **GR 3** | **GR4** | **GR5** | **GR6** | **GR7** | **GR8** | **GR9** | **GR10** | **GR11** | **GR12** |
| | $p_0$ | 0.81467 | 0.58851 | 0.76326 | 0.82639 | 0.65602 | 0.75180 | 0.82921 | 0.62212 | 0.58002 | 0.69137 | 0.81508 | 0.76591 |
| | $p_1$ | 0.18122 | 0.00000 | 0.21946 | 0.15170 | 0.30452 | 0.20453 | 0.14031 | 0.33966 | 0.27455 | 0.25551 | 0.16766 | 0.15610 |
| **M2a** | $p_2$ | 0.00410 | 0.41149 | 0.01729 | 0.02191 | 0.03946 | 0.04366 | 0.03048 | 0.03822 | 0.14543 | 0.05312 | 0.01726 | 0.07799 |
| | $\omega_2$ | 3.97984 | 1.51071 | 3.76954 | 2.85535 | 3.96942 | 2.87054 | 2.34407 | 3.04424 | 2.82709 | 2.67130 | 2.65679 | 2.12447 |
| | ***lnL*** | -5059.921483 | -1211.381107 | -5397.045087 | -9304.423044 | -5289.395383 | -5713.611109 | -11342.326713 | -9053.610777 | -4315.948000 | -9222.323267 | -7772.779594 | -3953.982877 |
| | $p_0$ | 0.58332 | 0.58851 | 0.76552 | 0.80958 | 0.73032 | 0.78935 | 0.76004 | 0.53892 | 0.62849 | 0.68595 | 0.76264 | 0.80718 |
| **M3** | $p_1$ | 0.35161 | 0.13611 | 0.21827 | 0.16392 | 0.23582 | 0.19702 | 0.17272 | 0.36640 | 0.26443 | 0.25517 | 0.19963 | 0.18940 |
| | $p_2$ | 0.06507 | 0.27538 | 0.01622 | 0.02650 | 0.03386 | 0.01363 | 0.06724 | 0.09467 | 0.10708 | 0.05887 | 0.03773 | 0.00341 |

| | | | | | | | | | | | | | |
|---|---|---|---|---|---|---|---|---|---|---|---|---|---|
| | $\omega_0$ | 0.04657 | 0.29695 | 0.08337 | 0.06080 | 0.26344 | 0.09879 | 0.04973 | 0.02481 | 0.08967 | 0.03038 | 0.04025 | 0.15817 |
| | $\omega_1$ | 0.45267 | 1.51071 | 1.13408 | 0.79733 | 1.21337 | 1.38720 | 0.58398 | 0.64228 | 1.30665 | 0.94832 | 0.70367 | 1.52521 |
| | $\omega_2$ | 1.72582 | 1.51072 | 4.16281 | 2.50663 | 4.27373 | 4.26718 | 1.78145 | 2.16544 | 3.07725 | 2.56541 | 2.13370 | 4.41213 |
| | ***lnL*** | -5052.892647 | -1211.381107 | -5396.002334 | -9300.606905 | -5289.011093 | -5710.513088 | -11335.539722 | -9049.122334 | -4315.713162 | -9222.209445 | -7770.677152 | -3953.943748 |
| **Model** | | **GR1 Sugar** | **GR2 Fructose** | **GR 3** | **GR4** | **GR5** | **GR6** | **GR7** | **GR8** | **GR9** | **GR10** | **GR11** | **GR12** |
| **M7** | ***p*** | 0.17455 | 0.03469 | 0.05396 | 0.04292 | 0.14846 | 0.01359 | 0.04789 | 0.09230 | 0.01421 | 0.01600 | 0.07647 | 0.11235 |
| | ***q*** | 0.56922 | 0.01280 | 0.27935 | 0.26699 | 0.28807 | 0.04376 | 0.29159 | 0.19519 | 0.01886 | 0.05103 | 0.29329 | 0.18386 |
| | ***lnL*** | -5077.945793 | -1212.499290 | -5480.009432 | -9379.699699 | -5365.565151 | -5773.318298 | -11401.784294 | -9100.108902 | -4367.195585 | -9293.315424 | -7804.778381 | -3966.461479 |
| **M8** | $p_0$ | 0.95375 | 0.59059 | 0.98039 | 0.97635 | 0.95253 | 0.92994 | 0.95670 | 0.94099 | 0.84540 | 0.93706 | 0.96119 | 0.86209 |
| | $p_1$ | 0.04625 | 0.40941 | 0.01961 | 0.02365 | 0.04747 | 0.07006 | 0.04330 | 0.05901 | 0.15460 | 0.06294 | 0.03881 | 0.13791 |
| | ***p*** | 0.43013 | 42.38437 | 0.01815 | 0.10074 | 0.46429 | 0.15777 | 0.12647 | 0.18259 | 0.12182 | 0.07722 | 0.17162 | 1.22942 |
| | ***q*** | 1.49525 | 99.00000 | 0.05848 | 0.50118 | 0.60192 | 0.53504 | 0.63448 | 0.38289 | 0.22543 | 0.22473 | 0.79586 | 4.67910 |
| | $\omega$ | 1.89569 | 1.51316 | 3.58829 | 2.50414 | 3.69320 | 2.51872 | 1.97278 | 2.47927 | 2.78821 | 2.42426 | 2.09829 | 1.83933 |
| | ***lnL*** | -5053.994057 | -1211.382858 | -5394.753409 | -9290.460615 | -5290.583339 | -5716.276926 | -11327.505439 | -9048.607034 | -4316.436398 | -9221.616983 | -7771.030353 | -3954.014079 |

**Table S14** continuation. **For OR subfamilies.**

| | | **OR1 Orco** | **OR2** | **OR3** | **OR4** | **OR5** | **OR6** | **OR7** | **OR8a** | **OR8b** | **OR9** | **OR10** | **OR11** |
|---|---|---|---|---|---|---|---|---|---|---|---|---|---|
| ***M0*** | ***N*** | 10 | 11 | 5 | 9 | 9 | 37 | 8 | 15 | 31 | 12 | 10 | 8 |
| | ***S*** | 270 | 1128 | 1104 | 795 | 795 | 312 | 1002 | 444 | 60 | 939 | 1011 | 579 |
| | ***ω*** | 0.71014 | 0.67310 | 0.47414 | 1.24881 | 1.39432 | 0.85984 | 0.75226 | 1.50944 | 1.87082 | 1.81900 | 0.94383 | 0.98620 |
| | ***lnL*** | -1277.419980 | -4855.183909 | -3324.042385 | -3848.105608 | -3568.098925 | -3408.209158 | -4479.693944 | -3631.526679 | -856.474056 | -5319.972747 | -3857.677939 | -2047.402217 |
| ***M1a*** | ***$p_0$*** | 0.36814 | 0.55993 | 0.71671 | 0.34264 | 0.47116 | 0.76251 | 0.60290 | 0.57692 | 0.72571 | 0.52422 | 0.44514 | 0.39714 |
| | ***$p_1$*** | 0.63186 | 0.44007 | 0.28329 | 0.65736 | 0.52884 | 0.23749 | 0.39710 | 0.42308 | 0.27429 | 0.47578 | 0.55486 | 0.60286 |
| | ***lnL*** | -1259.699478 | -4797.822208 | -3308.891909 | -3832.394693 | -3543.684662 | -3296.34482 | -4396.215633 | -3570.720139 | -827.407331 | -5267.011477 | -3823.483550 | -2037.472383 |
| | | **OR1 Orco** | **OR2** | **OR3** | **OR4** | **OR5** | **OR6** | **OR7** | **OR8a** | **OR8b** | **OR9** | **OR10** | **OR11** |
| ***M2a*** | ***$p_0$*** | 0.21281 | 0.53032 | 0.71671 | 0.67958 | 0.30517 | 0.71475 | 0.65679 | 0.53348 | 0.50886 | 0.27401 | 0.38696 | 0.63895 |
| | ***$p_1$*** | 0.74794 | 0.46054 | 0.15964 | 0.12322 | 0.61663 | 0.24850 | 0.26918 | 0.25713 | 0.41403 | 0.60330 | 0.54882 | 0.18189 |
| | ***$p_2$*** | 0.03925 | 0.00914 | 0.12365 | 0.19720 | 0.07820 | 0.03676 | 0.07403 | 0.20939 | 0.07711 | 0.12270 | 0.06423 | 0.17916 |
| | ***$ω_2$*** | 7.15980 | 5.78168 | 1.00000 | 3.11829 | 6.07725 | 2.77295 | 2.72780 | 2.98276 | 6.40163 | 5.40352 | 4.57668 | 2.77583 |
| | ***lnL*** | -1251.776345 | -4781.144351 | -3308.891909 | -3801.445999 | -3497.405393 | -3276.383988 | -4382.104041 | -3522.928259 | -786.311979 | -5179.266618 | -3799.778882 | -2031.484327 |
| ***M3*** | ***$p_0$*** | 0.32170 | 0.71351 | 0.31804 | 0.05548 | 0.72186 | 0.74631 | 0.74238 | 0.59648 | 0.51417 | 0.41306 | 0.65438 | 0.19889 |
| | ***$p_1$*** | 0.64604 | 0.28469 | 0.67990 | 0.74503 | 0.27074 | 0.23425 | 0.24481 | 0.29658 | 0.40955 | 0.48803 | 0.32655 | 0.62448 |
| | ***$p_2$*** | 0.03227 | 0.00180 | 0.00206 | 0.19949 | 0.00740 | 0.01944 | 0.01282 | 0.10694 | 0.07628 | 0.09891 | 0.01907 | 0.17663 |
| | ***$ω_0$*** | 0.25649 | 0.12934 | 0.00000 | 0.00000 | 0.31501 | 0.08512 | 0.13582 | 0.10659 | 0.00000 | 0.11346 | 0.19958 | 0.00000 |
| | ***$ω_1$*** | 1.25267 | 1.51353 | 0.60434 | 0.58481 | 3.03382 | 1.27091 | 1.55645 | 1.63950 | 1.04134 | 1.36181 | 1.80986 | 0.59144 |

| | | | | | | | | | | | | | |
|---|---|---|---|---|---|---|---|---|---|---|---|---|---|
| | $\omega_2$ | 8.23044 | 10.27914 | 5.54959 | 3.11919 | 14.95929 | 3.40857 | 4.21949 | 3.66459 | 6.52613 | 6.01030 | 6.72778 | 2.81656 |
| | *lnL* | -1251.614549 | -4779.219529 | -3307.756934 | -3801.408614 | -3495.220514 | -3275.335808 | -4382.009169 | -3522.071847 | -786.305067 | -5179.035727 | -3799.432971 | -2031.469449 |
| | | **OR1 Orco** | **OR2** | **OR3** | **OR4** | **OR5** | **OR6** | **OR7** | **OR8a** | **OR8b** | **OR9** | **OR10** | **OR11** |
| ***M7*** | *p* | 0.56952 | 0.00534 | 0.53125 | 0.03523 | 4.36286 | 0.06430 | 0.01469 | 0.01169 | 0.01581 | 0.01886 | 0.00751 | 0.00726 |
| | *q* | 0.22355 | 0.00663 | 0.72315 | 0.01998 | 0.00500 | 0.16543 | 0.01920 | 0.00500 | 0.09509 | 0.01688 | 0.00500 | 0.00500 |
| | *lnL* | -1260.710668 | -4798.331056 | -3308.277304 | -3833.774262 | -3569.585274 | -3303.021239 | -4396.427598 | -3589.963693 | -827.608477 | -5267.344755 | -3823.955115 | -2037.473281 |
| ***M8*** | $p_0$ | 0.95789 | 0.99135 | 0.99906 | 0.80324 | 0.92481 | 0.94742 | 0.90001 | 0.77658 | 0.92524 | 0.87820 | 0.93845 | 0.82429 |
| | $p_1$ | 0.04211 | 0.00865 | 0.00094 | 0.19676 | 0.07519 | 0.05258 | 0.09999 | 0.22342 | 0.07476 | 0.12180 | 0.06155 | 0.17571 |
| | *p* | 0.45550 | 0.00516 | 0.59207 | 3.49022 | 0.01159 | 0.13305 | 0.27595 | 0.15528 | 0.01774 | 0.01159 | 0.00739 | 0.97790 |
| | *q* | 0.11128 | 0.00502 | 0.82021 | 2.84556 | 0.00500 | 0.32512 | 0.60179 | 0.28619 | 0.02063 | 0.00500 | 0.00500 | 1.17869 |
| | $\omega$ | 6.96738 | 6.08225 | 6.10592 | 3.12781 | 6.31042 | 2.55244 | 2.51752 | 2.94391 | 6.46621 | 5.46641 | 4.69363 | 2.80853 |
| | *lnL* | -1251.866492 | -4781.370179 | -3308.148161 | -3801.432365 | -3497.460429 | -3279.169203 | -4382.169578 | -3523.414754 | -786.377213 | -5179.281417 | -3799.798015 | -2031.476879 |
| | | **OR12** | **OR13** | **OR14** | **OR15** | **OR16** | **OR17** | **OR18** | **OR19** | **OR20** | **OR21** | **OR22** | **OR23** |
| **M0** | ***N*** | 25 | 11 | 17 | 14 | 15 | 25 | 21 | 21 | 14 | 15 | 8 | 22 |
| | ***S*** | 468 | 885 | 306 | 678 | 378 | 387 | 702 | 1062 | 1086 | 195 | 618 | 300 |
| | $\omega$ | 1.16038 | 0.44317 | 1.29761 | 0.93656 | 0.84268 | 0.53613 | 1.04444 | 0.47798 | 0.75217 | 0.92974 | 1.13172 | 1.29806 |
| | *lnL* | -3588.139836 | -3745.413691 | -2194.753612 | -5214.486356 | -2242.826138 | -4940.105104 | -8552.735563 | -8993.558702 | -6162.761013 | -2125.005660 | -3330.081615 | -2678.888227 |
| **M1a** | $p_0$ | 0.68868 | 0.79216 | 0.59661 | 0.70533 | 0.69271 | 0.83371 | 0.75104 | 0.81679 | 0.67477 | 0.73123 | 0.38003 | 0.67300 |
| | $p_1$ | 0.31132 | 0.20784 | 0.40339 | 0.29467 | 0.30729 | 0.16629 | 0.24896 | 0.18321 | 0.32523 | 0.26877 | 0.61997 | 0.32700 |
| | *lnL* | -3486.586634 | -3681.069046 | -2158.354702 | -5063.053386 | -2186.389432 | -4711.510404 | -8191.950929 | -8678.778617 | -6020.326260 | -2036.359161 | -3304.977694 | -2617.608997 |
| | | **OR12** | **OR13** | **OR14** | **OR15** | **OR16** | **OR17** | **OR18** | **OR19** | **OR20** | **OR21** | **OR22** | **OR23** |
| **M2a** | $p_0$ | 0.59234 | 0.83799 | 0.42401 | 0.64662 | 0.59354 | 0.82025 | 0.64308 | 0.80504 | 0.61277 | 0.64163 | 0.31707 | 0.53927 |

| | | | | | | | | | | | | | |
|---|---|---|---|---|---|---|---|---|---|---|---|---|---|
| | ***p₁*** | 0.31969 | 0.14657 | 0.51653 | 0.29392 | 0.38315 | 0.17491 | 0.30966 | 0.18309 | 0.36564 | 0.33663 | 0.61026 | 0.36370 |
| | ***p₂*** | 0.08797 | 0.01544 | 0.05945 | 0.05946 | 0.02331 | 0.00484 | 0.04725 | 0.01187 | 0.02159 | 0.02174 | 0.07267 | 0.09703 |
| | ***ω₂*** | 3.77467 | 4.40582 | 4.43938 | 3.07580 | 5.99272 | 3.49845 | 3.49847 | 2.79929 | 4.95223 | 3.60716 | 3.85225 | 3.97549 |
| | ***lnL*** | -3438.453620 | -3662.868100 | -2131.706907 | -5031.428120 | -2156.215036 | -4701.791333 | -8097.582281 | -8663.562640 | -5963.486385 | -2021.667637 | -3284.332897 | -2574.028379 |
| **M3** | ***p₀*** | 0.67486 | 0.58639 | 0.58207 | 0.65284 | 0.52051 | 0.64560 | 0.63140 | 0.73960 | 0.73515 | 0.64471 | 0.37781 | 0.60685 |
| | ***p₁*** | 0.26661 | 0.39277 | 0.39502 | 0.29005 | 0.45173 | 0.27770 | 0.31566 | 0.23038 | 0.25744 | 0.33664 | 0.57315 | 0.31410 |
| | ***p₂*** | 0.05853 | 0.02084 | 0.02291 | 0.05711 | 0.02776 | 0.07670 | 0.05294 | 0.03002 | 0.00741 | 0.01865 | 0.04904 | 0.07905 |
| | ***ω₀*** | 0.04735 | 0.04683 | 0.10372 | 0.10828 | 0.00000 | 0.00435 | 0.04540 | 0.03742 | 0.07173 | 0.00000 | 0.04229 | 0.13042 |
| | ***ω₁*** | 1.45850 | 0.54834 | 1.61123 | 1.02520 | 0.82450 | 0.39270 | 0.91872 | 0.67852 | 1.48937 | 1.07150 | 1.17034 | 1.28588 |
| | ***ω₂*** | 4.41467 | 3.99716 | 6.64091 | 3.11859 | 5.48638 | 1.38536 | 3.28300 | 2.12257 | 7.27234 | 3.96416 | 4.43243 | 4.34745 |
| | ***lnL*** | -3437.771521 | -3662.608215 | -2130.789706 | -5031.424098 | -2155.939738 | -4695.518331 | -8097.321728 | -8659.143002 | -5961.340929 | -2021.568078 | -3284.228800 | -2573.756787 |
| | | **OR12** | **OR13** | **OR14** | **OR15** | **OR16** | **OR17** | **OR18** | **OR19** | **OR20** | **OR21** | **OR22** | **OR23** |
| **M7** | ***p*** | 0.00945 | 0.06887 | 0.00856 | 0.09476 | 0.01392 | 0.09327 | 0.06380 | 0.09623 | 0.01291 | 0.01452 | 0.02078 | 0.01675 |
| | ***q*** | 0.01886 | 0.17861 | 0.00576 | 0.17918 | 0.02706 | 0.42014 | 0.19649 | 0.36116 | 0.02602 | 0.02855 | 0.01255 | 0.03282 |
| | ***lnL*** | -3486.779859 | -3684.323225 | -2164.038648 | -5071.873297 | -2186.708961 | -4703.931918 | -8190.929871 | -8685.315986 | -6020.805255 | -2034.553460 | -3305.056752 | -2621.680496 |
| **M8** | ***p₀*** | 0.90256 | 0.98037 | 0.93716 | 0.91468 | 0.97500 | 0.98193 | 0.94664 | 0.97281 | 0.97938 | 0.98072 | 0.91736 | 0.89775 |
| | ***p₁*** | 0.09744 | 0.01963 | 0.06284 | 0.08532) | 0.02500 | 0.01807 | 0.05336 | 0.02719 | 0.02062 | 0.01928 | 0.08264 | 0.10225 |
| | ***p*** | 0.01433 | 0.43423 | 0.00597 | 0.29272 | 0.09010 | 0.13751 | 0.13389 | 0.17680 | 0.00500 | 0.01672 | 0.03341 | 0.18363 |
| | ***q*** | 0.02810 | 1.28183 | 0.00500 | 0.58188 | 0.13759 | 0.61739 | 0.29295 | 0.73183 | 0.00758 | 0.03256 | 0.01869 | 0.22021 |
| | ***ω*** | 3.58020 | 4.07223 | 4.23499 | 2.71380 | 5.73082 | 2.05538 | 3.11261 | 2.15466 | 5.15913 | 3.52442 | 3.63801 | 3.93879 |
| | ***lnL*** | -3438.692997 | -3662.634207 | -2131.892309 | -5031.964759 | -2156.051260 | -4693.116033 | -8097.050353 | -8659.506895 | -5963.772860 | -2021.029561 | -3284.481367 | -2574.659541 |

**Table S15**. Likelihood-ratio test results for site-specific evolutionary models of olfactory receptor (OR) and gustatory receptor (GR) subfamilies, conducted using PAML v4 (Yang, 2007).

| Model | | GR1 Sugar | GR2 Fructose | GR 3 | GR4 | GR5 | GR6 | GR7 | GR8 | GR9 | GR10 | GR11 | GR12 |
|---|---|---|---|---|---|---|---|---|---|---|---|---|---|
| | ***lnL*** | -5231.825340 | -1216.789767 | -5789.152113 | -9998.205966 | -5479.083036 | -6049.200897 | -12089.821784 | -9356.298793 | -4445.435595 | -9732.621983 | -8102.433935 | -4030.348531 |
| | ***lnL*** | -5052.892647 | -1211.381107 | -5396.002334 | -9300.606905 | -5289.011093 | -5710.513088 | -11335.539722 | -9049.122334 | -4315.713162 | -9222.209445 | -7770.677152 | -3953.943748 |
| M0 vs M3 | ***2ΔL*** | -357.87 | 10.817 | 786.3 | 1395.2 | 380.14 | 677.38 | 1508.6 | 614.35 | 259.44 | 1020.8 | 663.51 | 152.81 |
| | ***K*** | 4 | 4 | 4 | 4 | 4 | 4 | 4 | 4 | 4 | 4 | 4 | 4 |
| | ***alpha*** | 0.05 | 0.05 | 0.05 | 0.05 | 0.05 | 0.05 | 0.05 | 0.05 | 0.05 | 0.05 | 0.05 | 0.05 |
| | ***p*** | 2.2e-16 | 0.0287 | 2.2e-16 | 2.2e-16 | 2.2e-16 | 2.2e-16 | 2.2e-16 | 2.2e-16 | 2.2e-16 | 2.2e-16 | 2.2e-16 | 2.2e-16 |
| | $\chi^2$ | 9.4877 | 9.4877 | 9.4877 | 9.4877 | 9.4877 | 9.4877 | 9.4877 | 9.4877 | 9.4877 | 9.4877 | 9.4877 | 9.4877 |
| | ***lnL*** | -5077.671870 | -1212.480402 | -5490.213977 | -9402.039709 | -5353.305732 | -5767.175345 | -11420.967944 | -9101.153882 | -4366.772067 | -9296.978281 | -7794.902158 | -3961.899495 |
| | ***lnL*** | -5059.921483 | -1211.381107 | -5397.045087 | -9304.423044 | -5289.395383 | -5713.611109 | -11342.326713 | -9053.610777 | -4315.948000 | -9222.323267 | -7772.779594 | -3953.982877 |
| M1a vs M2a | ***2ΔL*** | 35.501 | 2.1986 | 186.34 | 195.23 | 127.82 | 107.13 | 157.28 | 95.086 | 101.65 | 149.31 | 44.245 | 15.833 |
| | ***K*** | 2 | 2 | 2 | 2 | 2 | 2 | 2 | 2 | 2 | 2 | 2 | 2 |
| | ***alpha*** | 0.05 | 0.05 | 0.05 | 0.05 | 0.05 | 0.05 | 0.05 | 0.05 | 0.05 | 0.05 | 0.05 | 0.05 |
| | ***p*** | 1.954811e-08 | 0.6992874 | 1 | 2.2e-16 | 2.2e-16 | 2.2e-16 | 2.2e-16 | 2.2e-16 | 2.2e-16 | 2.2e-16 | 2.468e-10 | 0.0003646 |
| | $\chi^2$ | 5.9915 | 5.9915 | 5.9915 | 5.9915 | 5.9915 | 5.9915 | 5.9915 | 5.9915 | 5.9915 | 5.9915 | 5.9915 | 5.9915 |

| | | | | | | | | | | | | | |
|---|---|---|---|---|---|---|---|---|---|---|---|---|---|
| | | ***lnL*** | -5077.945793 | -1212.499290 | -5480.009432 | -9379.699699 | -5365.565151 | -5773.318298 | -11401.784294 | -9100.108902 | -4367.195585 | -9293.315424 | -7804.778381 | -3966.461479 |
| | | ***lnL*** | -5053.994057 | -1211.382858 | -5394.753409 | -9290.460615 | -5290.583339 | -5716.276926 | -11327.505439 | -9048.607034 | -4316.436398 | -9221.616983 | -7771.030353 | -3954.014079 |
| | M7 vs M8 | ***2ΔL*** | 47.903 | 2.2329 | 170.51 | 178.48 | 149.96 | 114.08 | 148.56 | 103 | 101.52 | 143.4 | 67.496 | 24.895 |
| | | ***K*** | 2 | 2 | 2 | 2 | 2 | 2 | 2 | 2 | 2 | 2 | 2 | 2 |
| | | ***alpha*** | 0.05 | 0.05 | 0.05 | 0.05 | 0.05 | 0.05 | 0.05 | 0.05 | 0.05 | 0.05 | 0.05 | 0.05 |
| | | ***p*** | 3.961806e-11 | 0.327446 | 9.414119e-38 | 1.753712e-39 | 2.727802e-33 | 1.687512e-25 | 5.509385e-33 | 4.295591e-23 | 9.027452e-23 | 7.273823e-32 | 2.205046e-15 | 3.927922e-06 |
| | | $\chi^2$ | 5.9915 | 5.9915 | 5.9915 | 5.9915 | 5.9915 | 5.9915 | 5.9915 | 5.9915 | 5.9915 | 5.9915 | 5.9915 | 5.9915 |

**Table S15 continuation (for OR subfamilies)**. Likelihood-ratio test results for site-specific evolutionary models of olfactory receptor (OR) and gustatory receptor (GR) subfamilies, conducted using PAML v4 (Yang, 2007)..

| Model | | OR1 Orco | OR2 | OR3 | OR4 | OR5 | OR6 | OR7 | OR8a | OR8b | OR9 | OR10 | OR11 |
|---|---|---|---|---|---|---|---|---|---|---|---|---|---|
| | ***lnL*** | -1277.419980 | -4855.183909 | -3324.042385 | -3848.105608 | -3568.098925 | -3408.209158 | -4479.693944 | -3631.526679 | -856.474056 | -5319.972747 | -3857.677939 | -2047.402217 |
| | ***lnL*** | -1251.614549 | -4779.219529 | -3307.756934 | -3801.408614 | -3495.220514 | -3275.335808 | -4382.009169 | -3522.071847 | -786.305067 | -5179.035727 | -3799.432971 | -2031.469449 |
| | ***2ΔL*** | 51.611 | 151.93 | 32.571 | 93.394 | 145.76 | 265.75 | 195.37 | 218.91 | 140.34 | 281.87 | 116.49 | 31.866 |
| **M0 vs M3** | ***K*** | 4 | 4 | 4 | 4 | 4 | 4 | 4 | 4 | 4 | 4 | 4 | 4 |
| | ***alpha*** | 0.05 | 0.05 | 0.05 | 0.05 | 0.05 | 0.05 | 0.05 | 0.05 | 0.05 | 0.05 | 0.05 | 0.05 |
| | ***p*** | 1.663667e-10 | 2.2e-16 | 1.462175e-06 | 2.2e-16 | 2.2e-16 | 2.2e-16 | 2.2e-16 | 2.2e-16 | 2.2e-16 | 2.2e-16 | 2.2e-16 | 2.038e-06 |
| | $\chi^2$ | 9.4877 | 9.4877 | 9.4877 | 9.4877 | -380.14 | 9.4877 | 9.4877 | 9.4877 | 9.4877 | 9.4877 | 9.4877 | 9.4877 |
| | ***lnL*** | -1259.699478 | -4797.822208 | -3308.891909 | -3832.394693 | -3543.684662 | -3296.34482 | -4396.215633 | -3570.720139 | -827.407331 | -5267.011477 | -3823.483550 | -2037.472383 |
| | ***lnL*** | -1251.776345 | -4781.144351 | -3308.891909 | -3801.445999 | -3497.405393 | -3276.383988 | -4382.104041 | -3522.928259 | -786.311979 | -5179.266618 | -3799.778882 | -2031.484327 |
| | ***2ΔL*** | 15.846 | 33.356 | 0 | 61.897 | 92.559 | 39.922 | 28.223 | 95.584 | 82.191 | 175.49 | 47.409 | 11.976 |
| **M1a vs M2a** | ***K*** | 2 | 2 | 2 | 2 | 2 | 2 | 2 | 2 | 2 | 2 | 2 | 2 |
| | ***alpha*** | 0.05 | 0.05 | 0.05 | 0.05 | 0.05 | 0.05 | 0.05 | 0.05 | 0.05 | 0.05 | 0.05 | 0.05 |
| | ***p*** | 0.0003623 | 5.713e-08 | 1 | 3.624e-14 | 7.964689e-21 | 2.143e-09 | 7.437e-07 | 1.754891e-21 | 1.420743e-18 | 7.814333e-39 | 5.072e-11 | 0.002509 |
| | $\chi^2$ | 5.9915 | 5.9915 | 5.9915 | 5.9915 | 5.9915 | 5.9915 | 5.9915 | 5.9915 | 5.9915 | 5.9915 | 5.9915 | 5.9915 |
| | ***lnL*** | -1260.710668 | -4798.331056 | -3308.277304 | -3833.774262 | -3569.585274 | -3303.021239 | -4396.427598 | -3589.963693 | -827.608477 | -5267.344755 | -3823.955115 | -2037.473281 |
| | ***lnL*** | -1251.866492 | -4781.370179 | -3308.148161 | -3801.432365 | -3497.460429 | -3279.169203 | -4382.169578 | -3523.414754 | -786.377213 | -5179.281417 | -3799.798015 | -2031.476879 |
| | ***2ΔL*** | 17.688 | 33.922 | 0.25829 | 64.684 | 144.25 | 47.704 | 28.516 | 133.1 | 82.463 | 176.13 | 48.314 | 11.993 |
| **M7 vs M8** | ***K*** | 2 | 2 | 2 | 2 | 2 | 2 | 2 | 2 | 2 | 2 | 2 | 2 |
| | ***alpha*** | 0.05 | 0.05 | 0.05 | 0.05 | 0.05 | 0.05 | 0.05 | 0.05 | 0.05 | 0.05 | 0.05 | 0.05 |
| | ***p*** | 0.0001442 | 4.305e-08 | 0.8788 | 8.997e-15 | 2.2e-16 | 4.377e-11 | 6.424e-07 | 2.2e-16 | 2.2e-16 | 2.2e-16 | 3.226e-11 | 0.002488 |
| | $\chi^2$ | 5.9915 | 5.9915 | 5.9915 | 5.9915 | 5.9915 | 5.9915 | 5.9915 5.9915 | 5.9915 | 5.9915 | 5.9915 | 5.9915 | 5.9915 |

**Table S15 continuation (for OR subfamilies)**. Likelihood-ratio test results for site-specific evolutionary models of olfactory receptor (OR) and gustatory receptor (GR) subfamilies, conducted using PAML v4 (Yang, 2007).

| | | OR 12 | OR 13 | OR 14 | OR 15 | OR 16 | OR 17 | OR 18 | OR 19 | OR 20 | OR 21 | OR 22 | OR 23 |
|---|---|---|---|---|---|---|---|---|---|---|---|---|---|
| ***M0 vs M3*** | ***lnL*** | -3588.139836 | -3745.413691 | -2194.753612 | -5214.486356 | -2242.826138 | -4940.105104 | -8552.735563 | -8993.558702 | -6162.761013 | -2125.005660 | -3330.081615 | -2678.888227 |
| | ***lnL*** | -3437.771521 | -3662.608215 | -2130.789706 | -5031.424098 | -2155.939738 | -4695.518331 | -8097.321728 | -8659.143002 | -5961.340929 | -2021.568078 | -3284.228800 | -2573.756787 |
| | ***2ΔL*** | 300.74 | 165.61 | 127.93 | 366.12 | 173.77 | 489.17 | 910.83 | 668.83 | 402.84 | -206.88 | 91.706 | 210.26 |
| | ***K*** | 4 | 4 | 4 | 4 | 4 | 4 | 4 | 4 | 4 | 4 | 4 | 4 |
| | ***alpha*** | 0.05 | 0.05 | 0.05 | 0.05 | 0.05 | 0.05 | 0.05 | 0.05 | 0.05 | 0.05 | 0.05 | 0.05 |
| | ***p*** | 2.2e-16 | 2.2e-16 | 2.2e-16 | 2.2e-16 | 2.2e-16 | 2.2e-16 | 2.2e-16 | 2.2e-16 | 2.2e-16 | 2.2e-16 | 2.2e-16 | 2.2e-16 |
| | $\chi^2$ | 9.4877 | 9.4877 | 9.4877 | 9.4877 | 9.4877 | 9.4877 | 9.4877 | 9.4877 | 9.4877 | 9.4877 | 9.4877 | 9.4877 |
| ***M1a vs M2a*** | ***lnL*** | -3486.586634 | -3681.069046 | -2158.354702 | -5063.053386 | -2186.389432 | -4711.510404 | -8191.950929 | -8678.778617 | -6020.326260 | -2036.359161 | -3304.977694 | -2617.608997 |
| | ***lnL*** | -3438.453620 | -3662.868100 | -2131.706907 | -5031.428120 | -2156.215036 | -4701.791333 | -8097.582281 | -8663.562640 | -5963.486385 | -2021.667637 | -3284.332897 | -2574.028379 |
| | ***2ΔL*** | 96.266 | 36.402 | 53.296 | 63.251 | 60.349 | 19.438 | 188.74 | 30.432 | 113.68 | 29.383 | 41.29 | 87.161 |
| | ***K*** | 2 | 2 | 2 | 2 | 2 | 2 | 2 | 2 | 2 | 2 | 2 | 2 |
| | ***alpha*** | 0.05 | 0.05 | 0.05 | 0.05 | 0.05 | 0.05 | 0.05 | 0.05 | 0.05 | 0.05 | 0.05 | 0.05 |
| | ***p*** | 1.247664e-21 | 1.245746e-08 | 2.673e-12 | 1.842e-14 | 7.86e-14 | 6.013e-05 | 1.038047e-41 | 2.465e-07 | 2.06422e-25 | 4.164e-07 | 1.082e-09 | 1.183527e-19 |
| | $\chi^2$ | 5.9915 | 5.9915 | 5.9915 | 5.9915 | 5.9915 | 5.9915 | 5.9915 | 5.9915 | 5.9915 | 5.9915 | 5.9915 | 5.9915 |
| ***M7 vs M8*** | ***lnL*** | -3486.779859 | -3684.323225 | -2164.038648 | -5071.873297 | -2186.708961 | -4703.931918 | -8190.929871 | -8685.315986 | -6020.805255 | -2034.553460 | -3305.056752 | -2621.680496 |
| | ***lnL*** | -3438.692997 | -3662.634207 | -2131.892309 | -5031.964759 | -2156.051260 | -4693.116033 | -8097.050353 | -8659.506895 | -5963.772860 | -2021.029561 | -3284.481367 | -2574.659541 |
| | ***2ΔL*** | 96.174 | 43.378 | 64.293 | 79.817 | 61.315 | 21.632 | 187.76 | 51.618 | 114.06 | 27.048 | 41.151 | 94.042 |
| | ***K*** | 2 | 2 | 2 | 2 | 2 | 2 | 2 | 2 | 2 | 2 | 2 | 2 |
| | ***alpha*** | 0.05 | 0.05 | 0.05 | 0.05 | 0.05 | 0.05 | 0.05 | 0.05 | 0.05 | 0.05 | 0.05 | 0.05 |
| | ***p*** | 2.2e-16 | 3.807e-10 | 1.094e-14 | 2.2e-16 | 4.848e-14 | 2.008e-05 | 2.2e-16 | 6.184e-12 | 2.2e-16 | 1.339e-06 | 1.159e-09 | 2.2e-16 |

| | | | | | | | | | | | | | |
|---|---|---|---|---|---|---|---|---|---|---|---|---|---|
| | $\chi^2$ | 5.9915 | 5.9915 | 5.9915 | 5.9915 | 5.9915 | 5.9915 | 5.9915 | 5.9915 | 5.9915 | 5.9915 | 5.9915 | 5.9915 |

**Table S16**. Number of gustatory receptors (GR) and olfactory receptors (OR) subfamilies under positive selection (M8 model) based on the site model analysis PAML v4 (Yang, 2007).

| **Genes** | **Number of genes** | **Sites** | **M0 vs M3** | **M1a vs M2a** | **M7 vs M8** | **Positively selected sites*** |
|---|---|---|---|---|---|---|
| GR01 | 44 | 354 | M0 | M1a | M7 | |
| GR02 | 5 | 285 | M0 | M2a | M8 | 1S,2T,3F,4T,11C,13Y,16F,21I,26K,29N,39Q,41T,44S,45K,46C,48K,72G,73F,93L |
| GR03 | 56 | 234 | M0 | M2a | M7 | |
| GR04 | 57 | 408 | M0 | M1a | M7 | |
| GR05 | 25 | 519 | M0 | M1a | M7 | |
| GR06 | 32 | 456 | M0 | M1a | M7 | |
| GR07 | 67 | 441 | M0 | M1a | M7 | |
| GR08 | 24 | 822 | M0 | M1a | M7 | |
| GR09 | 22 | 441 | M0 | M1a | M7 | |
| GR10 | 32 | 639 | M0 | M1a | M7 | |
| GR11 | 19 | 858 | M0 | M1a | M7 | |
| GR12 | 10 | 789 | M0 | M1a | M7 | |
| Total | 393 | 6246 | | | | |

*Positive selection sites estimated under model M8 by Bayes empirical Bayes (BEB) approach with posterior probabilities (PPs) > 95%. The overlapping sites between M8 and M2a are underlined.

**Table S16 continuation (for OR)**. Number of gustatory receptors (GR) and olfactory receptors (OR) subfamilies under positive selection (M8 model) based on the site model analysis PAML v4 (Yang, 2007).

| Genes | Number of genes | Sites | M0 vs M3 | M1a vs M2a | M7 vs M8 | Positively Selected Sites |
|---|---|---|---|---|---|---|
| OR01 (Orco) | 10 | 270 | M0 | M1a | M7 | |
| OR02 | 11 | 1128 | M0 | M1a | M7 | |
| OR03 | 5 | 1104 | M0 | M2a | M8 | 345R |
| OR04 | 9 | 795 | M0 | M1a | M7 | |
| OR05 | 9 | 795 | M0 | M1a | M7 | |
| OR06 | 37 | 312 | M0 | M1a | M7 | |
| OR07 | 8 | 1002 | M0 | M1a | M7 | |
| OR08a | 15 | 444 | M0 | M1a | M7 | |
| OR08b | 31 | 60 | M0 | M1a | M7 | |
| OR09 | 12 | 939 | M0 | M1a | M7 | |
| OR10 | 10 | 1011 | M0 | M1a | M7 | |
| OR11 | 8 | 579 | M0 | M1a | M7 | |
| OR12 | 25 | 468 | M0 | M1a | M7 | |
| OR13 | 11 | 885 | M0 | M1a | M7 | |
| OR14 | 17 | 306 | M0 | M1a | M7 | |
| OR15 | 14 | 678 | M0 | M1a | M7 | |
| OR16 | 15 | 378 | M0 | M1a | M7 | |
| OR17 | 25 | 387 | M0 | M1a | M7 | |
| OR18 | 21 | 702 | M0 | M1a | M7 | |
| OR19 | 21 | 1062 | M0 | M1a | M7 | |
| OR20 | 14 | 1086 | M0 | M1a | M7 | |
| OR21 | 15 | 195 | M0 | M1a | M7 | |
| OR22 | 8 | 618 | M0 | M1a | M7 | |
| OR23 | 22 | 300 | M0 | M1a | M7 | |
| Total | 373 | 15504 | | | | |

*Positive selection sites estimated under model M8 by Bayes empirical Bayes (BEB) approach with posterior probabilities (PPs) > 95%. The overlapping sites between M8 and M2a are underlined.

**Table S17**. Summary of selection test results based on the branch model applied to single-copy orthologous groups. Selection types identified include positive selection (ω > 0), purifying selection (ω < 0), and neutral selection (ω = 0) as inferred from PAML v4 analysis (Yang, 2007).

| | OG01 | OG02 | OG03 | OG04 | OG05 | OG06 | OG07 | OG08 | OG09 | OG10 |
|---|---|---|---|---|---|---|---|---|---|---|
| ***N*** | 13 | 13 | 13 | 13 | 13 | 13 | 13 | 13 | 13 | 13 |
| ***S*** | 477 | 3060 | 1371 | 450 | 738 | 2679 | 1380 | 159 | 192 | 930 |
| | 2.24858 | 0.12296 | 0.50758 | 0.28491 | 0.49332 | 0.06533 | 0.91522 | 0.61627 | 2.26590 | 1.83966 |
| ***lnL*** | -2160.159209 | -12064.036465 | -5078.987736 | -3104.222491 | -5169.454320 | -10524.953899 | -9353.560017 | -763.114795 | -556.016689 | -6222.568903 |
| ***lnL*** | -2151.066978 | -12046.973192 | -5060.117689 | -3010.991489 | -5144.933406 | -10221.358852 | -9340.036163 | -754.339339 | -552.390548 | -6205.533917 |
| ***2ΔL*** | 18.184 | 34.127 | 37.74 | 186.46 | 170.02 | 607.19 | 27.048 | 17.551 | 7.2523 | 34.07 |
| ***K*** | 12 | 12 | 12 | 12 | 12 | 12 | 12 | 12 | 12 | 12 |
| ***alpha*** | 0,05 | 0,05 | 0,05 | 0,05 | 0,05 | 0,05 | 0,05 | 0,05 | 0,05 | 0,05 |
| ***p*** | 0.1102 | 0.0006442 | 0.0001693 | 2.2e-16 | 2.2e-16 | 2.2e-16 | 0.007606 | 0.13 | 0.8405 | 0.0006576 |
| ***χ*** | 21.026 | 21.026 | 21.026 | 21.026 | 21.026 | 21.026 | 21.026 | 21.026 | 21.026 | 21.026 |

| | OG11 | OG12 | OG13 | OG14 | OG15 | OG16 | OG17 | OG18 | OG19 | OG20 |
|---|---|---|---|---|---|---|---|---|---|---|
| ***N*** | 13 | 13 | 13 | 13 | 13 | 13 | 13 | 13 | 13 | 13 |
| ***S*** | 1617 | 1512 | 474 | 2178 | 813 | 945 | 1296 | 1005 | 678 | 939 |
| | 1.22681 | 0.09122 | 2.69628 | 0.31425 | 0.03437 | 0.09813 | 0.10343 | 0.06158 | 0.06811 | 0.30141 |
| ***lnL*** | -13934.579561 | -6346.701804 | -2279.718515 | -12827.691404 | -2882.013677 | -3921.725208 | -5865.130797 | -3778.694526 | -2451.561446 | -5439.865077 |
| ***lnL*** | -13926.456658 | -6329.805317 | -2268.388919 | -12797.149749 | -2871.227283 | -3910.795705 | -5588.520143 | -3746.812685 | -2438.768897 | -5407.142548 |
| ***2ΔL*** | 16.246 | 33.793 | 22.659 | 61.083 | 21.573 | 21.859 | 553.22 | 63.764 | 25.585 | 65.445 |
| ***K*** | 12 | 12 | 12 | 12 | 12 | 12 | 12 | 12 | 12 | 12 |
| ***alpha*** | 0,05 | 0,05 | 0,05 | 0,05 | 0,05 | 0,05 | 0,05 | 0,05 | 0,05 | 0,05 |
| ***p*** | 0.1802 | 0.0007271 | 0.03076 | 1.432e-08 | 0.0426 | 0.03913 | 2.2e-16 | 4.611e-09 | 0.01228 | 2.256e-09 |
| ***χ*** | 21.026 | 21.026 | 21.026 | 21.026 | 21.026 | 21.026 | 21.026 | 21.026 | 21.026 | 21.026 |

**Table S17 continuation (OG 21 to 40)**. Summary of selection test results based on the branch model applied to single-copy orthologous groups. Selection types identified include positive selection (ω > 0), purifying selection (ω < 0), and neutral selection (ω = 0) as inferred from PAML v4 analysis (Yang, 2007).

| | **OG21** | **OG22** | **OG23** | **OG24** | **OG25** | **OG26** | **OG27** | **OG28** | **OG29** | **OG30** |
|---|---|---|---|---|---|---|---|---|---|---|
| ***N*** | 13 | 13 | 13 | 13 | 13 | 13 | 13 | 13 | 13 | 13 |
| ***S*** | 1317 | 537 | 435 | 600 | 909 | 336 | 606 | 1575 | 477 | 702 |
| | 3.27177 | 0.06587 | 0.25804 | 0.02001 | 0.29891 | 0.05698 | 0.63361 | 0.38606 | 2.24858 | 0.23299 |
| ***lnL*** | -7167.360574 | -1972.640187 | -2845.469017 | -1942.623794 | -6987.667834 | -1181.017753 | -4152.181626 | -11145.162243 | -3419.975689 | -4661.712646 |
| ***lnL*** | -7127.128098 | -1959.182468 | -2799.426152 | -1927.852835 | -6868.104794 | -1171.684502 | -4118.493890 | -10923.892619 | -3299.384156 | -4457.970644 |
| ***2ΔL*** | 80.465 | 26.915 | 92.086 | 29.542 | 239.13 | 18.667 | 67.375 | 442.54 | 241.18 | 407.48 |
| ***K*** | 12 | 12 | 12 | 12 | 12 | 12 | 12 | 12 | 12 | 12 |
| ***alpha*** | 0,05 | 0,05 | 0,05 | 0,05 | 0,05 | 0,05 | 0,05 | 0,05 | 0,05 | 0,05 |
| ***p*** | 3.365e-12 | 0.007947 | 1.946e-14 | 0.003271 | 2.2e-16 | 0.0969 | 9.888e-10 | 2.2e-16 | 2.2e-16 | 2.2e-16 |
| ***χ*** | 21.026 | 21.026 | 21.026 | 21.026 | 21.026 | 21.026 | 21.026 | 21.026 | 21.026 | 21.026 |

| | **OG31** | **OG32** | **OG33** | **OG34** | **OG35** | **OG36** | **OG37** | **OG38** | **OG39** | **OG40** |
|---|---|---|---|---|---|---|---|---|---|---|
| ***N*** | 13 | 13 | 13 | 13 | 13 | 13 | 13 | 13 | 13 | 13 |
| ***S*** | 1527 | 435 | 1854 | 1800 | 1029 | 879 | 1707 | 480 | 840 | 1425 |
| | 0.46942 | 1.12776 | 0.78526 | 0.22210 | 2.81737 | 0.04742 | 0.27643 | 0.93398 | 0.24967 | 0.13977 |
| ***lnL*** | -8759.946303 | -2848.854168 | -14544.838549 | -8886.695081 | -3913.272980 | -2874.368149 | -13163.628869 | -3897.522790 | -5862.293072 | -6962.314369 |
| ***lnL*** | -8751.416317 | -2840.262743 | -14508.528505 | -8867.252088 | -3902.348755 | -2858.457621 | -12674.445587 | -3885.905342 | -5699.290353 | -6772.689588 |
| ***2ΔL*** | 17.06 | 17.183 | 72.62 | 38.886 | 21.848 | 31.821 | 978.37 | 23.235 | 326.01 | 379.25 |
| ***K*** | 12 | 12 | 12 | 12 | 12 | 12 | 12 | 12 | 12 | 12 |
| ***alpha*** | 0,05 | 0,05 | 0,05 | 0,05 | 0,05 | 0,05 | 0,05 | 0,05 | 0,05 | 0,05 |
| ***p*** | 0.1474 | 0.1428 | 1.033e-10 | 0.0001099 | 0.03925 | 0.001475 | 2.2e-16 | 0.0258 | 2.2e-16 | 2.2e-16 |
| ***χ*** | 21.026 | 21.026 | 21.026 | 21.026 | 21.026 | 21.026 | 21.026 | 21.026 | 21.026 | 21.026 |

**Table S17 continuation (OG 41 to 50)**. Summary of selection test results based on the branch model applied to single-copy orthologous groups. Selection types identified include positive selection (ω > 0), purifying selection (ω < 0), and neutral selection (ω = 0) as inferred from PAML v4 analysis (Yang, 2007).

| | **OG41** | **OG42** | **OG43** | **OG44** | **OG45** | **OG46** | **OG47** | **OG48** | **OG49** | **OG50** |
|---|---|---|---|---|---|---|---|---|---|---|
| ***N*** | 13 | 13 | 13 | 13 | 13 | 13 | 13 | 13 | 13 | 13 |
| ***S*** | 1425 | 609 | 795 | 912 | 1434 | 948 | 696 | 297 | 1806 | 495 |
| | 3.00446 | 1.16905 | 4.06583 | 5.07984 | 0.21805 | 0.02876 | 0.23508 | 1.81814 | 0.03415 | 0.11660 |
| ***lnL*** | -1708.725361 | -2766.130509 | -3105.508058 | -3766.866329 | -9067.084798 | -3342.235869 | -3052.410174 | -1334.131281 | -6072.369179 | -2532.568823 |
| ***lnL*** | -1695.593096 | -2755.461167 | -3092.621569 | -3756.804592 | -8838.336867 | -3317.490890 | -3035.356911 | -1324.903251 | -6057.761169 | -2478.000576 |
| ***2ΔL*** | 26.265 | 21.339 | 25.773 | 20.123 | 457.5 | 49.49 | 34.107 | 18.456 | 29.216 | 109.14 |
| ***K*** | 12 | 12 | 12 | 12 | 12 | 12 | 12 | 12 | 12 | 12 |
| ***alpha*** | 0,05 | 0,05 | 0,05 | 0,05 | 0,05 | 0,05 | 0,05 | 0,05 | 0,05 | 0,05 |
| ***p*** | 0.009846 | 0.04564 | 0.01156 | 0.06479 | 2.2e-16 | 1.717e-06 | 0.0006489 | 0.1025 | 0.003659 | 2.2e-16 |
| ***χ*** | 21.026 | 21.026 | 21.026 | 21.026 | 21.026 | 21.026 | 21.026 | 21.026 | 21.026 | 21.026 |

**Table S18**. Single-copy orthologous groups (OGs) classified under positive selection (ω > 0), purifying selection (ω < 0), and neutral selection (ω = 0), based on branch model analysis using PAML v4 (Yang, 2007).

| Genes | Number of genes | Sites | Free Omega | ω>0 | ω<0 | ω=0 | Saturated |
|---|---|---|---|---|---|---|---|
| **OG01** | 13 | 477 | No | **4** | **1** | **0** | 8 |
| **OG02** | 13 | 3060 | Yes | **0** | **13** | **0** | 0 |
| **OG03** | 13 | 1371 | Yes | **1** | **11** | **0** | 1 |
| **OG04** | 13 | 450 | Yes | **1** | **11** | **0** | 1 |
| **OG05** | 13 | 738 | Yes | **1** | **11** | **0** | 1 |
| **OG06** | 13 | 2679 | Yes | **0** | **13** | **0** | 0 |
| **OG07** | 13 | 1380 | Yes | **2** | **9** | **0** | 2 |
| **OG08** | 13 | 159 | No | **1** | **8** | **0** | 4 |
| **OG09** | 13 | 192 | No | **4** | **3** | **0** | 6 |
| **OG10** | 13 | 930 | Yes | **4** | **2** | **0** | 7 |
| **OG11** | 13 | 1617 | No | **10** | **2** | **0** | 1 |
| **OG12** | 13 | 1512 | Yes | **0** | **13** | **0** | 0 |
| **OG13** | 13 | 474 | Yes | **4** | **1** | **0** | 8 |
| **OG14** | 13 | 2178 | Yes | **1** | **12** | **0** | 0 |
| **OG15** | 13 | 813 | Yes | **0** | **13** | **0** | 0 |
| **OG16** | 13 | 945 | Yes | **0** | **13** | **0** | 0 |
| **OG17** | 13 | 1296 | Yes | **0** | **13** | **0** | 0 |
| **OG18** | 13 | 1005 | Yes | **1** | **12** | **0** | 0 |
| **OG19** | 13 | 678 | Yes | **0** | **13** | **0** | 0 |
| **OG20** | 13 | 939 | Yes | **0** | **13** | **0** | 0 |
| **OG21** | 13 | 1317 | Yes | **0** | **3** | **0** | 10 |
| **OG22** | 13 | 537 | Yes | **1** | **12** | **0** | 0 |
| **OG23** | 13 | 435 | Yes | **0** | **11** | **0** | 2 |
| **OG24** | 13 | 600 | Yes | **0** | **12** | **0** | 1 |
| **OG25** | 13 | 909 | Yes | **1** | **12** | **0** | 0 |
| **OG26** | 13 | 336 | Yes | **0** | **13** | **0** | 0 |
| **OG27** | 13 | 606 | Yes | **2** | **11** | **0** | 0 |
| **OG28** | 13 | 1575 | Ha | **0** | **11** | **0** | 2 |
| **OG29** | 13 | 477 | Yes | **2** | **9** | **0** | 2 |
| **OG30** | 13 | 702 | Yes | **1** | **12** | **0** | 0 |

**Table S18 continuation**. Single-copy orthologous groups (OGs) classified under positive selection (ω > 0), purifying selection (ω < 0), and neutral selection (ω = 0), based on branch model analysis using PAML v4 (Yang, 2007).

| Genes | Number of genes | Sites | Free Omega | ω>0 | ω<0 | ω=0 | Saturated |
|---|---|---|---|---|---|---|---|
| **OG31** | 13 | 1527 | No | **1** | **12** | **0** | 0 |
| **OG32** | 13 | 435 | No | **6** | **2** | **0** | 5 |
| **OG33** | 13 | 1854 | Yes | **2** | **11** | **0** | 0 |
| **OG34** | 13 | 1800 | Yes | **0** | **13** | **0** | 0 |
| **OG35** | 13 | 1029 | Yes | **2** | **1** | **0** | 10 |
| **OG36** | 13 | 879 | Yes | **0** | **13** | **0** | 0 |
| **OG37** | 13 | 1707 | Yes | **1** | **12** | **0** | 0 |
| **OG38** | 13 | 480 | Yes | **3** | **9** | **0** | 1 |
| **OG39** | 13 | 840 | Yes | 0 | 13 | 0 | 0 |
| **OG40** | 13 | 1425 | Yes | 1 | 12 | 0 | 0 |
| **OG41** | 13 | 1425 | Yes | 3 | 1 | 0 | 9 |
| **OG42** | 13 | 609 | Yes | 5 | 4 | 0 | 4 |
| **OG43** | 13 | 795 | Yes | 1 | 0 | 0 | 12 |
| **OG44** | 13 | 912 | No | 1 | 0 | 0 | 12 |
| **OG45** | 13 | 1434 | Yes | 1 | 12 | 0 | 0 |
| **OG46** | 13 | 948 | Yes | 0 | 13 | 0 | 0 |
| **OG47** | 13 | 696 | Yes | 1 | 10 | 0 | 2 |
| **OG48** | 13 | 297 | Yes | 1 | 4 | 0 | 8 |
| **OG49** | 13 | 1806 | Yes | 0 | 13 | 0 | 0 |
| **OG50** | 13 | 495 | Yes | 0 | 13 | 0 | 0 |
| **Total** | 650 | 51780 | | 70 | 461 | 0 | 119 |

**Table S19**. Summary statistics for branch-specific evolutionary models of single-copy orthologous subfamilies, analyzed using PAML v4 (Yang, 2007).

| Model | Statistics | OG01 | OG02 | OG03 | OG04 | OG05 | OG06 | OG07 | OG08 | OG09 | OG10 |
|---|---|---|---|---|---|---|---|---|---|---|---|
| | ***N*** | 13 | 13 | 13 | 13 | 13 | 13 | 13 | 13 | 13 | 13 |
| | ***S*** | 477 | 3060 | 1371 | 450 | 738 | 2679 | 1380 | 159 | 192 | 930 |
| **M0** | ***ω*** | 2.03903 | 0.12625 | 0.49153 | 0.31310 | 0.58051 | 0.06688 | 0.96506 | 0.63003 | 1.98810 | 2.06902 |
| | ***lnL*** | -2162.948435 | -12069.630996 | -5080.833417 | -3121.452981 | -5223.453997 | -10548.369935 | -9359.540371 | -763.252274 | -557.219077 | -6236.572796 |
| **M1a** | ***p0*** | 0.25982 | 0.93998 | 0.66882 | 0.84218 | 0.77591 | 0.96046 | 0.69863 | 0.69878 | 0.00001 | 0.68480 |
| | ***p1*** | 0.74018 | 0.06002 | 0.33118 | 0.15782 | 0.22409 | 0.03954 | 0.30137 | 0.30122 | 0.99999 | 0.31520 |
| | ***lnL*** | -2169.235915 | -11808.787819 | -4986.978991 | -3034.793061 | -4977.844425 | -10364.330361 | -9050.292984 | -748.377623 | -558.740901 | -6083.510086 |
| **M2a** | ***p0*** | 0.53529 | 0.93998 | 0.70344 | 0.84029 | 0.76595 | 0.96046 | 0.65404 | 0.77167 | 0.48256 | 0.62337 |
| | ***p1*** | 0.33891 | 0.06002 | 0.25842 | 0.15850 | 0.19908 | 0.00841 | 0.27025 | 0.18466 | 0.00000 | 0.21548 |
| | ***p2*** | 0.12580 | 0.00000 | 0.03814 | 0.00122 | 0.03497 | 0.03113 | 0.07571 | 0.04367 | 0.51744 | 0.16115 |
| | ***ω2*** | 7.63607 | 27.59557 | 3.00940 | 4.36479 | 2.65723 | 1.00000 | 3.64310 | 3.88546 | 4.01022 | 5.72098 |
| | ***lnL*** | -2120.742582 | -11808.787825 | -4977.452908 | -3033.125159 | -4963.865531 | -10364.330360 | -8947.296962 | -743.102722 | -553.225759 | -5937.927170 |
| **M3** | ***p0*** | 0.83666 | 0.84481 | 0.77514 | 0.70166 | 0.76994 | 0.87104 | 0.70207 | 0.87603 | 0.48256 | 0.70541 |
| | ***p1*** | 0.16304 | 0.14619 | 0.21972 | 0.29002 | 0.20264 | 0.01563 | 0.24234 | 0.12193 | 0.11207 | 0.19961 |
| | ***p2*** | 0.00030 | 0.00900 | 0.00515 | 0.00831 | 0.02742 | 0.11333 | 0.05559 | 0.00204 | 0.40537 | 0.09499 |
| | ***ω0*** | 0.96768 | 0.01450 | 0.06293 | 0.00000 | 0.00000 | 0.00000 | 0.04499 | 0.19065 | 0.00000 | 0.01740 |
| | ***ω1*** | 6.52139 | 0.41428 | 1.45284 | 0.54568 | 1.06355 | 0.00000 | 1.28657 | 2.27276 | 4.01022 | 2.43871 |
| | ***ω2*** | 82.49708 | 1.96107 | 5.42739 | 2.60606 | 2.84132 | 0.42166 | 4.04399 | 11.24185 | 4.01022 | 6.94049 |
| | ***lnL*** | -2112.487363 | -11800.164060 | -4977.163013 | -3028.144815 | -4963.837256 | -10328.578162 | -8946.887586 | -742.909191 | -553.225759 | -5937.036209 |

**Table S19 continuation**. Summary statistics for branch-specific evolutionary models of single-copy orthologous subfamilies, analyzed using PAML v4 (Yang, 2007).

| Mode l | Statistics | OG11 | OG12 | OG13 | OG14 | OG15 | OG16 | OG17 | OG18 | OG19 | OG20 |
|---|---|---|---|---|---|---|---|---|---|---|---|
| | ***N*** | 13 | 13 | 13 | 13 | 13 | 13 | 13 | 13 | 13 | 13 |
| | ***S*** | 1617 | 1512 | 474 | 2178 | 813 | 945 | 1296 | 1005 | 678 | 939 |
| **M0** | ***ω*** | 1.27020 | 0.09304 | 2.51172 | 0.31578 | 0.03617 | 0.09932 | 0.10720 | 0.06236 | 0.10720 | 0.06236 |
| | ***lnL*** | -14004.587976 | -6347.668937 | -2282.294164 | -12827.962628 | -2885.791887 | -3922.364464 | -5887.196356 | -3779.892485 | -5887.196356 | -3779.892485 |
| **M1a** | ***p0*** | 0.63453 | 0.95844 | 0.25594 | 0.83268 | 0.98920 | 0.95672 | 0.92350 | 0.96656 | 0.92350 | 0.96656 |
| | ***p1*** | 0.36547 | 0.04156 | 0.74406 | 0.16732 | 0.01080 | 0.04328 | 0.07650 | 0.03344 | 0.07650 | 0.03344 |
| | ***lnL*** | -13663.361889 | -6211.419555 | -2297.410342 | -12519.847267 | -2836.443831 | -3834.113964 | -5765.926799 | -3693.393850 | -5765.926799 | -3693.393850 |
| **M2a** | ***p0*** | 0.59730 | 0.95844 | 0.49500 | 0.83268 | 0.99005 | 0.95672 | 0.92350 | 0.96656 | 0.92350 | 0.96656 |
| | ***p1*** | 0.29631 | 0.04156 | 0.29370 | 0.09452 | 0.00926 | 0.04328 | 0.04754 | 0.01417 | 0.04754 | 0.01417 |
| | ***p2*** | 0.10640 | 0.00000 | 0.21130 | 0.07280 | 0.00069 | 0.00000 | 0.02896 | 0.01926 | 0.02896 | 0.01926 |
| | ***ω2*** | 3.78312 | 29.83620 | 30.60936 | 1.00000 | 2.17707 | 35.25915 | 1.00000 | 1.00000 | 1.00000 | 1.00000 |
| | ***lnL*** | -13467.529096 | -6211.419555 | -2245.293250 | -12519.847267 | -2836.296475 | -3834.113964 | -5765.926798 | -3693.393850 | -5765.926798 | -3693.393850 |
| **M3** | ***p0*** | 0.65100 | 0.21929 | 0.68509 | 0.57719 | 0.67016 | 0.75483 | 0.87749 | 0.90476 | 0.87749 | 0.90476 |
| | ***p1*** | 0.30146 | 0.72816 | 0.31003 | 0.35780 | 0.32330 | 0.22457 | 0.07794 | 0.09465 | 0.07794 | 0.09465 |
| | ***p2*** | 0.04754 | 0.05255 | 0.00488 | 0.06500 | 0.00653 | 0.02060 | 0.04457 | 0.00059 | 0.04457 | 0.00059 |
| | ***ω0*** | 0.00855 | 0.02486 | 0.92539 | 0.00000 | 0.00000 | 0.00000 | 0.00000 | 0.00000 | 0.00000 | 0.00000 |
| | ***ω1*** | 1.57218 | 0.02486 | 5.61221 | 0.33608 | 0.05044 | 0.19885 | 0.60403 | 0.39796 | 0.60403 | 0.39796 |
| | ***ω2*** | 5.02400 | 0.80591 | 30.60936 | 1.43687 | 1.32205 | 1.11094 | 0.60404 | 4.69666 | 0.60404 | 4.69666 |
| | ***lnL*** | -13464.492160 | -6209.966418 | -2241.487657 | -12510.085125 | -2836.115816 | -3825.078218 | -5754.333055 | -3682.333934 | -5754.333055 | -3682.333934 |

**Table S19 continuation**. Summary statistics for branch-specific evolutionary models of single-copy orthologous subfamilies, analyzed using PAML v4 (Yang, 2007).

| Model | Statistics | OG21 | OG22 | OG23 | OG24 | OG25 | OG26 | OG27 | OG28 | OG29 | OG30 |
|---|---|---|---|---|---|---|---|---|---|---|---|
| | ***N*** | 13 | 13 | 13 | 13 | 13 | 13 | 13 | 13 | 13 | 13 |
| | ***S*** | 1317 | 537 | 435 | 600 | 909 | 336 | 606 | 1575 | 477 | 702 |
| **M0** | ***ω*** | 3.15645 | 0.06869 | 0.32988 | 0.02128 | 0.37197 | 0.05701 | 0.75611 | 0.48197 | 0.17739 | 0.24196 |
| | ***lnL*** | -7173.277608 | -1973.476058 | -2893.070677 | -1945.235038 | -7079.463353 | -1181.331501 | -4165.726469 | -11255.347165 | -3447.167177 | -4718.510529 |
| **M1a** | ***p0*** | 0.34621 | 0.97365 | 0.87254 | 0.98696 | 0.84245 | 0.98405 | 0.76337 | 0.80323 | 0.89282 | 0.94565 |
| | ***p1*** | 0.65379 | 0.02635 | 0.12746 | 0.01304 | 0.15755 | 0.01595 | 0.23663 | 0.19677 | 0.10718 | 0.05435 |
| | ***lnL*** | -7228.052639 | -1943.028431 | -2749.813253 | -1919.354366 | -6804.813715 | -1169.588702 | -4009.024521 | -10796.382824 | -3363.304373 | -4647.758019 |
| **M2a** | ***p0*** | 0.25129 | 0.97365 | 0.87022 | 0.98696 | 0.83854 | 0.98405 | 0.74170 | 0.79261 | 0.89281 | 0.94565 |
| | ***p1*** | 0.53641 | 0.02635 | 0.12640 | 0.00219 | 0.15942 | 0.00105 | 0.21948 | 0.19533 | 0.10719 | 0.01669 |
| | ***p2*** | 0.21230 | 0.00000 | 0.00338 | 0.01084 | 0.00204 | 0.01490 | 0.03882 | 0.01206 | 0.00000 | 0.03766 |
| | ***ω2*** | 8.43115 | 43.72036 | 3.61611 | 1.00000 | 4.04644 | 1.00000 | 3.84791 | 3.81021 | 13.80350 | 1.00000 |
| | ***lnL*** | -7015.381481 | -1943.028431 | -2747.641931 | -1919.354367 | -6801.142675 | -1169.588702 | -3981.442258 | -10767.549294 | -3363.304385 | -4647.758019 |
| **M3** | ***p0*** | 0.52851 | 0.52210 | 0.8101 | 0.03091 | 0.65664 | 0.11535 | 0.79904 | 0.76887 | 0.74728 | 0.56162 |
| | ***p1*** | 0.37439 | 0.45088 | 0.18157 | 0.95034 | 0.32681 | 0.86845 | 0.18712 | 0.21466 | 0.25199 | 0.32317 |
| | ***p2*** | 0.09710 | 0.02702 | 0.00831 | 0.01876 | 0.01655 | 0.01621 | 0.01384 | 0.01647 | 0.00073 | 0.11521 |
| | ***ω0*** | 0.48277 | 0.00000 | 0.02082 | 0.00000 | 0.00000 | 0.02986 | 0.04974 | 0.01540 | 0.00000 | 0.00000 |
| | ***ω1*** | 3.44893 | 0.05436 | 0.66638 | 0.00000 | 0.45069 | 0.02986 | 1.52866 | 0.88168 | 0.44942 | 0.40801 |
| | ***ω2*** | 11.61563 | 0.90949 | 2.83159 | 0.72032 | 2.10823 | 0.98764 | 5.53708 | 3.46481 | 3.96368 | 0.40801 |
| | ***lnL*** | -7012.525052 | -1942.808084 | -2746.114847 | -1918.986824 | -6782.359176 | -1169.588354 | -3980.362094 | -10767.412053 | -3347.711898 | -4625.850279 |

**Table S19 continuation**. Summary statistics for branch-specific evolutionary models of single-copy orthologous subfamilies, analyzed using PAML v4 (Yang, 2007).

| Model | Statistics | OG31 | OG32 | OG33 | OG34 | OG35 | OG36 | OG37 | OG38 | OG39 | OG40 |
|---|---|---|---|---|---|---|---|---|---|---|---|
| | ***N*** | 13 | 13 | 13 | 13 | 13 | 13 | 13 | 13 | 13 | 13 |
| | ***S*** | 1527 | 435 | 1854 | 1800 | 1029 | 879 | 1707 | 480 | 840 | 1425 |
| **M0** | ***ω*** | 0.45877 | 1.34443 | 0.92485 | 0.23130 | 2.36009 | 0.04643 | 0.32492 | 1.13044 | 0.28521 | 0.15448 |
| | ***lnL*** | -8768.449410 | -2868.612793 | -14621.285148 | -8894.045372 | -3929.296093 | -2874.910280 | -13312.710408 | -3928.586702 | -5930.825560 | -7009.639226 |
| **M1a** | ***p0*** | 0.76233 | 0.73731 | 0.68654 | 0.89133 | 0.17106 | 0.98330 | 0.92134 | 0.72567 | 0.91023 | 0.92920 |
| | ***p1*** | 0.23767 | 0.26269 | 0.31346 | 0.10867 | 0.82894 | 0.01670 | 0.07866 | 0.27433 | 0.08977 | 0.07080 |
| | ***lnL*** | -8564.647612 | -2752.368582 | -14140.983283 | -8644.277513 | -3942.697472 | -2853.656918 | -13093.487688 | -3756.388973 | -5805.481261 | -6890.667418 |
| **M2a** | ***p0*** | 0.75935 | 0.65871 | 0.63893 | 0.88985 | 0.21863 | 0.98330 | 0.92134 | 0.66751 | 0.91022 | 0.92920 |
| | ***p1*** | 0.23161 | 0.28368 | 0.30690 | 0.10982 | 0.61608 | 0.00645 | 0.05631 | 0.26635 | 0.04861 | 0.07080 |
| | ***p2*** | 0.00904 | 0.05761 | 0.05417 | 0.00033 | 0.16529 | 0.01025 | 0.02235 | 0.06614 | 0.04116 | 0.00000 |
| | ***ω2*** | 3.98219 | 5.98054 | 3.35478 | 9.17946 | 8.67694 | 1.00000 | 1.00000 | 3.93325 | 1.00000 | 16.97923 |
| | ***lnL*** | -8545.108356 | -2690.644641 | -14061.772045 | -8635.257293 | -3856.077852 | -2853.656918 | -13093.487688 | -3712.184796 | -5805.481261 | -6890.667420 |
| **M3** | ***p0*** | 0.77399 | 0.74277 | 0.59986 | 0.82584 | 0.58799 | 0.79781 | 0.49340 | 0.75091 | 0.53446 | 0.70661 |
| | ***p1*** | 0.21901 | 0.24127 | 0.32875 | 0.16955 | 0.37949 | 0.19925 | 0.50364 | 0.23725 | 0.46308 | 0.28955 |
| | ***p2*** | 0.00700 | 0.01596 | 0.07139 | 0.00462 | 0.03252 | 0.00294 | 0.00296 | 0.01184 | 0.00246 | 0.00384 |
| | ***ω0*** | 0.08158 | 0.00072 | 0.00097 | 0.03691 | 0.83729 | 0.00000 | 0.00000 | 0.03166 | 0.00000 | 0.00000 |
| | ***ω1*** | 1.06919 | 2.21512 | 0.83809 | 0.63693 | 4.26197 | 0.16510 | 0.41658 | 1.88447 | 0.37786 | 0.34956 |
| | ***ω2*** | 4.33771 | 10.76177 | 3.05736 | 3.06968 | 20.76164 | 2.04210 | 2.90455 | 7.17490 | 4.53184 | 1.99404 |
| | ***lnL*** | -8544.994586 | -2687.374379 | -14061.496172 | -8630.576510 | -3850.677314 | -2853.074629 | -13065.035639 | -3710.943836 | -5773.580811 | -6872.342203 |

**Table S19 continuation**. Summary statistics for branch-specific evolutionary models of single-copy orthologous subfamilies, analyzed using PAML v4 (Yang, 2007).

| **Model** | **Statistics** | **OG41** | **OG42** | **OG43** | **OG44** | **OG45** | **OG46** | **OG47** | **OG48** | **OG49** | **OG50** |
|---|---|---|---|---|---|---|---|---|---|---|---|
| | ***N*** | 13 | 13 | 13 | 13 | 13 | 13 | 13 | 13 | 13 | 13 |
| | ***S*** | 1425 | 609 | 795 | 912 | 1434 | 948 | 696 | 297 | 1806 | 495 |
| **M0** | ***ω*** | 3.01909 | 1.15538 | 3.53209 | 4.26656 | 0.25442 | 0.03075 | 0.24336 | 1.64748 | 0.03449 | 0.14152 |
| | ***lnL*** | -1708.733339 | -2773.327806 | -3118.301126 | -3785.486916 | -9144.155714 | -3347.179895 | -3053.244689 | -1345.203631 | -6073.114803 | -2544.769522 |
| **M1a** | ***p0*** | 0.33890 | 0.35150 | 0.06555 | 0.00001 | 0.89824 | 0.98803 | 0.88764 | 0.29450 | 0.98742 | 0.92757 |
| | ***p1*** | 0.66110 | 0.64850 | 0.93445 | 0.99999 | 0.10176 | 0.01197 | 0.11236 | 0.70550 | 0.01258 | 0.07243 |
| | ***lnL*** | -1719.991061 | -2753.946889 | -3152.415326 | -3843.277682 | -8919.315424 | -3303.994599 | -3004.787145 | -1335.653420 | -6008.522814 | -2505.373631 |
| **M2a** | ***p0*** | 0.37669 | 0.39400 | 0.71781 | 0.03567 | 0.89824 | 0.98803 | 0.89076 | 0.75688 | 0.98742 | 0.92757 |
| | ***p1*** | 0.44150 | 0.33095 | 0.02892 | 0.67106 | 0.04521 | 0.01197 | 0.10743 | 0.00000 | 0.00385 | 0.00796 |
| | ***p2*** | 0.18181 | 0.27505 | 0.25328 | 0.29327 | 0.05655 | 0.00000 | 0.00181 | 0.24312 | 0.00873 | 0.06447 |
| | ***ω2*** | 7.16173 | 2.37170 | 8.59261 | 10.38207 | 1.00000 | 1.00000 | 3.06773 | 4.20740 | 1.00000 | 1.00000 |
| | ***lnL*** | -1676.231683 | -2741.944017 | -3069.716063 | -3713.752060 | -8919.315424 | -3303.994599 | -3004.514350 | -1315.858938 | -6008.522814 | -2505.373631 |
| **M3** | ***p0*** | 0.32369 | 0.42989 | 0.72723 | 0.16197 | 0.63657 | 0.98235 | 0.83758 | 0.70683 | 0.00002 | 0.36931 |
| | ***p1*** | 0.61593 | 0.55964 | 0.21808 | 0.70016 | 0.35649 | 0.00000 | 0.14373 | 0.05005 | 0.98385 | 0.41799 |
| | ***p2*** | 0.06038 | 0.01046 | 0.05469 | 0.13786 | 0.00694 | 0.01765 | 0.01870 | 0.24312 | 0.01613 | 0.21270 |
| | ***ω0*** | 0.00000 | 0.00000 | 1.80838 | 0.00000 | 0.00000 | 0.00735 | 0.06465 | 0.59565 | 0.01229 | 0.00615 |
| | ***ω1*** | 2.80261 | 1.64296 | 6.37883 | 3.08324 | 0.37640 | 0.04821 | 0.65885 | 0.59565 | 0.01324 | 0.00621 |
| | ***ω2*** | 12.47007 | 6.90660 | 19.93826 | 17.55211 | 2.23659 | 0.71720 | 1.82138 | 4.20740 | 0.82041 | 0.39239 |
| | ***lnL*** | -1673.816166 | -2741.298992 | -3064.991199 | -3709.550828 | -8887.890899 | -3303.396341 | -3004.355660 | -1315.858938 | -6008.146911 | -2494.922373 |

**Table S19 continuation**. Summary statistics for branch-specific evolutionary models of single-copy orthologous subfamilies, analyzed using PAML v4 (Yang, 2007).
2007).

| Model | Statistics | OG01 | OG02 | OG03 | OG04 | OG05 | OG06 | OG07 | OG08 | OG09 | OG10 |
|---|---|---|---|---|---|---|---|---|---|---|---|
| **M7** | ***p*** | 3.99238 | 0.07707 | 0.00581 | 0.09053 | 0.00500 | 0.02946 | 0.01058 | 0.00563 | 7.28889 | 0.00500 |
| | ***q*** | 0.00500 | 0.72204 | 0.01124 | 0.42311 | 0.01944 | 0.53253 | 0.02130 | 0.00707 | 0.00500 | 0.01165 |
| | ***lnL*** | -2174.312191 | -11814.932584 | -4987.502003 | -3032.430812 | -4978.958372 | -10329.253345 | -9050.732544 | -748.936935 | -558.740903 | -6083.778050 |
| **M8** | ***p0*** | 0.87420 | 0.99124 | 0.95553 | 0.99660 | 0.96159 | 0.99929 | 0.92585 | 0.93849 | 0.48256 | 0.83322 |
| | ***p1*** | 0.12580 | 0.00876 | 0.04447 | 0.00340 | 0.03841 | 0.00071 | 0.07415 | 0.06151 | 0.51744 | 0.16678 |
| | ***p*** | 2.75653 | 0.12909 | 0.05686 | 0.12628 | 0.00502 | 0.02987 | 0.01847 | 0.58485 | 0.00500 | 0.01733 |
| | ***q*** | 0.00500 | 1.50574 | 0.14691 | 0.60838 | 0.02101 | 0.55288 | 0.03754 | 1.55468 | 2.06968 | 0.05382 |
| | ***ω*** | 7.63607 | 1.96894 | 2.89305 | 3.26448 | 2.57060 | 1.61632 | 3.69431 | 3.43188 | 4.01022 | 5.63760 |
| | ***lnL*** | -2120.742584 | -11800.268657 | -4977.528737 | -3029.343683 | -4963.886655 | -10328.535146 | -8947.455539 | -743.192940 | -553.225759 | -5938.151822 |

**Table S19 continuation**. Summary statistics for branch-specific evolutionary models of single-copy orthologous subfamilies, analyzed using PAML v4 (Yang, 2007).

| Model | Statistics | OG11 | OG12 | OG13 | OG14 | OG15 | OG16 | OG17 | OG18 | OG19 | OG20 |
|---|---|---|---|---|---|---|---|---|---|---|---|
| **M7** | ***p*** | 0.00500 | 0.07424 | 64.77837 | 0.12198 | 0.04111 | 0.07992 | 0.01493 | 0.01399 | 0.01493 | 0.01399 |
| | ***q*** | 0.00749 | 0.91728 | 0.00500 | 0.46995 | 1.04170 | 0.97517 | 0.23436 | 0.27037 | 0.23436 | 0.27037 |
| | ***lnL*** | -13665.327862 | -6218.707134 | -2303.037618 | -12521.830412 | -2846.434892 | -3829.332524 | -5755.809986 | -3688.029348 | -5755.809986 | -3688.029348 |
| **M8** | ***p0*** | 0.88231 | 0.96369 | 0.78870 | 0.97109 | 0.99347 | 0.98088 | 0.99999 | 0.99938 | 0.99999 | 0.99938 |
| | ***p1*** | 0.11769 | 0.03631 | 0.21130 | 0.02891 | 0.00653 | 0.01912 | 0.00001 | 0.00062 | 0.00001 | 0.00062 |
| | ***p*** | 0.00500 | 0.46232 | 98.99668 | 0.20687 | 0.31352 | 0.13805 | 0.01230 | 0.00550 | 0.01230 | 0.00550 |
| | ***q*** | 0.01164 | 13.24185 | 0.00500 | 1.01252 | 17.22106 | 2.50834 | 0.19903 | 0.10876 | 0.19903 | 0.10876 |
| | ***ω*** | 3.61834 | 1.00000 | 6.86935 | 1.76407 | 1.32106 | 1.11066 | 1.69552 | 4.60556 | 1.69552 | 4.60556 |
| | ***lnL*** | -13467.984638 | -6210.890798 | -2245.293250 | -12510.567346 | -2836.132958 | -3825.849050 | -5755.813062 | -3682.383972 | -5755.813062 | -3682.383972 |

**Table S19 continuation**. Summary statistics for branch-specific evolutionary models of single-copy orthologous subfamilies, analyzed using PAML v4 (Yang, 2007).

| Model | Statistics | OG21 | OG22 | OG23 | OG24 | OG25 | OG26 | OG27 | OG28 | OG29 | OG30 |
|---|---|---|---|---|---|---|---|---|---|---|---|
| **M7** | ***p*** | 2.14132 | 0.06406 | 0.06593 | 0.01157 | 0.11791 | 0.09797 | 0.00500 | 0.01324 | 0.12438 | 0.39905 |
| | ***q*** | 0.00500 | 1.03157 | 0.37167 | 0.29128 | 0.55212 | 1.71366 | 0.01043 | 0.04401 | 0.91871 | 1.75626 |
| | ***lnL*** | -7260.652141 | -1945.359118 | -2750.956795 | -1925.351999 | -6793.559395 | -1171.791179 | -4011.459227 | -10797.123547 | -3351.699957 | -4631.216856 |
| **M8** | ***p0*** | 0.78768 | 0.97751 | 0.99254 | 0.98709 | 0.99358 | 0.98435 | 0.95524 | 0.98419 | 0.99947 | 0.99980 |
| | ***p1*** | 0.21232 | 0.02249 | 0.00746 | 0.01291 | 0.00642 | 0.01565 | 0.04476 | 0.01581 | 0.00053 | 0.00020 |
| | ***p*** | 3.80620 | 0.37626 | 0.11086 | 0.01337 | 0.17085 | 3.12902 | 0.04091 | 0.05149 | 0.13684 | 0.42276 |
| | ***q*** | 0.00500 | 12.46098 | 0.66878 | 1.53760 | 0.83982 | 99.00000 | 0.13223 | 0.19849 | 1.02612 | 1.87258 |
| | ***ω*** | 8.43172 | 1.00000 | 2.88271 | 1.00000 | 2.75890 | 1.00000 | 3.66200 | 3.49762 | 4.13548 | 5.57258 |
| | ***lnL*** | -7015.381484 | -1942.826664 | -2746.445973 | -1919.346836 | -6785.007090 | -1169.662614 | -3981.889340 | -10767.435440 | -3350.794888 | -4630.941590 |

**Table S19 continuation**. Summary statistics for branch-specific evolutionary models of single-copy orthologous subfamilies, analyzed using PAML v4 (Yang, 2007).

| Model | Statistics | OG31 | OG32 | OG33 | OG34 | OG35 | OG36 | OG37 | OG38 | OG39 | OG40 |
|---|---|---|---|---|---|---|---|---|---|---|---|
| **M7** | ***p*** | 0.09682 | 0.00500 | 0.01361 | 0.12853 | 2.93311 | 0.05823 | 0.42458 | 0.00500 | 0.27664 | 0.15619 |
| | ***q*** | 0.23303 | 0.01169 | 0.02728 | 0.71945 | 0.00500 | 1.15745 | 1.52297 | 0.01164 | 1.18964 | 1.22612 |
| | ***lnL*** | -8574.059132 | -2753.230060 | -14141.789697 | -8646.652568 | -3947.301727 | -2855.538348 | -13079.471158 | -3756.977191 | -5794.915321 | -6875.501372 |
| **M8** | ***p0*** | 0.97856 | 0.94224 | 0.94255 | 0.99376 | 0.83471 | 0.99740 | 0.99922 | 0.93748 | 0.99805 | 0.99865 |
| | ***p1*** | 0.02144 | 0.05776 | 0.05745 | 0.00624 | 0.16529 | 0.00260 | 0.00078 | 0.06252 | 0.00195 | 0.00135 |
| | ***p*** | 0.19284 | 0.00500 | 0.02227 | 0.18461 | 5.34057 | 0.11980 | 0.51350 | 0.00500 | 0.43635 | 0.17342 |
| | ***q*** | 0.53024 | 0.01181 | 0.04396 | 1.14440 | 0.00500 | 2.99264 | 1.86388 | 0.01182 | 1.94329 | 1.40620 |
| | ***ω*** | 2.99660 | 5.96985 | 3.27229 | 2.75081 | 8.67694 | 2.14111 | 4.67567 | 4.05177 | 4.92497 | 2.42542 |
| | ***lnL*** | -8547.602397 | -2690.644795 | -14061.493950 | -8630.991740 | -3856.077856 | -2853.096431 | -13070.395831 | -3712.245453 | -5779.817707 | -6874.751966 |

**Table S19 continuation**. Summary statistics for branch-specific evolutionary models of single-copy orthologous subfamilies, analyzed using PAML v4 (Yang, 2007).

| Model | Statistics | OG41 | OG42 | OG43 | OG44 | OG45 | OG46 | OG47 | OG48 | OG49 | OG50 |
|---|---|---|---|---|---|---|---|---|---|---|---|
| **M7** | ***p*** | 2.75291 | 2.75463 | 13.81559 | 5.88250 | 0.17780 | 0.01270 | 0.16541 | 18.57656 | 0.02479 | 0.13611 |
| | ***q*** | 0.00500 | 0.00500 | 0.00500 | 0.00500 | 0.98920 | 0.30150 | 0.73963 | 0.00500 | 0.60206 | 1.34146 |
| | ***lnL*** | -1727.628142 | -2767.010532 | -3152.738153 | -3843.277680 | -8900.392501 | -3308.237776 | -3007.209124 | -1340.019158 | -6016.002738 | -2495.413217 |
| **M8** | ***p0*** | 0.81819 | 0.68841 | 0.74672 | 0.70673 | 0.99690 | 0.99170 | 0.95382 | 0.75717 | 0.98870 | 0.99999 |
| | ***p1*** | 0.18181 | 0.31159 | 0.25328 | 0.29327 | 0.00310 | 0.00830 | 0.04618 | 0.24283 | 0.01130 | 0.00001 |
| | ***p*** | 1.68251 | 0.01415 | 2.73622 | 1.08685 | 0.23483 | 0.01760 | 0.48285 | 99.00000 | 0.35122 | 0.13612 |
| | ***q*** | 0.00500 | 0.01909 | 0.00500 | 0.00500 | 1.37695 | 0.63598 | 3.34051 | 66.97474 | 20.92990 | 1.34165 |
| | ***ω*** | 7.16174 | 2.26902 | 8.59261 | 10.38204 | 2.77638 | 1.00000 | 1.43038 | 4.20947 | 1.00000 | 1.00000 |
| | ***lnL*** | -1676.231688 | -2741.965899 | -3069.716070 | -3713.752084 | -8891.840868 | -3303.653754 | -3004.453461 | -1315.860983 | -6008.361196 | -2495.413624 |

399

**Table S20.** Likelihood-ratio test results for site-specific evolutionary models of single-copy orthologous subfamilies, conducted using PAML v4 (Yang, 2007).

| Model | | OG01 | OG02 | OG03 | OG04 | OG05 | OG06 | OG07 | OG08 | OG09 | OG10 |
|---|---|---|---|---|---|---|---|---|---|---|---|
| M0 vs M3 | lnL | -2162.948435 | -12069.630996 | -5080.833417 | -3121.452981 | -5223.453997 | -10548.369935 | -9359.540371 | -763.252274 | -557.219077 | -6236.572796 |
| | lnL | -2112.487363 | -11800.164060 | -4977.163013 | -3028.144815 | -4963.837256 | -10328.578162 | -8946.887586 | -742.909191 | -553.225759 | -5937.036209 |
| | 2ΔL | 100.92 | 538.93 | 207.34 | 186.62 | 519.23 | 439.58 | 825.31 | 40.686 | 7.9866 | 599.07 |
| | *K* | 4 | 4 | 4 | 4 | 4 | 4 | 4 | 4 | 4 | 4 |
| | *alpha* | 0.05 | 0.05 | 0.05 | 0.05 | 0.05 | 0.05 | 0.05 | 0.05 | 0.05 | 0.05 |
| | *p* | 2.2e-16 | 2.2e-16 | 2.2e-16 | 2.2e-16 | 2.2e-16 | 2.2e-16 | 2.2e-16 | 3.122e-08 | 0.09207 | 2.2e-16 |
| | $\chi$ | 9.4877 | 9.4877 | 9.4877 | 9.4877 | 9.4877 | 9.4877 | 9.4877 | 9.4877 | 9.4877 | 9.4877 |
| M1a vs M2a | lnL | -2169.235915 | -11808.787819 | -4986.978991 | -3034.793061 | -4977.844425 | -10364.330361 | -9050.292984 | -748.377623 | -558.740901 | -6083.510086 |
| | lnL | -2120.742582 | -11808.787825 | -4977.452908 | -3033.125159 | -4963.865531 | -10364.330360 | -8947.296962 | -743.102722 | -553.225759 | -5937.927170 |
| | 2ΔL | 96.987 | -1.2e-05 | 19.052 | 3.3358 | 27.958 | 2,00E-06 | 205.99 | 10.55 | 11.03 | 291.17 |
| | *K* | 2 | 2 | 2 | 2 | 2 | 2 | 2 | 2 | 2 | 2 |
| | *alpha* | 0.05 | 0.05 | 0.05 | 0.05 | 0.05 | 0.05 | 0.05 | 0.05 | 0.05 | 0.05 |
| | *p* | 2.2e-16 | 1 | 7.292e-05 | 0.1886 | 8.493e-07 | 1 | 2.2e-16 | 0.005118 | 0.004025 | 2.2e-16 |
| | $\chi$ | 5.9915 | 5.9915 | 5.9915 | 5.9915 | 5.9915 | 5.9915 | 5.9915 | 5.9915 | 5.9915 | 5.9915 |
| M7 vs M8 | lnL | -2174.312191 | -11814.932584 | -4987.502003 | -3032.430812 | -4978.958372 | -10329.253345 | -9050.732544 | -748.936935 | -558.740903 | -6083.778050 |
| | lnL | -2120.742584 | -11800.268657 | -4977.528737 | -3029.343683 | -4963.886655 | -10328.535146 | -8947.455539 | -743.192940 | -553.225759 | -5938.151822 |
| | 2ΔL | 107.14 | 29.328 | 19.947 | 6.1743 | 30.143 | 1.4364 | 206.55 | 11.488 | 11.03 | 291.25 |
| | *K* | 2 | 2 | 2 | 2 | 2 | 2 | 2 | 2 | 2 | 2 |
| | *alpha* | 0.05 | 0.05 | 0.05 | 0.05 | 0.05 | 0.05 | 0.05 | 0.05 | 0.05 | 0.05 |
| | *p* | 2.2e-16 | 4.281e-07 | 4.663e-05 | 0.04563 | 2.847e-07 | 0.4876 | 2.2e-16 | 0.003202 | 0.004025 | 2.2e-16 |
| | $\chi$ | 5.9915 | 5.9915 | 5.9915 | 5.9915 | 5.9915 | 5.9915 | 5.9915 | 5.9915 | 5.9915 | 5.9915 |

**Table S20 continuation.** Likelihood-ratio test results for site-specific evolutionary models of single-copy orthologous subfamilies, conducted using PAML v4 (Yang, 2007).

| Model | | OG11 | OG12 | OG13 | OG14 | OG15 | OG16 | OG17 | OG18 | OG19 | OG20 |
|---|---|---|---|---|---|---|---|---|---|---|---|
| | lnL | -14004.587976 | -6347.668937 | -2282.294164 | -12827.962628 | -2885.791887 | -3922.364464 | -5887.196356 | -3779.892485 | -5887.196356 | -3779.892485 |
| | lnL | -13464.492160 | -6209.966418 | -2241.487657 | -12510.085125 | -2836.115816 | -3825.078218 | -5754.333055 | -3682.333934 | -5754.333055 | -3682.333934 |
| | 2ΔL | 1080.2 | 275.41 | 81.613 | 635.76 | 99.352 | 194.5 | 265.73 | 195.12 | 265.73 | 195.12 |
| M0 vs M3 | *K* | 4 | 4 | 4 | 4 | 4 | 4 | 4 | 4 | 4 | 4 |
| | *alpha* | 0.05 | 0.05 | 0.05 | 0.05 | 0.05 | 0.05 | 0.05 | 0.05 | 0.05 | 0.05 |
| | *p* | 2.2e-16 | 2.2e-16 | 2.2e-16 | 2.2e-16 | 2.2e-16 | 2.2e-16 | 2.2e-16 | 2.2e-16 | 2.2e-16 | 2.2e-16 |
| | *χ* | 9.4877 | 9.4877 | 9.4877 | 9.4877 | 9.4877 | 9.4877 | 9.4877 | 9.4877 | 9.4877 | 9.4877 |
| | lnL | -13663.361889 | -6211.419555 | -2297.410342 | -12519.847267 | -2836.443831 | -3834.113964 | -5765.926799 | -3693.393850 | -5765.926799 | -3693.393850 |
| | lnL | -13467.529096 | -6211.419555 | -2245.293250 | -12519.847267 | -2836.296475 | -3834.113964 | -5765.926798 | -3693.393850 | -5765.926798 | -3693.393850 |
| | 2ΔL | 391.67 | 0 | 104.23 | 0 | 0.29471 | 0 | 2,00E-06 | 0 | 2,00E-06 | 0 |
| M1a vs M2a | *K* | 2 | 2 | 2 | 2 | 2 | 2 | 2 | 2 | 2 | 2 |
| | *alpha* | 0.05 | 0.05 | 0.05 | 0.05 | 0.05 | 0.05 | 0.05 | 0.05 | 0.05 | 0.05 |
| | *p* | 2.2e-16 | 1 | 2.2e-16 | 1 | 0.863 | 1 | 1 | 1 | 1 | 1 |
| | *χ* | 5.9915 | 5.9915 | 5.9915 | 5.9915 | 5.9915 | 5.9915 | 5.9915 | 5.9915 | 5.9915 | 5.9915 |
| | lnL | -13665.327862 | -6218.707134 | -2303.037618 | -12521.830412 | -2846.434892 | -3829.332524 | -5755.809986 | -3688.029348 | -2406.888553 | -3688.029348 |
| | lnL | -13467.984638 | -6210.890798 | -2245.293250 | -12510.567346 | -2836.132958 | -3825.849050 | -5755.813062 | -3682.383972 | -2396.599605 | -3682.383972 |
| | 2ΔL | 394.69 | 15.633 | 115.49 | 22.526 | 20.604 | 6.9669 | -0.006152 | 11.291 | 20.578 | 11.291 |
| M7 vs M8 | *K* | 2 | 2 | 2 | 2 | 2 | 2 | 2 | 2 | 2 | 2 |
| | *alpha* | 0.05 | 0.05 | 0.05 | 0.05 | 0.05 | 0.05 | 0.05 | 0.05 | 0.05 | 0.05 |
| | *p* | 2.2e-16 | 0.0004031 | 2.2e-16 | 1.284e-05 | 3.357e-05 | 0.0307 | 1 | 0.003534 | 3.401e-05 | 0.003534 |
| | *χ* | 5.9915 | 5.9915 | 5.9915 | 5.9915 | 5.9915 | 5.9915 | 5.9915 | 5.9915 | 5.9915 | 5.9915 |

**Table S20 continuation.** Likelihood-ratio test results for site-specific evolutionary models of single-copy orthologous subfamilies, conducted using PAML v4 (Yang, 2007).

| Model | | OG21 | OG22 | OG23 | OG24 | OG25 | OG26 | OG27 | OG28 | OG29 | OG30 |
|---|---|---|---|---|---|---|---|---|---|---|---|
| M0 vs M3 | **lnL** | -7173.277608 | -1973.476058 | -2893.070677 | -1945.235038 | -7079.463353 | -1181.331501 | -4165.726469 | -11255.347165 | -3447.167177 | -4718.510529 |
| | **lnL** | -7012.525052 | -1942.808084 | -2746.114847 | -1918.986824 | -6782.359176 | -1169.588354 | -3980.362094 | -10767.412053 | -3347.711898 | -4625.850279 |
| | **2ΔL** | 321.51 | 61.336 | 293.91 | 52.496 | 594.21 | 23.486 | 370.73 | 975.87 | 198.91 | 185.32 |
| | ***K*** | 4 | 4 | 4 | 4 | 4 | 4 | 4 | 4 | 4 | 4 |
| | ***alpha*** | 0.05 | 0.05 | 0.05 | 0.05 | 0.05 | 0.05 | 0.05 | 0.05 | 0.05 | 0.05 |
| | ***p*** | 2.2e-16 | 1.519e-12 | 2.2e-16 | 1.086e-10 | 2.2e-16 | 0.0001012 | 2.2e-16 | 2.2e-16 | 2.2e-16 | 2.2e-16 |
| | ***χ*** | 9.4877 | 9.4877 | 9.4877 | 9.4877 | 9.4877 | 9.4877 | 9.4877 | 9.4877 | 9.4877 | 9.4877 |
| M1a vs M2a | **lnL** | -7228.052639 | -1943.028431 | -2749.813253 | -1919.354366 | -6804.813715 | -1169.588702 | -4009.024521 | -10796.382824 | -3363.304373 | -4647.758019 |
| | **lnL** | -7015.381481 | -1943.028431 | -2747.641931 | -1919.354367 | -6801.142675 | -1169.588702 | -3981.442258 | -10767.549294 | -3363.304385 | -4647.758019 |
| | **2ΔL** | 425.34 | 0 | 4.3426 | -2,00E-06 | 7.3421 | 0 | 55.165 | 57.667 | -2.4e-05 | 0 |
| | ***K*** | 2 | 2 | 2 | 2 | 2 | 2 | 2 | 2 | 2 | 2 |
| | ***alpha*** | 0.05 | 0.05 | 0.05 | 0.05 | 0.05 | 0.05 | 0.05 | 0.05 | 0.05 | 0.05 |
| | ***p*** | 2.2e-16 | 1 | 0.114 | 1 | 0.02545 | 1 | 1.05e-12 | 3.004e-13 | 1 | 1 |
| | ***χ*** | 5.9915 | 5.9915 | 5.9915 | 5.9915 | 5.9915 | 5.9915 | 5.9915 | 5.9915 | 5.9915 | 5.9915 |
| M7 vs M8 | **lnL** | -7260.652141 | -1945.359118 | -2750.956795 | -1925.351999 | -6793.559395 | -1171.791179 | -4011.459227 | -10797.123547 | -3351.699957 | -4631.216856 |
| | **lnL** | -7015.381484 | -1942.826664 | -2746.445973 | -1919.346836 | -6785.007090 | -1169.662614 | -3981.889340 | -10767.435440 | -3350.794888 | -4630.941590 |
| | **2ΔL** | 490.54 | 5.0649 | 9.0216 | 12.01 | 17.105 | 4.2571 | 59.14 | 59.376 | 1.8101 | 0.55053 |
| | ***K*** | 2 | 2 | 2 | 2 | 2 | 2 | 2 | 2 | 2 | 2 |
| | ***alpha*** | 0.05 | 0.05 | 0.05 | 0.05 | 0.05 | 0.05 | 0.05 | 0.05 | 0.05 | 0.05 |
| | ***p*** | 2.2e-16 | 0.07946 | 0.01099 | 0.002466 | 0.0001931 | 0.119 | 1.439e-13 | 1.278e-13 | 0.4045 | 0.7594 |

| | | | | | | | | | | | |
|---|---|---|---|---|---|---|---|---|---|---|---|
| | ***χ*** | 5.9915 | 5.9915 | 5.9915 | 5.9915 | 5.9915 | 5.9915 | 5.9915 | 5.9915 | 5.9915 | 5.9915 |

**Table S20 continuation.** Likelihood-ratio test results for site-specific evolutionary models of single-copy orthologous subfamilies, conducted using PAML v4 (Yang, 2007).

| Model | | OG31 | OG32 | OG33 | OG34 | OG35 | OG36 | OG37 | OG38 | OG39 | OG40 |
|---|---|---|---|---|---|---|---|---|---|---|---|
| **M0 vs M3** | **lnL** | -8768.449410 | -2868.612793 | -14621.285148 | -8894.045372 | -3929.296093 | -2874.910280 | -13312.710408 | -3928.586702 | -5930.825560 | -7009.639226 |
| | **lnL** | -8544.994586 | -2687.374379 | -14061.496172 | -8630.576510 | -3850.677314 | -2853.074629 | -13065.035639 | -3710.943836 | -5773.580811 | -6872.342203 |
| | **2ΔL** | 446.91 | 362.48 | 1119.6 | 526.94 | 157.24 | 43.671 | 495.35 | 435.29 | 314.49 | 274.59 |
| | ***K*** | 4 | 4 | 4 | 4 | 4 | 4 | 4 | 4 | 4 | 4 |
| | ***alpha*** | 0.05 | 0.05 | 0.05 | 0.05 | 0.05 | 0.05 | 0.05 | 0.05 | 0.05 | 0.05 |
| | ***p*** | 2.2e-16 | 2.2e-16 | 2.2e-16 | 2.2e-16 | 2.2e-16 | 7.508e-09 | 2.2e-16 | 2.2e-16 | 2.2e-16 | 2.2e-16 |
| | ***χ*** | 9.4877 | 9.4877 | 9.4877 | 9.4877 | 9.4877 | 9.4877 | 9.4877 | 9.4877 | 9.4877 | 9.4877 |
| **M1a vs M2a** | **lnL** | -8564.647612 | -2752.368582 | -14140.983283 | -8644.277513 | -3942.697472 | -2853.656918 | -13093.487688 | -3756.388973 | -5805.481261 | -6890.667418 |
| | **lnL** | -8545.108356 | -2690.644641 | -14061.772045 | -8635.257293 | -3856.077852 | -2853.656918 | -13093.487688 | -3712.184796 | -5805.481261 | -6890.667420 |
| | **2ΔL** | 39.079 | 123.45 | 158.42 | 18.04 | 173.24 | 0 | 0 | 88.408 | 0 | -4,00E-06 |
| | ***K*** | 2 | 2 | 2 | 2 | 2 | 2 | 2 | 2 | 2 | 2 |
| | ***alpha*** | 0.05 | 0.05 | 0.05 | 0.05 | 0.05 | 0.05 | 0.05 | 0.05 | 0.05 | 0.05 |
| | ***p*** | 3.267e-09 | 2.2e-16 | 2.2e-16 | 0.0001209 | 2.2e-16 | 1 | 1 | 2.2e-16 | 1 | 1 |
| | ***χ*** | 5.9915 | 5.9915 | 5.9915 | 5.9915 | 5.9915 | 5.9915 | 5.9915 | 5.9915 | 5.9915 | 5.9915 |
| **M7 vs M8** | **lnL** | -8574.059132 | -2753.230060 | -14141.789697 | -8646.652568 | -3947.301727 | -2855.538348 | -13079.471158 | -3756.977191 | -5794.915321 | -6875.501372 |
| | **lnL** | -8547.602397 | -2690.644795 | -14061.493950 | -8630.991740 | -3856.077856 | -2853.096431 | -13070.395831 | -3712.245453 | -5779.817707 | -6874.751966 |
| | **2ΔL** | 52.913 | 125.17 | 160.59 | 31.322 | 182.45 | 4.8838 | 18.151 | 89.463 | 30.195 | 1.4988 |
| | ***K*** | 2 | 2 | 2 | 2 | 2 | 2 | 2 | 2 | 2 | 2 |
| | ***alpha*** | 0.05 | 0.05 | 0.05 | 0.05 | 0.05 | 0.05 | 0.05 | 0.05 | 0.05 | 0.05 |
| | ***p*** | 3.236e-12 | 2.2e-16 | 2.2e-16 | 1.58e-07 | 2.2e-16 | 0.08699 | 0.0001145 | 2.2e-16 | 2.775e-07 | 0.4726 |

| | | | | | | | | | | | |
|---|---|---|---|---|---|---|---|---|---|---|---|
| | ***χ*** | 5.9915 | 5.9915 | 5.9915 | 5.9915 | 5.9915 | 5.9915 | 5.9915 | 5.9915 | 5.9915 | 5.9915 |

**Table S20 continuation.** Likelihood-ratio test results for site-specific evolutionary models of single-copy orthologous subfamilies, conducted using PAML v4 (Yang, 2007).

| Model | | OG41 | OG42 | OG43 | OG44 | OG45 | OG46 | OG47 | OG48 | OG49 | OG50 |
|---|---|---|---|---|---|---|---|---|---|---|---|
| **M0 vs M3** | **lnL** | -1708.733339 | -2773.327806 | -3118.301126 | -3785.486916 | -9144.155714 | -3347.179895 | -3053.244689 | -1345.203631 | -6073.114803 | -2544.769522 |
| | **lnL** | -1673.816166 | -2741.298992 | -3064.991199 | -3709.550828 | -8887.890899 | -3303.396341 | -3004.355660 | -1315.858938 | -6008.146911 | -2494.922373 |
| | **2ΔL** | 69.834 | 64.058 | 106.62 | 151.87 | 512.53 | 87.567 | 97.778 | 58.689 | 129.94 | 99.694 |
| | ***K*** | 4 | 4 | 4 | 4 | 4 | 4 | 4 | 4 | 4 | 4 |
| | ***alpha*** | 0.05 | 0.05 | 0.05 | 0.05 | 0.05 | 0.05 | 0.05 | 0.05 | 0.05 | 0.05 |
| | ***p*** | 2.46e-14 | 4.064e-13 | 2.2e-16 | 2.2e-16 | 2.2e-16 | 2.2e-16 | 2.2e-16 | 5.468e-12 | 2.2e-16 | 2.2e-16 |
| | ***χ*** | 9.4877 | 9.4877 | 9.4877 | 9.4877 | 9.4877 | 9.4877 | 9.4877 | 9.4877 | 9.4877 | 9.4877 |
| **M1a vs M2a** | **lnL** | -1719.991061 | -2753.946889 | -3152.415326 | -3843.277682 | -8919.315424 | -3303.994599 | -3004.787145 | -1335.653420 | -6008.522814 | -2505.373631 |
| | **lnL** | -1676.231683 | -2741.944017 | -3069.716063 | -3713.752060 | -8919.315424 | -3303.994599 | -3004.514350 | -1315.858938 | -6008.522814 | -2505.373631 |
| | **2ΔL** | 87.519 | 24.006 | 165.4 | 259.05 | 0 | 0 | 0.54559 | 39.589 | 0 | 0 |
| | ***K*** | 2 | 2 | 2 | 2 | 2 | 2 | 2 | 2 | 2 | 2 |
| | ***alpha*** | 0.05 | 0.05 | 0.05 | 0.05 | 0.05 | 0.05 | 0.05 | 0.05 | 0.05 | 0.05 |
| | ***p*** | 2.2e-16 | 6.127e-06 | 2.2e-16 | 2.2e-16 | 1 | 1 | 0.7612 | 2.531e-09 | 1 | 1 |
| | ***χ*** | 5.9915 | 5.9915 | 5.9915 | 5.9915 | 5.9915 | 5.9915 | 5.9915 | 5.9915 | 5.9915 | 5.9915 |
| **M7 vs M8** | **lnL** | -1727.628142 | -2767.010532 | -3152.738153 | -3843.277680 | -8900.392501 | -3308.237776 | -3007.209124 | -1340.019158 | -6016.002738 | -2495.413217 |
| | **lnL** | -1676.231688 | -2741.965899 | -3069.716070 | -3713.752084 | -8891.840868 | -3303.653754 | -3004.453461 | -1315.860983 | -6008.361196 | -2495.413624 |
| | **2ΔL** | 102.79 | 50.089 | 166.04 | 259.05 | 17.103 | 9.168 | 5.5113 | 48.316 | 15.283 | -0.000814 |
| | ***K*** | 2 | 2 | 2 | 2 | 2 | 2 | 2 | 2 | 2 | 2 |
| | ***alpha*** | 0.05 | 0.05 | 0.05 | 0.05 | 0.05 | 0.05 | 0.05 | 0.05 | 0.05 | 0.05 |
| | ***p*** | 2.2e-16 | 1.328e-11 | 2.2e-16 | 2.2e-16 | 0.0001932 | 0.01021 | 0.06357 | 3.223e-11 | 0.000480 | 1 |

| | | | | | | | | | | | |
|---|---|---|---|---|---|---|---|---|---|---|---|
| | | | | | | | | | | 1 | |
| | $\chi$ | 5.9915 | 5.9915 | 5.9915 | 5.9915 | 5.9915 | 5.9915 | 5.9915 | 5.9915 | 5.9915 | 5.9915 |

**Table S21**. Number of gustatory receptors (GR) and olfactory receptors (OR) subfamilies under positive selection (M8 model) and purifying (M0 and M1a, M7) based on the site model analysis PAML v4 (Yang, 2007).

| Genes | Number of genes | Sites | M0 vs M3 | M1a vs M2a | M7 vs M8 |
|---|---|---|---|---|---|
| **OG01** | 13 | 477 | M0 | M1a | M7 |
| **OG02** | 13 | 3060 | M0 | M2a | M7 |
| **OG03** | 13 | 1371 | M0 | M1a | M7 |
| **OG04** | 13 | 450 | M0 | M1a | M7 |
| **OG05** | 13 | 738 | M0 | M1a | M7 |
| **OG06** | 13 | 2679 | M0 | M2a | M7 |
| **OG07** | 13 | 1380 | M0 | M1a | M7 |
| **OG08** | 13 | 159 | M0 | M1a | M7 |
| **OG09** | 13 | 192 | M3 | M1a | M7 |
| **OG10** | 13 | 930 | M0 | M1a | M7 |
| **OG11** | 13 | 1617 | M0 | M1a | M7 |
| **OG12** | 13 | 1512 | M0 | - | M7 |
| **OG13** | 13 | 474 | M0 | M1a | M7 |
| **OG14** | 13 | 2178 | M0 | - | M7 |
| **OG15** | 13 | 813 | M0 | M2a | M7 |
| **OG16** | 13 | 945 | M0 | - | M7 |
| **OG17** | 13 | 1296 | M0 | M2a | M8 |
| **OG18** | 13 | 1005 | M0 | - | M7 |
| **OG19** | 13 | 678 | M0 | M2a | M7 |
| **OG20** | 13 | 939 | M0 | - | M7 |
| **OG21** | 13 | 1317 | M0 | M1a | M7 |
| **OG22** | 13 | 537 | M0 | - | M7 |
| **OG23** | 13 | 435 | M0 | M1a | M7 |
| **OG24** | 13 | 600 | M0 | M2a | M7 |
| **OG25** | 13 | 909 | M0 | M1a | M7 |
| **OG26** | 13 | 336 | M0 | - | M8 |
| **OG27** | 13 | 606 | M0 | M1a | M7 |
| **OG28** | 13 | 1575 | M0 | M1a | M7 |
| **OG29** | 13 | 477 | M0 | M2a | M8 |
| **OG30** | 13 | 702 | M0 | - | M8 |

**Table S20 continuation**. Single-copy orthologous groups (OG) suggested sites to evolve under positive and purifying selection based on the site model analysis.

| Genes | Number of genes | Sites | M0 vs M3 | M1a vs M2a | M7 vs M8 |
|---|---|---|---|---|---|
| **OG31** | 13 | 1527 | M0 | | M7 |
| **OG32** | 13 | 435 | M0 | M1a | M7 |
| **OG33** | 13 | 1854 | M0 | M1a | M7 |
| **OG34** | 13 | 1800 | M0 | M1a | M7 |
| **OG35** | 13 | 1029 | M0 | M1a | M7 |
| **OG36** | 13 | 879 | M0 | - | M8 |
| **OG37** | 13 | 1707 | M0 | - | M7 |
| **OG38** | 13 | 480 | M0 | M1a | M7 |
| **OG39** | 13 | 840 | M0 | - | M7 |
| **OG40** | 13 | 1425 | M0 | M2a | M8 |
| **OG41** | 13 | 1425 | M0 | M1a | M7 |
| **OG42** | 13 | 609 | M0 | M1a | M7 |
| **OG43** | 13 | 795 | M0 | M1a | M7 |
| **OG44** | 13 | 912 | M0 | M1a | M7 |
| **OG45** | 13 | 1434 | M0 | - | M7 |
| **OG46** | 13 | 948 | M0 | - | M7 |
| **OG47** | 13 | 696 | M0 | M2a | M8 |
| **OG48** | 13 | 297 | M0 | M1a | M7 |
| **OG49** | 13 | 1806 | M0 | - | M7 |
| **OG50** | 13 | 495 | M0 | - | M8 |

**Table S22**. Turkey test from the general linear model (P<0.001, ***$\chi$2***=192.8, df=82, Model 2) to test the distribution of the selection pressure among the different hemosensory genes and the single-copy orthologous gene groups (OG).

| contrast | | | Estimate | SE | df | z.ratio | p.value |
|---|---|---|---|---|---|---|---|
| OG | - | GR | -0.7977 | 0.146 | Inf | -5.478 | <.0001 |
| OG | - | OR | -0.7764 | 0.141 | Inf | -5.497 | <.0001 |
| GR | - | OR | 0.0213 | 0.118 | Inf | 0.181 | 0.9822 |

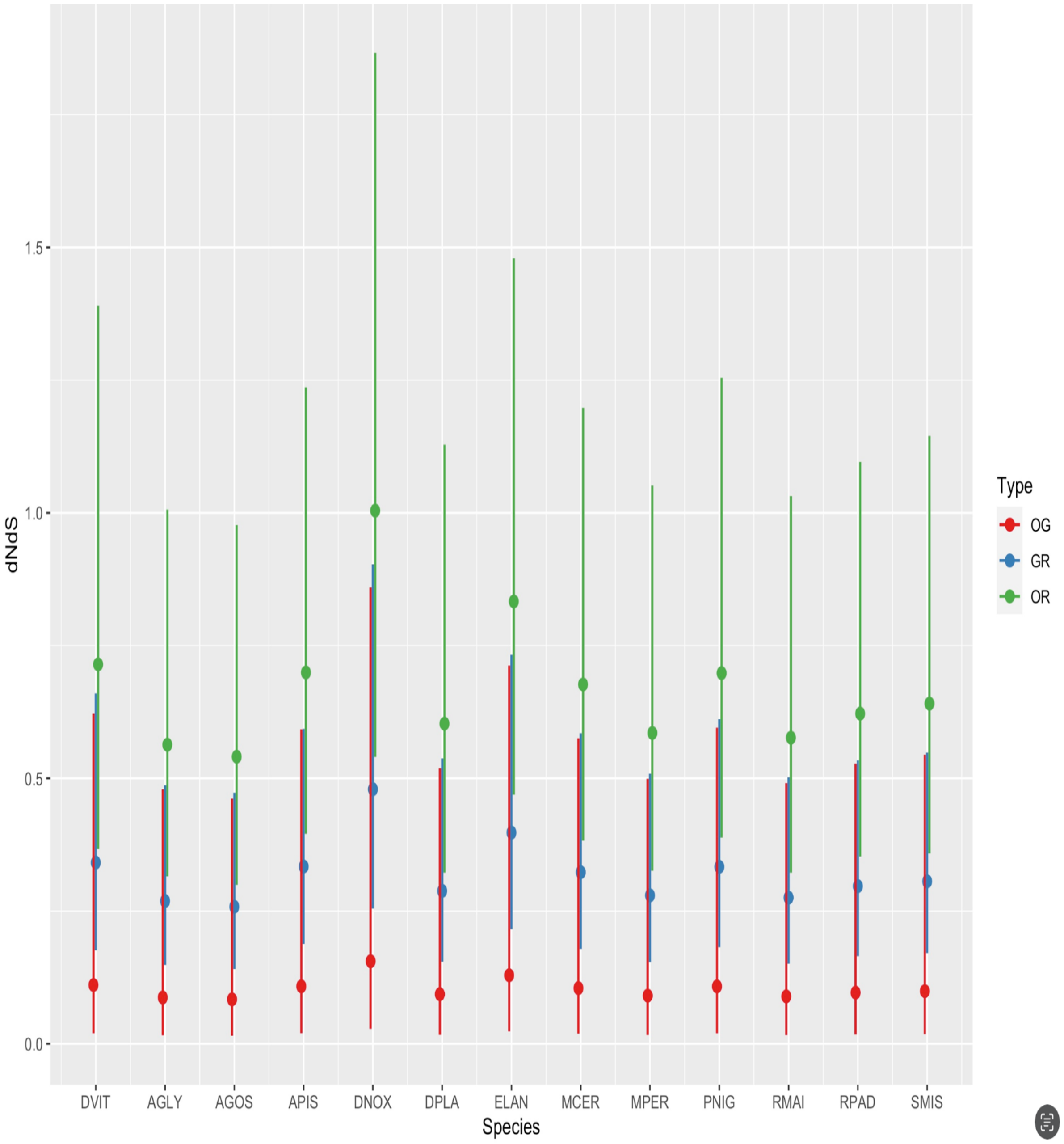

**Figure S10.** Distribution of dN/dS ratios for each olfactory receptor (OR) and gustatory receptor (GR) gene subfamily, as well as for single-copy orthologous gene families across different aphid species. For further details, refer to Table S22 (Model 2). AGLY = *Aphids glycines* clone BT1; AGOS = *Aphis gossypii*; APIS = *Acyrthosiphon pisum* clone LSR1; DNOX = *Diuraphis noxia*; DPLA = *Dysaphis plantaginea*; DVIT = *Daktulosphaira vitifoliae*; ELAN = *Eriosoma lanigerum*; MCER = *Myzus cerasi*; MPER = *Myzus persicae* clone O; PNIG = *Pentalonia nigronervosa*; RMAI = *Rhopalosiphum maidis*; RPAD = *Rhopalosiphum padi*; SMIS = *Sitobion miscanthi*.

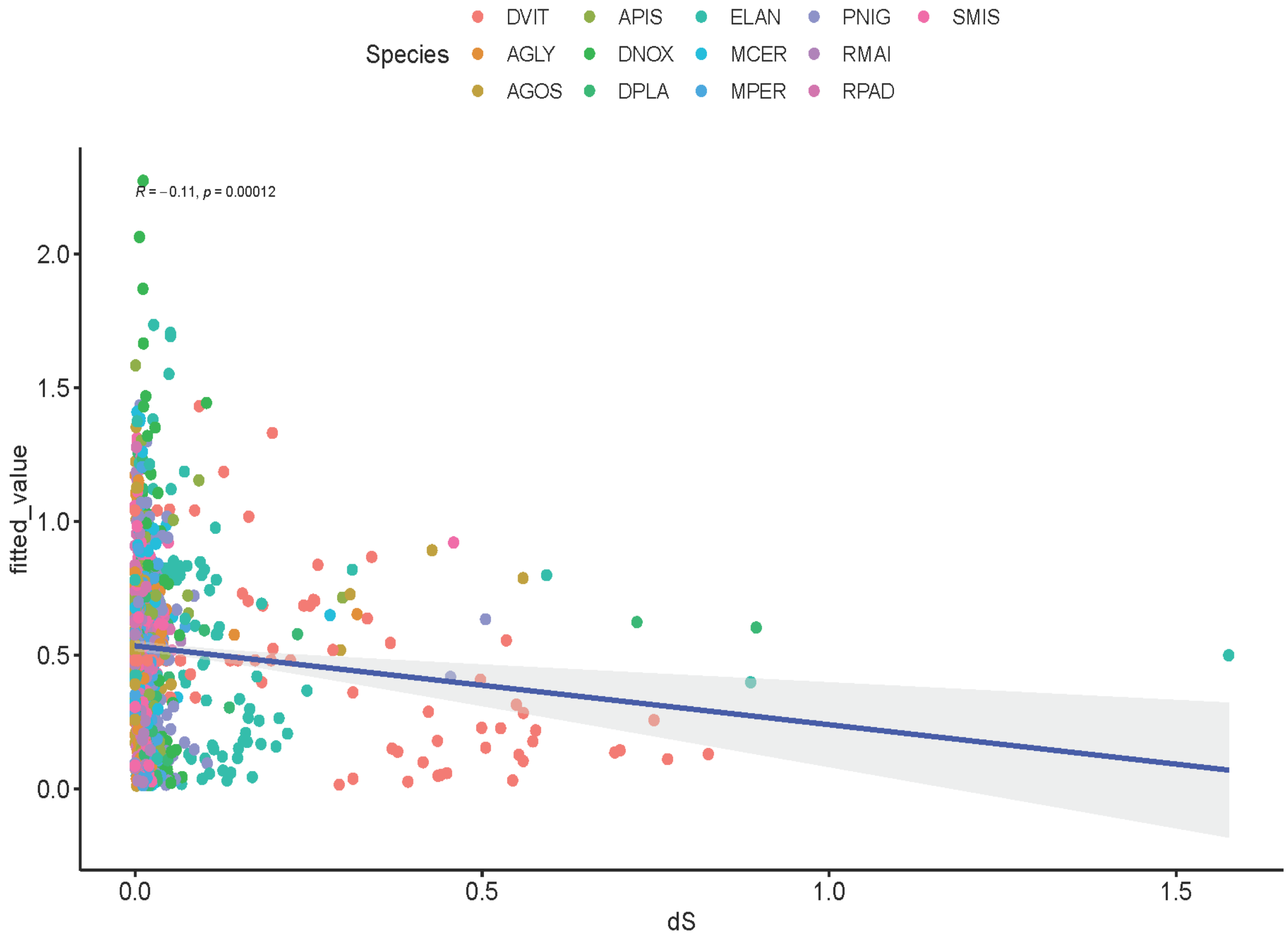

**Figure S11.** Correlation between the dN/dS ratio and synonymous divergence (dS) among gustatory receptor (GR) and olfactory receptor (OR) gene subfamilies, as well as single-copy orthologous gene families (OG) across 12 aphid species. . AGLY = *Aphids glycines* clone BT1; AGOS = *Aphis gossypii*; APIS = *Acyrthosiphon pisum* clone LSR1; DNOX = *Diuraphis noxia*; DPLA = *Dysaphis plantaginea*; DVIT = *Daktulosphaira vitifoliae*; ELAN = *Eriosoma lanigerum*; MCER = *Myzus cerasi*; MPER = *Myzus persicae* clone O; PNIG = *Pentalonia nigronervosa*; RMAI = *Rhopalosiphum maidis*; RPAD = *Rhopalosiphum padi*; SMIS = *Sitobion miscanthi*.

**Table S23. Key statistics from the genome assemblies utilized in the REPET pipeline for analyzing transposable elements across 12 aphid species and one aphid-like species.** The statistics include total assembly length, number of scaffolds, N50 values, and overall genome coverage, which collectively provide insights into the quality and completeness of the genomic data. AGLY = *Aphids glycines* clone BT1; AGOS = *Aphis gossypii*; APIS = *Acyrthosiphon pisum* clone LSR1; DNOX = *Diuraphis noxia*; DPLA = *Dysaphis plantaginea*; DVIT = *Daktulosphaira vitifoliae*; ELAN = *Eriosoma lanigerum*; MCER = *Myzus cerasi*; MPER = *Myzus persicae* clone O; PNIG = *Pentalonia nigronervosa*; RMAI = *Rhopalosiphum maidis*; RPAD = *Rhopalosiphum padi*; SMIS = *Sitobion miscanthi*.

| Species | DNOX | DPLA | DVIT |
|---|---|---|---|
| Sequences size (nb) | 5637 | 1457 | 10492 |
| Cum sequence lengths | 395073589 bp (395 Mb) | 416559878 bp (416 Mb) | 282671353 bp (282 Mb) |
| mean sequence length | 70085 bp | 285902 bp | 26941 bp |
| Var | 30456114370.67 | 180777758121.01 | 10953827094.39 |
| Sd | 174516.80 | 425179.68 | 104660.53 |
| cv | 2.49 | 1.49 | 3.88 |
| n | 5637 | 1457 | 10492 |
| min | 928 | 50051 | 141 |
| Q1 | 2242 | 80625 | 732 |
| median | 10659 | 146084 | 1077 |
| Q3 | 24771 | 296839 | 3389 |
| max | 2142037 | 5044119 | 2080308 |
| N50 | 281 | 184 | 236 |
| L50 | 397774nt | 518248nt | 341590nt |
| N75 | 661 | 504 | 557 |
| L75 | 169433nt | 218156nt | 129057 |
| N90 | 1362 | 900 | 1164 |
| L90 | 37041nt | 109054nt | 36802 |
| GC content | 86035086 (21.78%) | 125473212 (30.12%) | 75293698 (26.64%) |
| occurrences of A | 104953206 (26.57%) | 145476040 (34.92%) | 100560015 (35.57%) |
| occurrences of T | 104986705 (26.57%) | 145610626 (34.96%) | 100538992 (35.57%) |
| occurrences of G | 43027042 (10.89%) | 62727755 (15.06%) | 37634441 (13.31%) |
| occurrences of C | 43008044 (10.89%) | 62745457 (15.06%) | 37659257 (13.32%) |
| occurrences of N | 98535224 (24.94%) | 0 (0%) | 6278648 (2.22%) |
| occurrences of other | 563368 (0.14%) | | |

**Table S23.** Statistics from the genome assemblies utilized in the REPET pipeline for analyzing transposable elements in 12 aphid species and one aphid-like species. The statistics include total assembly length, number of scaffolds, N50 values, and overall genome coverage, which collectively provide insights into the quality and completeness of the genomic data. AGLY = *Aphids glycines* clone BT1; AGOS = *Aphis gossypii*; APIS = *Acyrthosiphon pisum* clone LSR1; DNOX = *Diuraphis noxia*; DPLA = *Dysaphis plantaginea*; DVIT = *Daktulosphaira vitifoliae*; ELAN = *Eriosoma lanigerum*; MCER = *Myzus cerasi*; MPER = *Myzus* *persicae* clone O; PNIG = *Pentalonia*

| Species | ELAN | MCER | MPER |
|---|---|---|---|
| **Sequences size (nb)** | 7146 | 49286 | 360 |
| **Cum sequence lengths** | 334868377 bp (334 Mb) | 405711039 bp (405 Mb) | 395142899 bp (395 Mb) |
| **mean sequence length** | 46860 bp | 8231 bp | 1097619 bp |
| **Var** | 2403432991129.35 | 208480886.95 | 79614281131038.03 |
| **Sd** | 1550300.94 | 14438.87 | 8922683.52 |
| **cv** | 33.08 | 1.75 | 8.13 |
| **n** | 7146 | 49286 | 360 |
| **min** | 1000 | 1001 | 5000 |
| **Q1** | 1380 | 1599 | 9881 |
| **median** | 2195 | 2957 | 17275 |
| **Q3** | 3792 | 7384 | 32495 |
| **max** | 71231741 | 265351 | 105178091 |
| **N50** | 3 | 4472 | 3 |
| **L50** | 62861676nt | 23273nt | 69480500nt |
| **N75** | 5 | 12033 | 4 |
| **L75** | 33516867nt | 7629nt | 62328371nt |
| **N90** | 6 | 25289 | 6 |
| **L90** | 29687234nt | 2853nt | 29865500 |
| **GC content** | 84889544 (25.35%) | 121142808 (29.86%) | 119241258 (30.18%) |
| **occurrences of A** | 121136053 (36.17%) | 142161864 (35.04%) | 137745117 (34.86%) |
| **occurrences of T** | 121083850 (36.16%) | 142206642 (35.05%) | 137769523 (34.87%) |
| **occurrences of G** | 42454008 (12.68%) | 60572748 (14.93%) | 59645355 (15.09%) |
| **occurrences of C** | 42435536 (12.67%) | 60570060 (14.93%) | 59595903 (15.08%) |
| **occurrences of N** | 7758930 (2.32%) | 199725 (0.05%) | 387001 (0.10%) |

*nigronervosa*; RMAI = *Rhopalosiphum maidis*; RPAD = *Rhopalosiphum padi*; SMIS = *Sitobion miscanthi*.

**Table S23 (continuation).** Key statistics from the genome assemblies utilized in the REPET pipeline for analyzing transposable elements across 12 aphid species and one aphid-like species. AGLY = *Aphids glycines* clone BT1; AGOS = *Aphis gossypii*; APIS = *Acyrthosiphon pisum* clone LSR1; DNOX = *Diuraphis noxia*; DPLA = *Dysaphis plantaginea*; DVIT = *Daktulosphaira vitifoliae*; ELAN = *Eriosoma lanigerum*; MCER = *Myzus cerasi*; MPER = *Myzus persicae* clone O; PNIG = *Pentalonia nigronervosa*; RMAI = *Rhopalosiphum maidis*; RPAD = *Rhopalosiphum padi*; SMIS = *Sitobion miscanthi*.

| Species | PNIG | RMAI | RPAD | SMIS |
|---|---|---|---|---|
| Sequences size (nb) | 18348 | 220 | 2172 | 653 |
| Cum sequence lengths | 375348459 bp (375 Mb) | 326023155 bp (326 Mb) | 321589008 bp (321 Mb) | 397852643 bp (397 Mb) |
| mean sequence length | 20457 bp | 1481923 bp | 148061 bp | 609268 bp |
| Var | 2203736131.51 | 119546129129072.09 | 104523010136.11 | 30160399665805.68 |
| Sd | 46943.97 | 10933715.25 | 323300.19 | 5491848.47 |
| cv | 2.29 | 7.38 | 2.18 | 9.01 |
| n | 18348 | 220 | 2172 | 653 |
| min | 1000 | 1096 | 1131 | 2682 |
| Q1 | 1522 | 7495 | 7982 | 16972 |
| median | 3033 | 20497 | 19633 | 29917 |
| Q3 | 11969 | 33477 | 123870 | 45044 |
| max | 631822 | 94224415 | 4088110 | 101470385 |
| N50 | 1038 | 2 | 140 | 4 |
| L50 | 103994nt | 93298903nt | 652723nt | 36263045nt |
| N75 | 2410 | 3 | 321 | 7 |
| L75 | 42723nt | 76887858nt | 298224nt | 32651544nt |
| N90 | 4986 | 4 | 583 | 9 |
| L90 | 9877nt | 56292413nt | 101141nt | 28943255nt |
| GC content | 107489442 (28.64%) | 90270308 (27.69%) | 89439794 (27.81%) | 119611961 (30.06%) |
| occurrences of A | 133781559 (35.64%) | 117853279 (36.15%) | 116094666 (36.10%) | 139175655 (34.98%) |
| occurrences of T | 133825498 (35.65%) | 117852668 (36.15%) | 116054548 (36.09%) | 139015827 (34.94%) |
| occurrences of G | 53753361 (14.32%) | 45115152 (13.84%) | 44701984 (13.90%) | 59809085 (15.03%) |
| occurrences of C | 53736081 (14.32%) | 45155156 (13.85%) | 44737810 (13.91%) | 59802876 (15.03%) |
| occurrences of N | 251960 (0.07%) | 46900 (0.01%) | 0 (0%) | 49200 (0.01%) |

**Table S23 (continuation).** Key statistics from the genome assemblies utilized in the REPET pipeline for analyzing transposable elements across 12 aphid species and one aphid-like species.

| Species | AGLY | AGOS | APIS |
|---|---|---|---|
| **Sequences size (nb)** | 1161 | 240 | 1604 |
| **Cum sequence lengths** | 2933887 bp (2 Mb) | 208307 bp | 1310621 bp (1 Mb) |
| **mean sequence length** | 2527 bp | 867 bp | 817 bp |
| **Var** | 6686596.36 | 310162.68 | 391656.36 |
| **Sd** | 2585.85 | 556.92 | 625.82 |
| **cv** | 1.02 | 0.64 | 0.77 |
| **n** | 1161 | 240 | 1604 |
| **min** | 347 | 326 | 331 |
| **Q1** | 718 | 535 | 542 |
| **median** | 1483 | 663 | 694 |
| **Q3** | 3445 | 979 | 938 |
| **max** | 16899 | 3996 | 18580 |
| **N50** | 215 | 68 | 512 |
| **L50** | 4159nt | 945nt | 858nt |
| **N75** | 441 | 137 | 963 |
| **L75** | 2545nt | 622nt | 611nt |
| **N90** | 712 | 193 | 1314 |
| **L90** | 9453nt | 500nt | 510nt |
| **GC content** | 803274 (27.38%) | 59362 (28.50%) | 390102 (29.76%) |
| **occurrences of A** | 1093377 (37.27%) | 74140 (35.59%) | 461661 (35.22%) |
| **occurrences of T** | 1037236 (35.35%) | 74805 (35.91%) | 458858 (35.01%) |
| **occurrences of G** | 403085 (13.74%) | 29798 (14.30%) | 194026 (14.80%) |
| **occurrences of C** | 400189 (13.64%) | 29564 (14.19%) | 196076 (14.96%) |
| **occurrences of N** | 0 (0%) | 0 (0%) | 0 (0%) |

**Table S23 (continuation).** Key statistics from the genome assemblies utilized in the REPET pipeline for analyzing transposable elements across 12 aphid species and one aphid-like species.

| Species | DNOX | DPLA | DVIT |
|---|---|---|---|
| **Sequences size (nb)** | 1262 | 2965 | 3535 |
| **Cum sequence lengths** | 893004 bp | 6321378 bp (6 Mb) | 4620002 bp (4 Mb) |
| **mean sequence length** | 707 bp | 2131 bp | 1306 bp |
| **Var** | 120586.28 | 5170311.22 | 1170010.58 |
| **Sd** | 347.26 | 2273.83 | 1081.67 |
| **cv** | 0.49 | 1.07 | 0.83 |
| **n** | 1262 | 2965 | 3535 |
| **min** | 341 | 327 | 329 |
| **Q1** | 511 | 640 | 566 |
| **median** | 610 | 1190 | 916 |
| **Q3** | 794 | 2746 | 1602 |
| **max** | 5409 | 18499 | 8462 |
| **N50** | 437 | 515 | 775 |
| **L50** | 709nt | 3869 | 12759nt |
| **N75** | 796 | 1108 | 1669 |
| **L75** | 551nt | 1886nt | 971nt |
| **N90** | 1056 | 1873 | 2609 |
| **L90** | 478nt | 824nt | 572nt |
| **GC content** | 281641 (31.54%) | 1946135 (30.79%) | 1417030 (30.67%) |
| **occurrences of A** | 311582 (34.89%) | 2247668 (35.56%) | 1634296 (35.37%) |
| **occurrences of T** | 299605 (33.55%) | 2127575 (33.66%) | 1568638 (33.95%) |
| **occurrences of G** | 138254 (15.48%) | 975599 (15.43%) | 708796 (15.34%) |
| **occurrences of C** | 143387 (16.06%) | 970536 (15.35%) | 708234 (15.33%) |
| **occurrences of N** | 176 (0.02%) | 0 (0%) | 38 (0.00%) |

**Table S23 (continuation).** Key statistics from the genome assemblies utilized in the REPET pipeline for analyzing transposable elements across 12 aphid species and one aphid-like species.

| Species | ELAN | MCER | MPER |
|---|---|---|---|
| **Sequences size (nb)** | 1157 | 1571 | 1806 |
| **Cum sequence lengths** | 2005004 bp (2 Mb) | 1172626 bp (1 Mb) | 3758845 bp (3 Mb) |
| **mean sequence length** | 1732 bp | 746 bp | 2081 bp |
| **Var** | 3739808.02 | 163345.43 | 5839490.59 |
| **Sd** | 1933.86 | 404.16 | 2416.50 |
| **cv** | 1.12 | 0.54 | 1.16 |
| **n** | 1157 | 1571 | 1806 |
| **min** | 343 | 332 | 325 |
| **Q1** | 629 | 515 | 626 |
| **median** | 1032 | 619 | 1038 |
| **Q3** | 1989 | 851 | 2580 |
| **max** | 16846 | 5297 | 20355 |
| **N50** | 199 | 515 | 281 |
| **L50** | 2612 | 751nt | 4092nt |
| **N75** | 477 | 966 | 632 |
| **L75** | 1273nt | 567nt | 1691nt |
| **N90** | 796 | 1302 | 1133 |
| **L90** | 701nt | 477nt | 745nt |
| **GC content** | 575011 (28.68%) | 349195 (29.78%) | 1179331 (31.37%) |
| **occurrences of A** | 735765 (36.70%) | 415464 (35.43%) | 1326250 (35.28%) |
| **occurrences of T** | 694168 (34.62%) | 407966 (34.79%) | 1253264 (33.34%) |
| **occurrences of G** | 287026 (14.32%) | 173180 (14.77%) | 586879 (15.61%) |
| **occurrences of C** | 287985 (14.36%) | 176015 (15.01%) | 592452 (15.76%) |
| **occurrences of N** | 60 (0.00%) | 1 (0.00%) | 0 (0%) |

**Table S23 (continuation).** Key statistics from the genome assemblies utilized in the REPET pipeline for analyzing transposable elements across 12 aphid species and one aphid-like species.

| Species | PNIG | RMAI | RPAD | SMIS |
|---|---|---|---|---|
| Sequences size (nb) | 920 | 830 | 1018 | 1516 |
| Cum sequence lengths | 1065104 bp (1 Mb) | 1597414 bp (1 Mb) | 1408656 bp (1 Mb) | 2458830 bp (2 Mb) |
| mean sequence length | 1157 bp | 1924 bp | 1383 bp | 1621 bp |
| Var | 786654.68 | 9038410.91 | 2499614.73 | 4049563.55 |
| Sd | 886.94 | 3006.40 | 1581.02 | 2012.35 |
| cv | 0.77 | 1.56 | 1.14 | 1.24 |
| n | 920 | 830 | 1018 | 1516 |
| min | 360 | 340 | 347 | 329 |
| Q1 | 566 | 629 | 576 | 596 |
| median | 809 | 851 | 764 | 877 |
| Q3 | 1400 | 1633 | 1451 | 1649 |
| max | 5831 | 23314 | 124843 | 17379 |
| N50 | 213 | 85 | 166 | 231 |
| L50 | 1467nt | 4206nt | 2173nt | 2500nt |
| N75 | 458 | 276 | 434 | 605 |
| L75 | 812nt | 1212nt | 842nt | 1128nt |
| N90 | 698 | 551 | 735 | 1047 |
| L90 | 564nt | 668nt | 591nt | 630nt |
| GC content | 343763 (32.28%) | 475307 (29.75%) | 422297 (29.98%) | 774646 (31.50%) |
| occurrences of A | 372134 (34.94%) | 567035 (35.50%) | 502243 (35.65%) | 859947 (34.97%) |
| occurrences of T | 349207 (32.79%) | 555072 (34.75%) | 484116 (34.37%) | 824237 (33.52%) |
| occurrences of G | 166441 (15.63%) | 233886 (14.64%) | 209464 (14.87%) | 380314 (15.47%) |
| occurrences of C | 177322 (16.65%) | 241421 (15.11%) | 212833 (15.11%) | 394332 (16.04%) |
| occurrences of N | 0 (0%) | 0 (0%) | 0 (0%) | 0 (0%) |

**Table S24.** Statistics of the transposable element consensus library from the REPET pipeline of the 12 aphid species and one aphid-like species.

| | AGLY | AGOS | APIS | DNOX | DPLA | DVIT | ELAN | MCER | MPER | PNIG | RMAI | RPAD | SMIS |
|---|---|---|---|---|---|---|---|---|---|---|---|---|---|
| **TEs in genome (Mb)** | 170 | 57 | 150 | 50 | 255 | 200 | 225 | 89 | 192 | 94 | 160 | 148 | 184 |
| **TE (%)** | 55.98 | 43.07 | 46.1 | 52.36 | 53.42 | 34.82 | 41.35 | 45.25 | 53.79 | 64.27 | 43.24 | 44.66 | 43.46 |
| **TE Class I (%)** | 39.31 | 18.98 | 21.4 | 26.46 | 12.94 | 20.31 | 15.79 | 14.83 | 20.78 | 17.95 | 22.23 | 14.98 | 19.38 |
| **TE Class II(%)** | 16.67 | 24.09 | 24.7 | 28.9 | 40.48 | 14.51 | 25.56 | 30.42 | 33.01 | 46.32 | 21.01 | 29.68 | 24.58 |
| **nb of sequences** | 1161 | 240 | 1604 | 1262 | 2965 | 3535 | 1157 | 1571 | 1806 | 920 | 830 | 1018 | 1516 |
| **nb of matched sequences** | 1160 | 240 | 1604 | 1262 | 2965 | 3535 | 1156 | 1571 | 1806 | 920 | 830 | 1018 | 1516 |
| **cumulative coverage bp** | 166110917 | 56228791 | 148644175 | 49251385 | 249297807 | 193123320 | 218938507 | 87955777 | 184846050 | 93472026 | 154699595 | 145805446 bp | 179825530 |
| **coverage percentage %** | 53.92 | 19.11 | 27.47 | 12.47 | 51.31 | 68.32 | 65.38 | 21.68 | 46.78 | 24.9 | 47.45 | 45.34% | 45.2 |
| **total nb of TE fragments** | 323127 | 182056 | 561840 | 211132 | 802447 | 574993 | 716784 | 355469 | 620004 | 303894 | 513091 | 498051 | 645412 |
| **total nb full-length fragments** | 4885 (1.51%) | 1895 (1.04%) | 18635 (3.32%) | 7956 (3.77%) | 17552 (2.19%) | 11822 (2.06%) | 5691 (0.79%) | 12342 (3.47%) | 10613 (1.71%) | 6628 (2.18%) | 5168 (1.01%) | 5864 (1.18%) | 10206 (1.58%) |
| **total nb of TE copies** | 294491 | 174024 | 513050 | 197322 | 739546 | 517136 | 649345 | 330577 | 567607 | 285699 | 476922 | 470171 | 588431 |
| **total nb full-length copies** | 5301 (1.80%) | 1987 (1.14%) | 19865 (3.87%) | 8336 (4.22%) | 18498 (2.50%) | 12616 (2.44%) | 6028 (0.93%) | 12995 (3.93%) | 11270 (1.99%) | 7023 (2.46%) | 5458 (1.14%) | 6146 (1.31%) | 10845 (1.84%) |
| **total nb full-length copies** | 5301 | 1987 | 19865 | 8336 | 18498 | 12616 | 6028 | 12995 | 11270 | 7023 | 5458 | 1001 | 10845 |
| **families with full-length fragments** | 1103 (95.00%) | 238 (99.17%) | 1586 (98.88%) | 1258 (99.68%) | 2915 (98.31%) | 3456 (97.77%) | 1128 (97.49%) | 1555 (98.98%) | 1762 (97.56%) | 914 (99.35%) | 809 (97.47%) | 1001 (98.33%) | 1486 (98.02%) |
| **with only one full-length fragment** | 344 | 10 | 169 | 174 | 456 | 1031 | 243 | 177 | 315 | 131 | 147 | 134 | 285 |
| **with only two full-length fragments** | 193 | 16 | 113 | 106 | 348 | 619 | 167 | 114 | 226 | 83 | 95 | 129 | 197 |
| **with only three full-length fragments** | 225 | 75 | 134 | 262 | 342 | 850 | 192 | 181 | 265 | 114 | 107 | 193 | 190 |
| **with more than three full-length fragments** | 341 | 137 | 1170 | 716 | 1769 | 956 | 526 | 1083 | 956 | 586 | 460 | 545 | 814 |

| | | | | | | | | | | | | | |
|---|---|---|---|---|---|---|---|---|---|---|---|---|---|
| **families with full-length copies** | 1116 (96.12 %) | 239 (99.58 %) | 1604 (100.00 %) | 1262 (100.00 %) | 2959 (99.80 %) | 3506 (99.18 %) | 1146 (99.05 %) | 1571 (100.00 %) | 1793 (99.28 %) | 919 (99.89 %) | 825 (99.40 %) | 1014 (99.61 %) | 1507 (99.41 %) |
| **with only one full-length copy** | 331 | 10 | 167 | 165 | 450 | 1005 | 240 | 181 | 309 | 122 | 142 | 132 | 273 |
| **with only two full-length copies** | 193 | 14 | 110 | 106 | 339 | 606 | 165 | 109 | 213 | 77 | 91 | 129 | 189 |
| **with only three full-length copies** | 237 | 77 | 130 | 250 | 337 | 838 | 185 | 164 | 264 | 112 | 113 | 182 | 192 |
| **with more than three full-length copies** | 355 | 138 | 1197 | 741 | 1833 | 1057 | 556 | 1117 | 1007 | 608 | 479 | 571 | 853 |
| **mean of median identity of all families** | 80.80 +- 7.54 | 80.23 +- 6.37 | 82.69 +- 6.36 | 80.56 +- 6.55 | 81.53 +- 7.27 | 83.91 +- 8.37 | 80.81 +- 7.18 | 81.72 +- 6.53 | 80.30 +- 6.58 | 79.87 +- 7.17 | 80.13 +- 6.46 | 79.47 +- 5.95 | 80.62 +- 6.07 |
| **mean of median length percentage of all families** | 23.53 +- 24.27 | 33.70 +- 24.35 | 29.72 +- 17.13 | 31.08 +- 16.72 | 21.95 +- 22.41 | 31.23 +- 26.30 | 22.51 +- 20.23 | 30.32 +- 16.46 | 22.39 +- 21.93 | 27.66 +- 18.37 | 25.03 +- 22.74 | 25.62 +- 20.06 | 23.21 +- 20.18 |

Table S25. Transposable elements (TEs) significant enrichment analysis in the Gustatory receptor (GR) and Olfactory receptor (OR) genes in a 2 kb and 10Kb window detected using Locus Overlap Analysis (LOLA)

| Species | Type | Enlargment size | qvalue | oddsRatio | support | rnkPV | rnkOR | rnkSup | maxRnk | meanRnk | b | c | d | TE name | size |
|---|---|---|---|---|---|---|---|---|---|---|---|---|---|---|---|
| AGLY | OR | 2kb | 1.20E-19 | 1398.89709 | 10 | 1 | 1 | 3 | 3 | 1.67 | 2 | 28 | 7976 | AGLY_TEdenovoGr-B-G3101-Map3 | 14 |
| DPLA | GR | 2kb | 3.85E-12 | 69.6311898 | 11 | 1 | 2 | 2 | 2 | 1.67 | 156 | 16 | 15816 | DPLA_TEdenovoGr-B-G77-Map20 | 293 |
| DPLA | OR | 2kb | 1.92E-13 | 30.6520993 | 21 | 1 | 10 | 1 | 10 | 4 | 1423 | 7 | 14548 | DPLA_TEdenovoGr-B-G4075-Map8_reversed | 3170 |
| DPLA | OR | 2kb | 6.66E-07 | 215.130799 | 5 | 2 | 2 | 10 | 10 | 4.67 | 16 | 23 | 15955 | DPLA_TEdenovoGr-B-G4329-Map4 | 35 |
| DPLA | OR | 2kb | 0.00393826 | 32.1842665 | 5 | 3 | 8 | 10 | 10 | 7 | 107 | 23 | 15864 | DPLA_TEdenovoGr-B-G161-Map14 | 191 |
| DPLA | OR | 2kb | 0.00951356 | 18.1687142 | 6 | 4 | 27 | 6 | 27 | 12.3 | 236 | 22 | 15735 | DPLA_TEdenovoGr-B-G149-Map20_reversed | 398 |
| DVIT | OR | 2kb | 1.86E-37 | 68.5243359 | 33 | 1 | 4 | 1 | 4 | 2 | 191 | 28 | 11135 | DVIT_TEdenovoGr-B-G6314-Map6 | 395 |
| DVIT | OR | 2kb | 7.89E-31 | 539.53315 | 17 | 2 | 2 | 2 | 2 | 2 | 8 | 44 | 11318 | DVIT_TEdenovoGr-B-G9099-Map3 | 36 |
| DVIT | OR | 2kb | 5.28E-05 | 47.2953315 | 6 | 3 | 7 | 22 | 22 | 10.7 | 26 | 55 | 11300 | DVIT_TEdenovoGr-B-G7829-Map4 | 43 |
| DVIT | OR | 2kb | 0.00051714 | Inf | 3 | 4 | 1 | 63 | 63 | 22.7 | 0 | 58 | 11326 | DVIT_TEdenovoGr-B-G678-Map3 | 5 |
| MCER | OR | 2kb | 0.00209786 | 61.7360344 | 4 | 1 | 1 | 11 | 11 | 4.33 | 18 | 59 | 16435 | MCER_TEdenovoGr-B-G1335-Map3 | 85 |

| | | | | | | | | | | | | | | | |
|---|---|---|---|---|---|---|---|---|---|---|---|---|---|---|---|
| RPAD | OR | 2kb | 2.31E-28 | 2754.73751 | 14 | 1 | 1 | 6 | 6 | 2.67 | 1 | 43 | 9380 | RPAD_TEdenovoGr-B-G1773-Map4 | 75 |
| RPAD | OR | 2kb | 0.00595325 | 15.6290395 | 6 | 2 | 3 | 29 | 29 | 11.3 | 70 | 51 | 9311 | RPAD_TEdenovoGr-B-G1099-Map3 | 150 |
| AGLY | OR | 10kb | 1.00E-10 | 481.965749 | 7 | 1 | 1 | 18 | 18 | 6.67 | 2 | 22 | 3146 | AGLY_TEdenovoGr-B-G3101-Map3 | 14 |
| DPLA | GR | 10kb | 2.25E-05 | 18.5275279 | 10 | 1 | 8 | 5 | 8 | 4.67 | 191 | 14 | 4964 | DPLA_TEdenovoGr-B-G77-Map20 | 293 |
| DPLA | OR | 10kb | 8.69E-06 | 18.9007361 | 19 | 1 | 13 | 1 | 13 | 5 | 1294 | 3 | 3863 | DPLA_TEdenovoGr-B-G4075-Map8_reversed | 3170 |
| DPLA | OR | 10kb | 0.00028014 | 62.409926 | 5 | 2 | 5 | 17 | 17 | 8 | 24 | 17 | 5133 | DPLA_TEdenovoGr-B-G4329-Map4 | 35 |
| DVIT | OR | 10kb | 2.34E-25 | 471.171951 | 16 | 1 | 1 | 9 | 9 | 3.67 | 4 | 36 | 4283 | DVIT_TEdenovoGr-B-G9099-Map3 | 36 |
| DVIT | OR | 10kb | 8.56E-17 | 19.0073531 | 27 | 2 | 6 | 1 | 6 | 3 | 230 | 25 | 4057 | DVIT_TEdenovoGr-B-G6314-Map6 | 395 |
| DVIT | OR | 10kb | 0.00782038 | 70.6851902 | 4 | 3 | 3 | 93 | 93 | 33 | 5 | 48 | 4282 | DVIT_TEdenovoGr-B-G4429-Map3 | 12 |
| DVIT | OR | 10kb | 0.01291861 | 58.9928594 | 4 | 4 | 4 | 93 | 93 | 33.7 | 6 | 48 | 4281 | DVIT_TEdenovoGr-B-G3298-Map9_reversed | 13 |
| DVIT | OR | 10kb | 0.02277339 | 258.20841 | 3 | 5 | 2 | 145 | 145 | 50.7 | 1 | 49 | 4286 | DVIT_TEdenovoGr-B-G678-Map3 | 5 |
| MCER | OR | 10kb | 0.00295944 | 29.4612514 | 5 | 1 | 1 | 13 | 13 | 5 | 35 | 57 | 11776 | MCER_TEdenovoGr-B-G1335-Map3 | 85 |
| RPAD | OR | 10kb | 4.72E-13 | 307.097008 | 9 | 1 | 1 | 35 | 35 | 12.3 | 3 | 31 | 3257 | RPAD_TEdenovoGr-B-G1773-Map4 | 75 |

505

**Table S26.** Relationships between enriched transposable elements (TEs) and associated Gustatory receptor (GR) and Olfactory receptor (OR) genes estimated using TEgriP.

See Table S26 Excel file

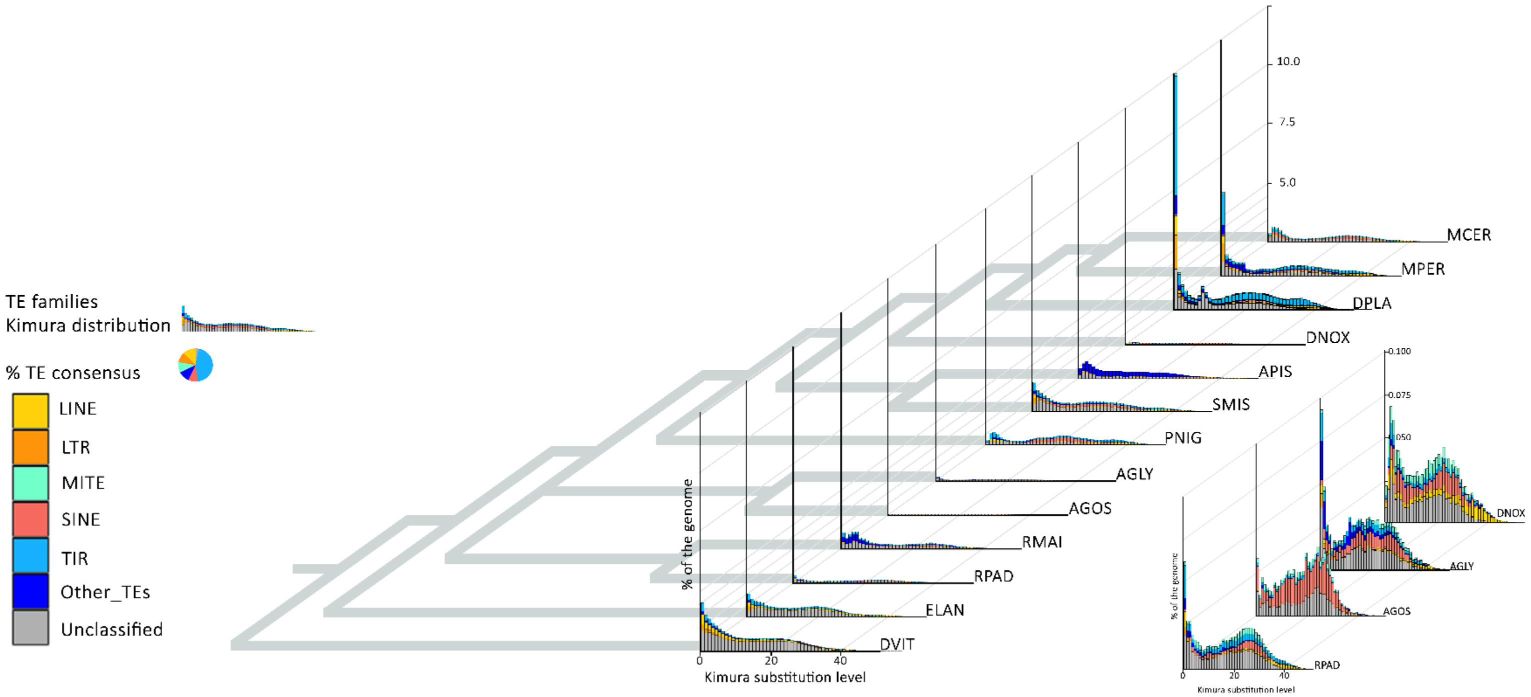

**Figure S12.** Transposable elements (TEs) content among twelve aphid species and one aphid-like species represented with the kimura distribution. AGLY = *Aphids glycines* clone BT1; AGOS = Aphis gossypii; APIS = *Acyrthosiphon pisum* clone LSR1; DNOX = *Diuraphis noxia*; DPLA = *Dysaphis plantaginea*; DVIT = *Daktulosphaira vitifoliae*; ELAN = *Eriosoma lanigerum*; MCER = *Myzus cerasi*; MPER = *Myzus persicae* clone O; PNIG = *Pentalonia nigronervosa*; RMAI = *Rhopalosiphum maidis*; RPAD = *Rhopalosiphum padi*; SMIS = *Sitobion miscanthi*.

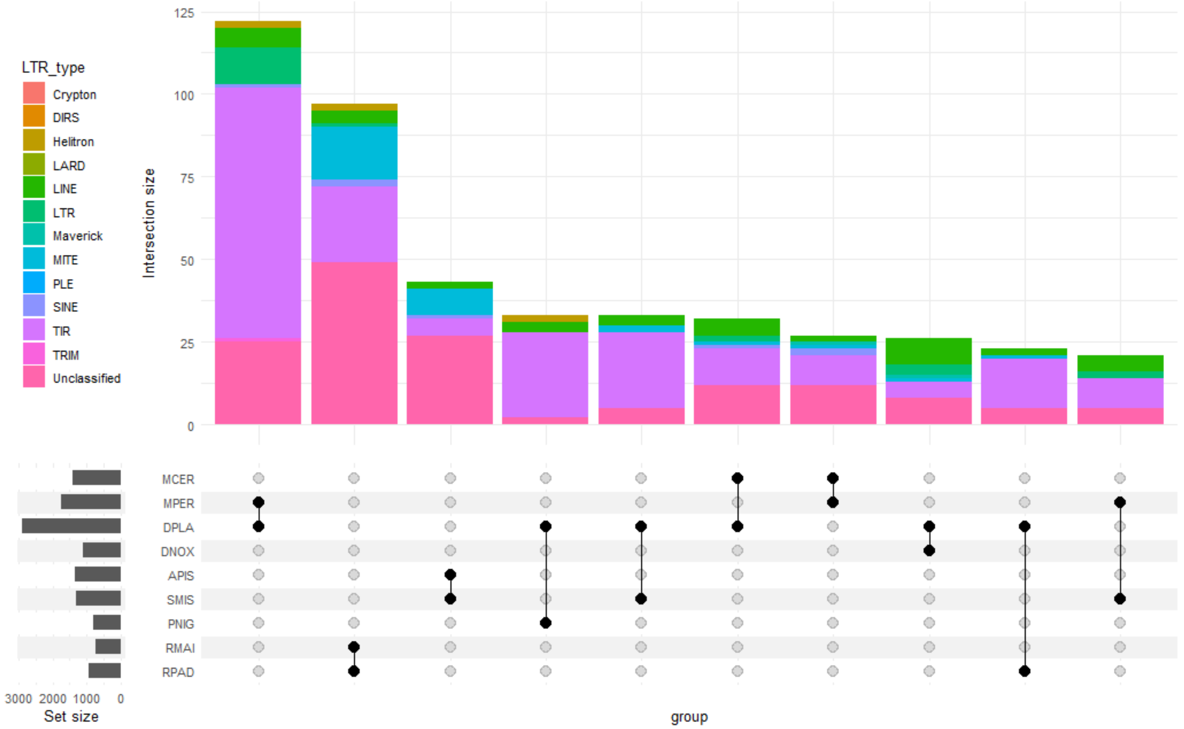

**Figure S13.** Transposable element-copy identity number among the 12 aphid species using an upset plot (UpSetR R package, Conway et al., 2017).

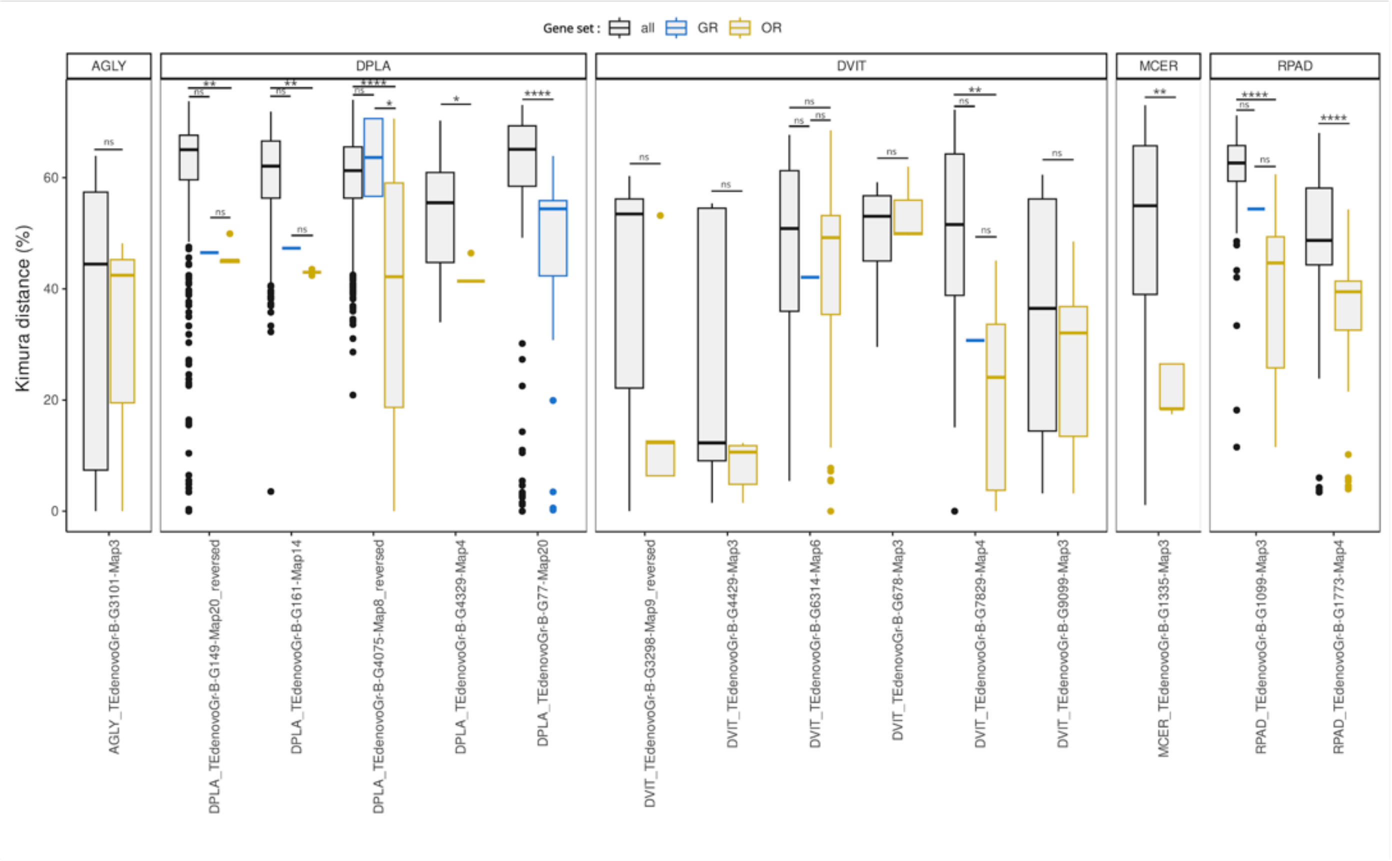

**Figure S14**. Distribution of Kimura distances between TE consensus sequence and each TE copies associated or not to chemosensory genes, for *A. glycines* (AGLY), *D. plantaginea* (DPLA), *D. vitifoliae* (DVIT), *M.cerasi* (MCER) and *R. padi* (RPAD). Statistical differences between the distribution of Kimura distances of all TE copies (in black), TE copies associated with gustatory receptor genes (in blue), and TE copies associated with olfactory receptor genes (in yellow) were computed using Kruskal-Wallis tests. ns: non-significant differences (p-value > 0.05) ; * : p-value < 0.05 ; ** : p-value < 0.01 ; *** : p-value < 0.005 ; **** : p-value < 0.001.

**Table S27. Available *de novo* consensus library done with REPET for the 12 aphid and one aphid-like species genomes on RepetDB (Amselem et al. 2019).** To download the fasta sequences, click on “Export,” then select “fasta sequences” format and click on “Download file”.

| Species | Link to de novo consensus libraries in RepetDB |
|---|---|
| *Aphis glycines (Clone BT1)* | https://urgi.versailles.inrae.fr/repetdb/begin.do#search?taxonGroup=307491 |
| *Aphis gossypii* | https://urgi.versailles.inrae.fr/repetdb/begin.do#search?taxonGroup=80765 |
| *Acyrthosiphon pisum (clone LSR1)* | https://urgi.versailles.inrae.fr/repetdb/begin.do#search?taxonGroup=7029 |
| *Diuraphis noxia* | https://urgi.versailles.inrae.fr/repetdb/begin.do#search?taxonGroup=143948 |
| *Dysaphis plantaginea* | https://urgi.versailles.inrae.fr/repetdb/begin.do#search?taxonGroup=214836 |
| *Daktulosphaira vitifoliae* | https://urgi.versailles.inrae.fr/repetdb/begin.do#search?taxonGroup=58002 |
| *Eriosoma lanigerum* | https://urgi.versailles.inrae.fr/repetdb/begin.do#search?taxonGroup=133082 |
| *Myzus cerasi* | https://urgi.versailles.inrae.fr/repetdb/begin.do#search?taxonGroup=93721 |
| *Myzus persicae (clone 0)* | https://urgi.versailles.inrae.fr/repetdb/begin.do#search?taxonGroup=13164 |
| *Pentalonia nigronervosa* | https://urgi.versailles.inrae.fr/repetdb/begin.do#search?taxonGroup=693967 |
| *Rhopalosiphum maidis* | https://urgi.versailles.inrae.fr/repetdb/begin.do#search?taxonGroup=43146 |
| *Rhopalosiphum padi* | https://urgi.versailles.inrae.fr/repetdb/begin.do#search?taxonGroup=40932 |
| *Sitobion miscanthi* | https://urgi.versailles.inrae.fr/repetdb/begin.do#search?taxonGroup=44668 |
| *Aphis glycines (Clone BT1)* | https://urgi.versailles.inrae.fr/repetdb/begin.do#search?taxonGroup=307491 |
