## Supplementary Material for "Comprehensive annotation of olfactory and gustatory receptor genes and transposable elements revealed their evolutionary dynamics in aphids"

**Text S1. Materials, methods, and results for *Dysaphis plantaginea* genome assembly, gene, and TE annotation.**

**Sample material, DNA extraction, and RNA extraction**

We used a clone (16-0042-0001) from a colony of *Dysaphis plantaginea* Passerini that was collected on Malus domestica Borkh in 2016 in Torchefelon (France). We reared it on Plantago lanceolata L. under controlled conditions (16h light, 8h dark at 20°C and 75% humidity) at the Agence Nationale de Sécurité Sanitaire de l'Alimentation, de l'Environnement et du Travail (ANSES, Lyon, France). For sequencing, we extracted high molecular weight DNA from clone 16_042 at the Centre National de Ressources Génomiques Végétales (CNRGV Toulouse, France).

Total RNA was extracted from aphid samples at various life cycle stages (Table S1) using the RNeasy Mini Kit (QIAGEN, Cat No./ID: 74104) following the manufacturer's protocol. The extracted RNA was quantified and assessed for integrity using a Bioanalyzer (Agilent Technologies). RNA sequencing was performed using paired-end 2 × 150 bp reads on the Illumina NovaSeq S4 platform at the Centre National de Ressources Génomiques Végétales (CNRGV Toulouse, France).

**Table S1. Details of the six developmental stages of the clone of *Dysaphis plantaginea* clone 16-0042-0001 used for gene annotation of the genome assembled in this study.**

| **ID** | **stage*** | **host** |
|---|---|---|
| n2_10J | Larviparous female - stage L1 - 10 days | apple |
| n2_10A | Adult female | apple |
| n2_funda | Fundatrigenae | apple |
| n2_males | winged male | flight from plantago to apple |
| n2_ovipares | Ovipareae | plantago |
| n2_gyno | winged female gynoparae | apple |

**Sequencing and assembly**

We sequenced it using Chromium 10X Genomics technology (Genopole Toulouse, Toulouse, France). We assembled the 10X reads using Supernova™ (Weisenfeld et al. 2017). We defined two assemblies as 56X and 100X due to the number of fragments involved in the process. We built the scaffolds using the ARCS (Assembly Round-up by Chromium Scaffolding) algorithm (Yeo et al. 2018). We assessed the quality of the *D. plantaginea* assembly version 1 using Benchmarking Universal Single-Copy Orthologs BUSCO (Waterhouse et al. 2018). We confronted the two assemblies (56X and 100X) with BWA (Li et al. 2009) and Samtools (Li et al. 2009) (Table S2). We utilized this first version of the *D. plantaginea* genome assembly to extract the complete *Buchnera aphidicola* Munson and mitochondrial genomes. We reconstructed the *B. aphidicola* genome with two methods. First, we identified the scaffolds by blasting the assemblages against the bacterium *B. aphidicola* of *Myzus persicae* Sulzer reference library (Jiang et al. 2013). Second, we filled the gaps using the raw data, and the scaffolds aligned on the *B. aphidicola* genome of *M. persicae*.

We improved the *D. plantaginea* genome assembly v1 using long reads obtained from Pacific Biosciences (PacBio) and Oxford Nanopore Technologies (ONT). We used wtdbg2 v2.5 (Ruan and Li 2020) on long reads and ntEdit (Warren et al. 2019) on the 10X Illumina sequences to search for inconsistencies (e.g., local misassemblies). We constructed scaffolds with Tigmint (Jackman et al. 2018) and ARCS (Yeo et al. 2018) using the 10X data. Finally, we reassembled the genome using FLYE (Kolmogorov et al. 2019), filtering out the reads from *B. aphidicola*. Afterward, we applied the software and procedure of the second version of the *D. plantaginea* genome assembly to obtain version 3 of the genome (Table S2). We constructed the optical mapping using MapSolver (OpGen®) to understand the genomic structure and structural variation.

### Gene and transposable (TE) annotation

We performed gene annotation using MAKER2 (Holt and Yandell 2011) and TE annotation with REPET (see details in the Materials and Methods (Flutre et al. 2011)). We detected and removed the mitochondrial genome sequence based on its similarity with *M. persicae* during genome assembly. The initial mitochondrial scaffold was 18.8 Kbp in length. We performed gene prediction on this scaffold with MITOS (Bernt et al. 2013) and ARWEN v1.2 (Laslett and Canbäck 2008) and RNA extracted above.

### ***Dysaphis plantaginea* final assembly (version 3) statistics**

The *D. plantaginea* genome v3 represented the twelfth released genome in the Aphidinae subfamily on the AphidBase-BIPAA platform. The *D. plantaginea* genome assembly version 3 had a total size of 485.98 Mb. The assembly statistics of different versions of the *D. plantaginea* are presented in Tables S2 and S3. The genome assembly comprised 2,366 scaffolds with a mean scaffold size of 205,401 bp and an N50 value of 17,731,133 bp. BUSCO analysis based on hemipteran conserved genes indicated the presence of 99.4% complete genes. We detected a genomic GC content of 30.14% and annotated 53,848 genes and 18,665 proteins.
